# The neuroscience of highly stable, positive, and refined states of consciousness during jhana-type advanced concentration absorption meditation (ACAM-J)

**DOI:** 10.1101/2025.11.12.688050

**Authors:** Winson F.Z. Yang, Ruby Potash, Grace Mackin, Isidora Beslic, Marta Bianciardi, Terje Sparby, Matthew D. Sacchet

## Abstract

Humans can experience a rich array of conscious experience, including highly stable, positive, and refined states of non-ordinary perception that can be elicited through advanced meditation. Here, we present the first group-level, ultra–high-field (7T) fMRI study of jhana advanced concentration absorption meditation (ACAM-J). We combined local (regional homogeneity), mesoscale (connectivity gradients), and global (geometric eigenmodes) human brain mapping with diverse and detailed phenomenology, meditative traits, behaviorally assessed cognitive functions, and extensive publicly available psychobehavioral affinity maps. Across eight successive states of ACAM-J, we observed reproducible neural trajectories marked by anterior-to-posterior reorganization, flattening of cortical hierarchies, nonlinear reconfiguration of global harmonics, and tight coupling between brain metrics and equanimity, attentional stability, and behavior. Neurosynth decoding further associate ACAM-J with reduced suffering-related processes and more with attentional monitoring. These findings suggest ACAM-J is a distinct, structured mode of awareness that persists despite radically reduced narrative thought and sensory content. Our findings also inform boundary conditions for current models of consciousness. More broadly, this research indicates that advanced meditation is a powerful framework for understanding psychological transformation, and for potential new opportunities for supporting human well-being and flourishing.

## Main Text

Human consciousness is not confined to ordinary perception and thought. With training, it can reorganize into highly phenomenologically structured, non-ordinary states of consciousness (*1, 2*). According to contemplative traditions, advanced meditation practices progress toward meditative endpoints (*2*) characterized by what are thought to be irreversible shifts in perception, identity, and cognition (*3, 4*), known variously as “enlightenment” (*5, 6*), “awakening” (*7*), or “salvation” (*8*). Rather than being driven by narrative processing and reactive affect and cognition, these states of consciousness cultivate a radically different organization of awareness rooted in equanimity, non-judgment, and openness (*9–13*). These states of consciousness are sometimes reported to be phenomenologically vivid despite the absence of ordinary narrative thought. These experiences remain almost entirely unstudied in modern science (*14*), despite their significance in contemplative traditions and their potential to inform both the upper limits of human potential and the mechanisms of psychological suffering (*15, 16*).

Unlike the heterogeneous effects of ordinary meditation, these advanced meditation states are structured and reproducible, with practitioners able to reliably enter and describe them (*2*). This makes advanced meditation states a unique natural model for testing deeper theoretical challenges of consciousness, including how the brain sustains phenomenally vivid awareness near absence of narrative thought, affect, and sensory content. Importantly, these states may strongly counteract the patterns of rumination, hyper-reactivity, compulsive craving that characterize psychiatric disorders, pointing to new frameworks for conceptualizing mental health, including well-being and flourishing.

Among the most structured and reproducible maps of advanced meditation described in contemplative traditions is *jhāna*, a sequence of absorption states preserved in Buddhism for over two millennia (*17*). Here, we refer to this practice scientifically as jhana advanced concentration absorption meditation (ACAM-J), thereby distinguishing the neurocognitive features under investigation scientifically from doctrinal interpretations. This thereby anchors in phenomenology and avoids, in some cases, thousands of years of controversies for example, related to defining jhana. ACAM-J are eight successive advanced meditation states that emerge through increasingly subtle absorption and attentional refinement (*17*). Early ACAM-J are characterized by strong positive affective qualities of joy, bliss, and energized yet stable attention. In contrast, later ACAM-J are characterized by a profound sense of formlessness, marked by the reduced sensory content and self-referential thought. Awareness becomes expansive, boundless, and subtle, often described as spacious emptiness or a state beyond ordinary perception. Each ACAM-J is phenomenologically distinct and reliably reportable by experienced practitioners (*18*). This makes ACAM-J a powerful and ideal framework for systematic investigation of non-ordinary consciousness. Despite this potential, ACAM-J remains unstudied in modern neuroscience, with prior studies constrained to limited methodology and single-case observations (*14, 17, 19–27*).

Contemporary models of consciousness such as the Global Neuronal Workspace Theory (GNWT) (*28*) and Integrated Information Theory (IIT) (*29*) emphasize sensory processing, task engagement, and narrative cognition, hypothesizing that conscious experience depends on information accessibility or integration to support goal-directed behavior. These models account well for ordinary experience but struggle to explain content-minimal yet phenomenologically vivid states like ACAM-J (*30, 31*). Recent proposals such as the Minimal Phenomenal Experience (MPE) (*32*) address this by characterizing sparse but present conscious states but remain untested against structured meditative trajectories. ACAM-J thus offers a unique test case for evaluating and extending theories of consciousness.

Here, we present the first group-level, multiscale neurophenomenological study of ACAM-J, combining ultra-high-field 7 Tesla fMRI we integrated local (regional homogeneity), mesoscale (connectivity gradients), and global (geometric eigenmodes) measures with detailed phenomenology, meditative traits, and behavioral assessments. Our findings reveal a reproducible neural trajectory of absorption marked by anterior-to-posterior reorganization, flattening of cortical hierarchies, nonlinear trends in global harmonics, and tight coupling to subjective equanimity and cognition. These results establish ACAM-J as a distinct, structured mode of awareness with a unique neural signature that challenges current models of consciousness and opens new avenues for the science of consciousness, wellbeing, and psychiatry.

## Results

Given that the two control conditions (counting and memory) showed minimal differences for regional homogeneity (ReHo), connectivity gradients, and geometric eigenmodes, we averaged them for primary comparisons with ACAM-J. Full control-versus-control analyses and extended ROI tables are provided in the **supplementary materials**.

### Brain activity: ReHo

Comprehensive results/summary tables of significant regions are included in the **supplementary excel**. Here, we only provide a summary of the results. Across the eight ACAM-J, ReHo revealed systematic reorganization of brain activity (**Fig. 1a**), with increasing trends in posterior regions (bilateral visual cortex, precuneus–posterior cingulate, parietal, and somatomotor cortices), and decreasing trends in anterior and subcortical hubs (bilateral prefrontal [PFC] and orbitofrontal [OFC] cortices, brainstem regions associated with autonomic function (locs coeruleus, inferior medullary reticular formation). Higher-order polynomial fits revealed additional non-linear modulations, with quadratic trends in midline regions, and isolated cubic effects in posterior regions. Compared with the control conditions, ACAM-J showed reductions in prefrontal, orbitofrontal, temporal cortices and substantia nigra (SN, associated with arousal), and increase ReHo in posterior sensory regions (**Fig. 1b**). These changes followed a state-dependent trajectory: early ACAM-J were dominated by anterior and subcortical decreases, whereas later ACAM-J (J7–J8) displayed pronounced increases in visual and somatomotor cortices.

**Fig. 1.**
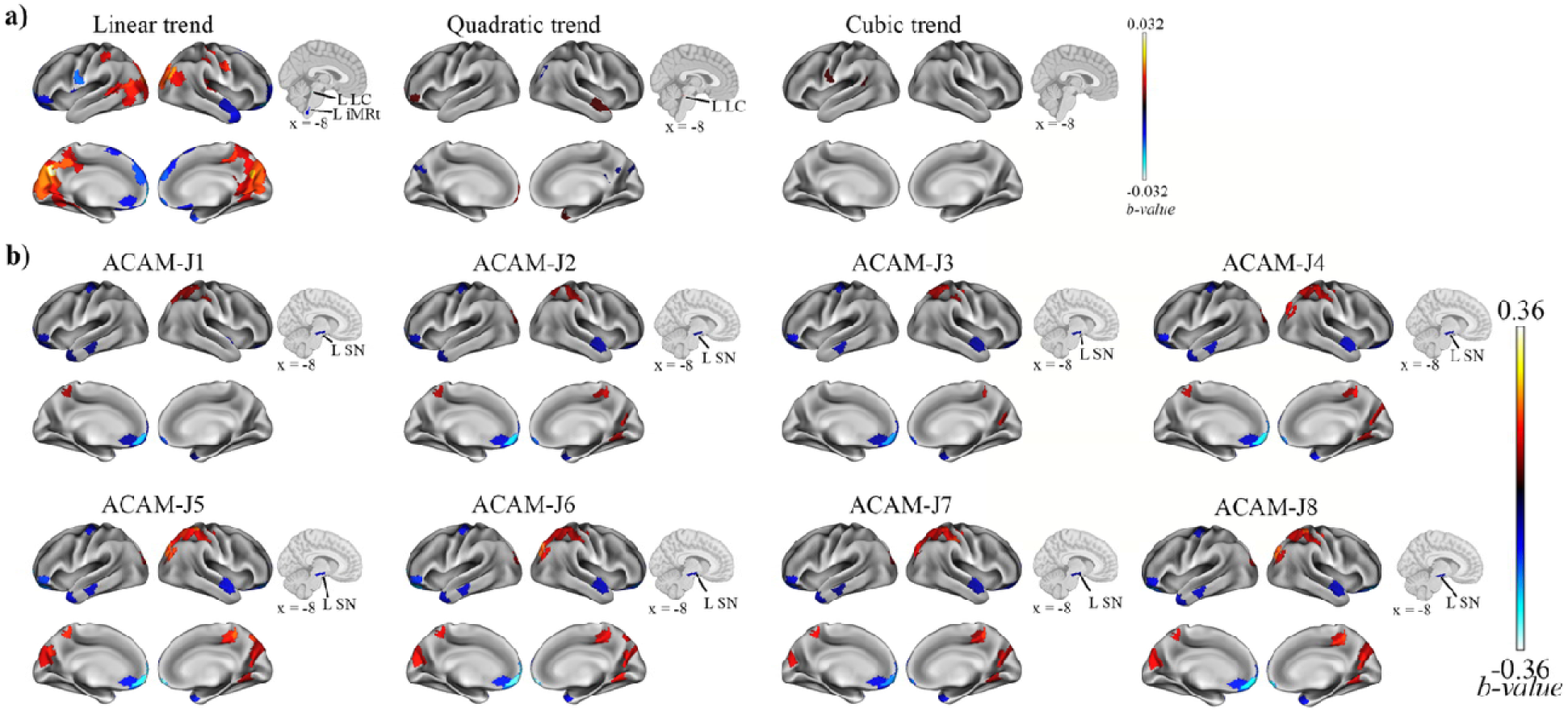
Distinct regional homogeneity (ReHo) during advanced concentration absorption meditation called jhana (ACAM-J). (a) Trends analyses show regions exhibiting linear, quadratic, and cubic trends across task ACAM-J. Linear trends highlights progressive and linear increase or decrease in specific brain regions while quadratic trends show mostly negative (inverted U-shaped) changes across ACAM-J. Positive but small cubic trend was found in the left somatomotor and parietal cortex. (b) ReHo differences between ACAM-J and the composite control reveal distinct patterns of cortical and brainstem substantia nigra activity.

### Macroscale hierarchy: Principal connectivity gradients

The principal connectivity gradient (G1) maintained a canonical unimodal-to-transmodal hierarchy during control tasks but was markedly flattened during ACAM-J (**Fig. 2a**). No significant trends were observed throughout ACAM-J. However, when compared directly to control conditions, ACAM-J was consistently associated with higher G1 values in visual and right temporal cortices and lower values in the left temporal pole (**Fig. 2b**). This pattern indicates that while hierarchical organization is globally compressed, regional reconfigurations are non-uniform and potentially reflect altered functional specialization during absorption. Full ROI-level results are reported in **Supplementary Table S3**.

**Fig. 2.**
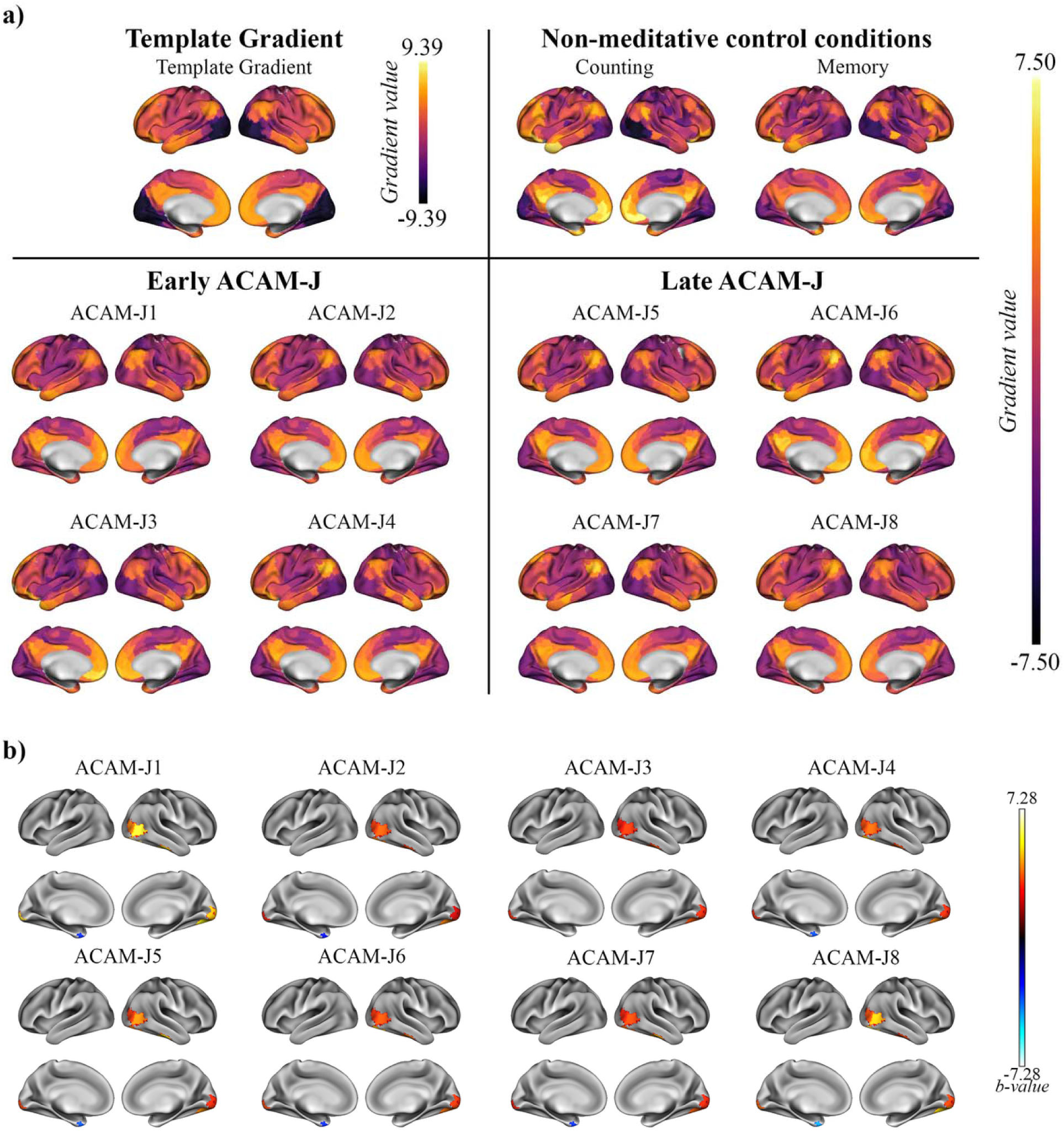
Principal connectivity gradient (G1) demonstrates functional hierarchical neural organization during ACAM-J. (a) Template principal gradient (G1) used for gradient alignment in this study, and average gradient maps for the non-meditative control conditions (counting, memory) and ACAM-J, grouped into early and late absorption phases. (b) Contrasts between individual ACAM-J and the composite of control conditions reveal robust changes along G1 in visual and temporal hubs highlighting regional-specific reorganization of large-scale cortical hierarchy during ACAM-J.

### Global cortical dynamics: Geometric eigenmodes

Overall mean power/energy were significantly lower during ACAM-J compared with controls, except for mean energy of ACAM-J1 and ACAM-J2 (**Fig. 3a–b**). Frequency-specific analyses showed the most remarkable differences between ACAM-J5-6 and control conditions (**Fig. 3c–d**). No significant interactions between eigenmodes and ACAM-J were observed (p > 0.05). In contrast, polynomial trend analyses uncovered a robust trajectory across ACAM-J. Both mean power/energy exhibited consistent negative linear and positive quadratic trends, indicating an early decline from ACAM-J1 through ACAM-J5 followed by recovery in later ACAM-J (**Fig. 3e–h**). Although both metrics followed the same U-shaped pattern, the quadratic effects were less pronounced for energy than for power. Together, these results point to a nonlinear reconfiguration of large-scale cortical harmonics during absorption, characterized by an initial compression of global dynamics followed by re-expansion in advanced absorption. Full polynomial coefficients and eigenmode-level statistics are provided in **Supplementary Tables S3**.

**Fig. 3.**
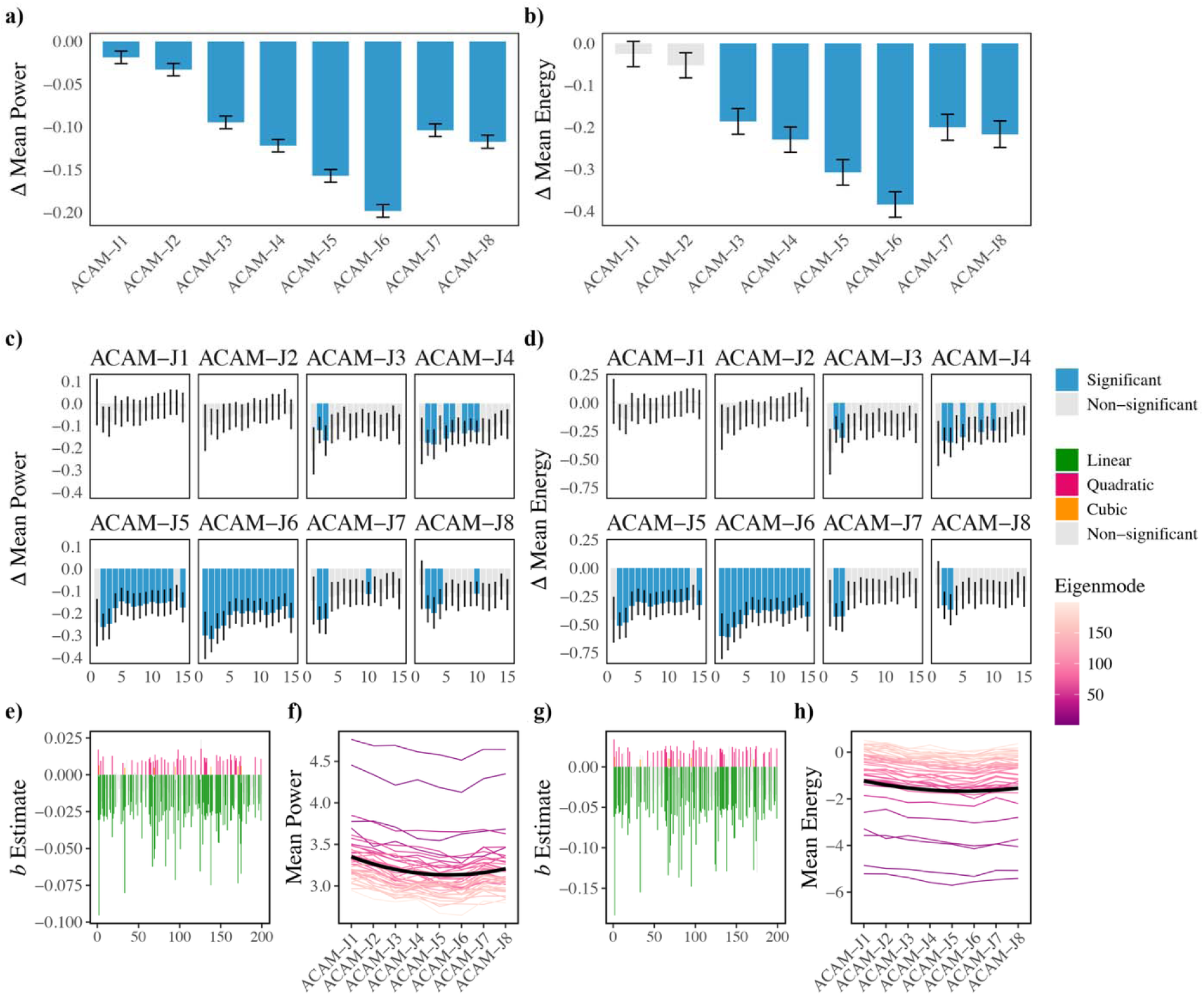
Geometric eigenmode decomposition reveals large-scale cortical dynamics during ACAM-J relative to non-meditative control conditions. Mean eigenmode power (a) and energy (b) during ACAM-J relative to the composite control condition. Eigenmode spectra for individual eigengroups highlight systematic reductions in power (c) and energy (d) during ACAM-J compared to control conditions, with strongest effects during mid-to-late ACAM-J. Plots (e, g) show distribution of significant trends across the eigenmodes indicate predominantly negative linear effects, reflecting progressive decreases with increasing absorption depth, along with a smaller number of quadratic effects. Plots (f, h) demonstrate changes in mean power (f) and mean energy (h) across ACAM-J, revealing positive quadratic trends: decreases in power and energy up to mid-ACAM-J, followed by recovery and increases through ACAM-J8. Error bars represent standard errors.

### Multivariate and univariate brain-experience relationships: Linking neuroimaging metrics with phenomenology, meditative traits, and cognition

To examine how neural dynamics relate to subjective and behavioral measures, we applied partial least squares correlation (PLSC) analyses linking ReHo, gradients, and eigenmodes with phenomenology, meditative traits, and cognition. For clarity, we report only the leading latent variables in the main text; full details of secondary LVs are provided in the **Supplementary results**.

#### Brain activity: ReHo

For ACAM-J phenomenology (39.52% shared covariance, p_FDR_ < 0.001; r = 0.80, p < 0.001; **Fig. 4a**), posterior sensory–attentional regions (visual, somatomotor, precuneus/posterior cingulate) loaded positively with correspondence of ACAM-J, stability, and width of attention, while anterior control and medial-temporal higher-order cognitive regions (medial/anterior prefrontal, ventral temporal/hippocampal, cerebellar), basal ganglia, and cerebellum loaded negatively with sensory experiences and narrative thought (**Fig. 4a**). Meditative traits (42.32% shared covariance, p_FDR_ < 0.001; r = 0.91, p < 0.001; **Fig. 4b**) showed primarily negative loadings, except for interoceptive awareness. This pattern was associated with positive weights in the medial frontal and posterior regions, temporal poles, and subcortical hubs (brainstem), and negatively in the lateral PFC, and visual regions. With cognition (27.27% shared covariance, p_FDR_ < 0.001; r = 0.94, p < 0.001; **Fig. 4c**), general cognitive performance (working memory, sustained attention, executive control) loaded positively with temporoparietal, dorsomedial PFC regions, and basal ganglia, while social cognition loaded negatively with lateral PFC, OFC, visual cortex, and brainstem.

**Fig. 4.**
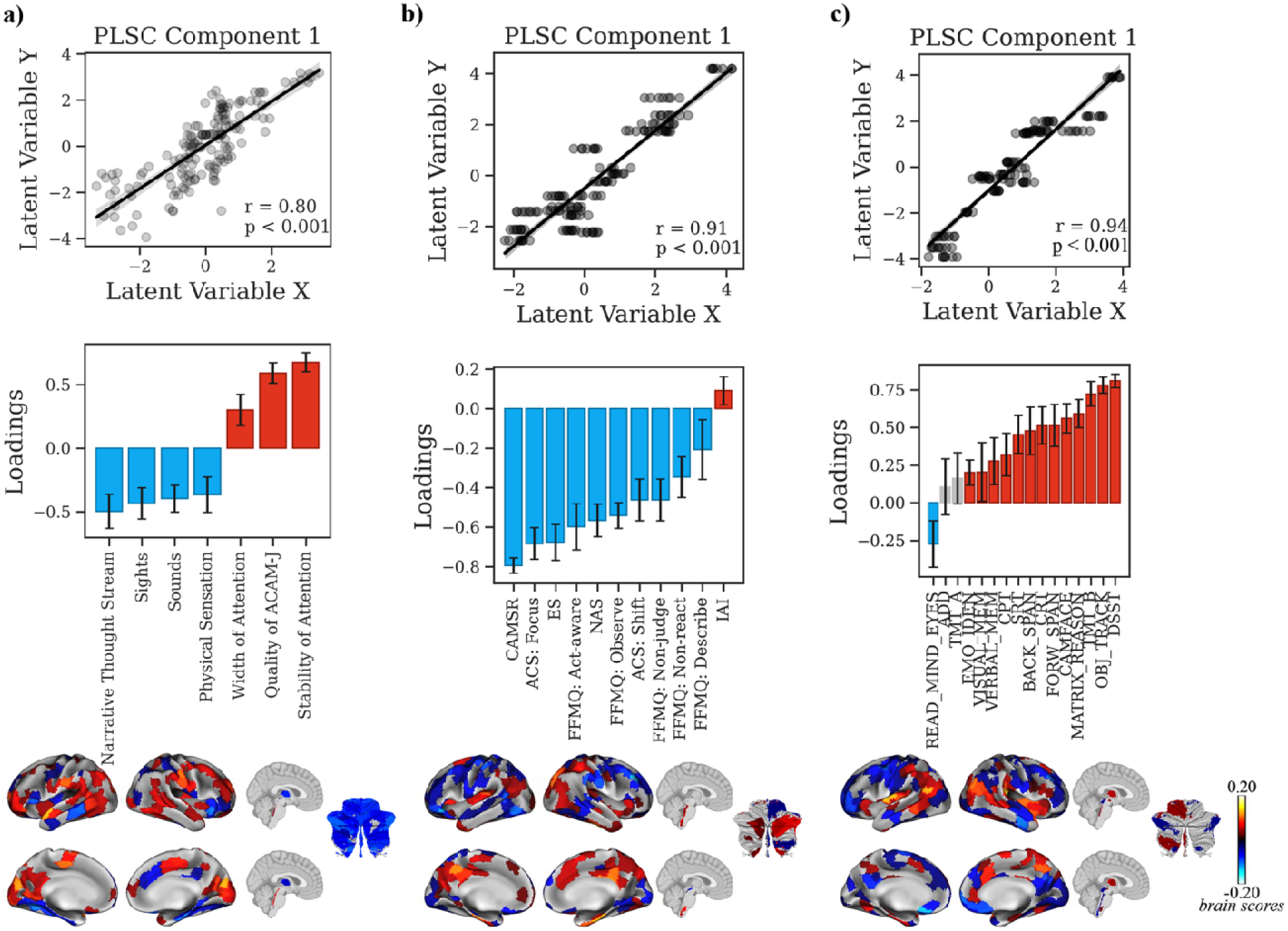
Multivariate relationship between ReHo during ACAM-J and behavioral patterns (ACAM-J phenomenology, meditative characteristics, and general behavioral performance). Partial least squares correlation (PLSC) analyses linking regional homogeneity (ReHo) during ACAM-J to three behavioral domains: (a) phenomenological ratings, (b) meditative characteristics, and (c) general behavioral performance. For each analysis, scatterplots of the first latent variable (top) show significant brain–behavior correlations (all p < 0.001). Bar plots (middle) display behavioral loadings, with error bars denoting bootstrap-estimated confidence intervals. Brain maps (bottom) depict regional ReHo loadings contributing to each multivariate pattern. Error bars represent standard errors.

#### Macroscale hierarchy: Principal gradients

Phenomenologically (49.27% shared variance, p_FDR_ = 0.018; r = 0.44, p < 0.001; **Fig. 5a**), higher G1 values in transmodal regions (medial/lateral PFC and anterior temporal regions) was associated with higher sensory vividness, whereas reduced unimodal loadings (visual, somatomotor, posterior parietal) were associated with attentional absorption (width of attention, stability of attention, and correspondence of ACAM-J). Meditative trait associations (40.75% shared variance, p_FDR_ = 0.0016; r = 0.51, p < 0.001; **Fig. 5b**) loaded positively on almost all traits, which covaried with loadings in transmodal regions (medial/lateral PFC, temporal poles), from non-significant descriptive/interoceptive awareness, which loaded on unimodal sensorimotor/parieto-occipital regions. For cognitive performance (38.29% shared variance, p_FDR_ < 0.001; r = 0.55, p < 0.001; **Fig. 5c**), social cognition mapped to sensorimotor/temporoparietal/visual regions, while general performance loaded on medial PFC and anterior temporal regions.

**Fig. 5.**
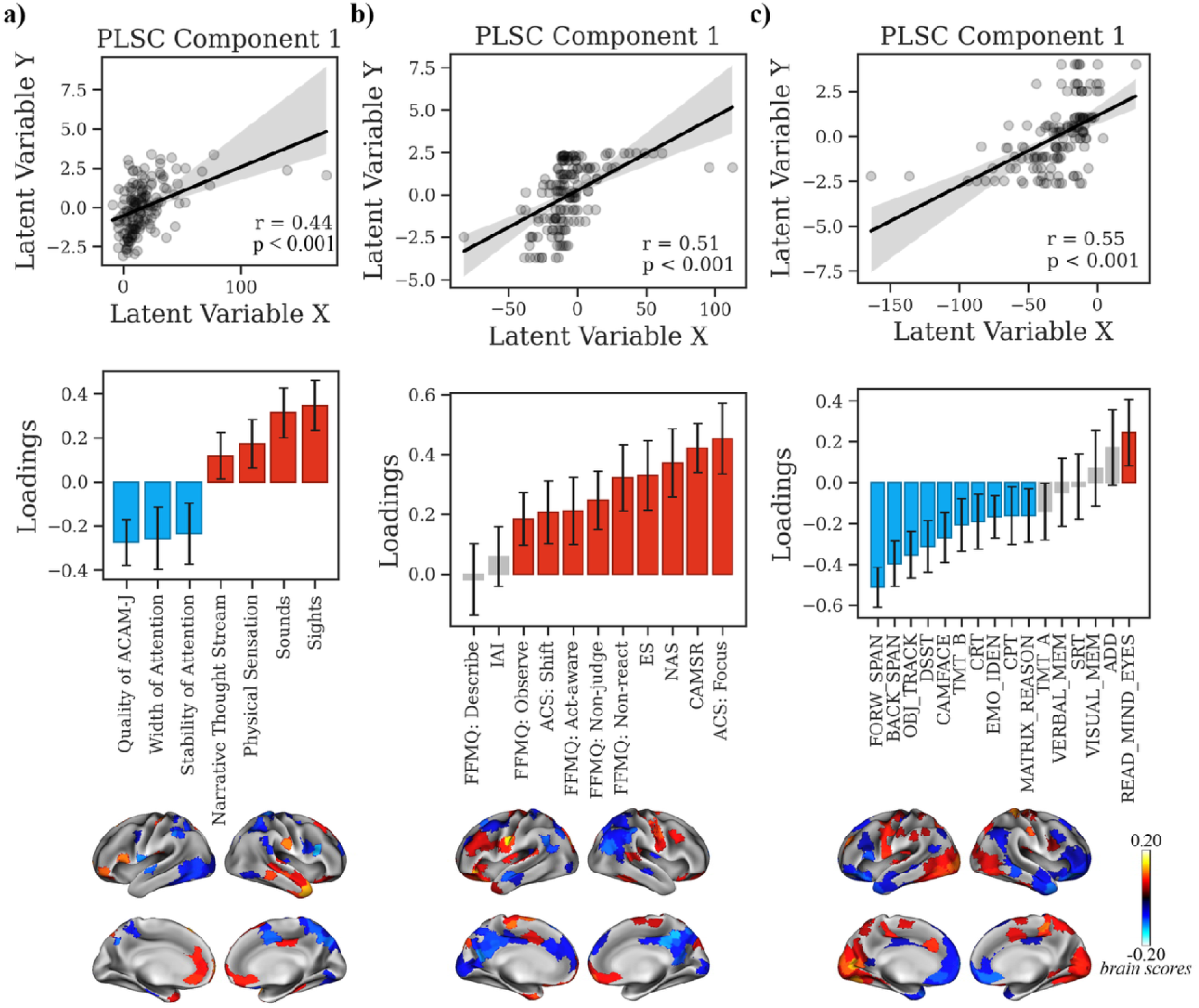
Multivariate relationship between G1 during ACAM-J and behavioral patterns (meditative characteristics, and general cognitive performance). Partial least squares correlation (PLSC) analyses linking principal gradient values (G1) during ACAM-J to three behavioral domains: (a) phenomenology, (b) meditative characteristics, and (c) general cognitive performance. For each analysis, scatterplots of the first latent variable (top) show significant brain–behavior correlations (all p < 0.001). Bar plots (middle) display behavioral loadings, with error bars denoting bootstrap-estimated confidence intervals. Brain maps (bottom) depict regional G1 loadings contributing to each multivariate pattern.

#### Global cortical dynamics: Geometric eigenmodes

Global spectral activation covaried positively together with sensation of sounds and oppositely phenomenology of sights and negative (97.00% shared variance, p_FDR_ < 0.001; r = 0.42, p < 0.001; **Fig. 6a**). In the meditative traits domain, global spectral activation positively linked to non-judgment and focus (98.47% shared variance, p_FDR_ = 0.006; r = 0.33, p < 0.001; **Fig. 6b**). In the cognitive domain, higher cognitive performance was associated with lower global spectral activation (99.46% shared variance, p_FDR_ < 0.001; r = 0.61, p < 0.001; **Fig. 6c**).

**Fig. 6.**
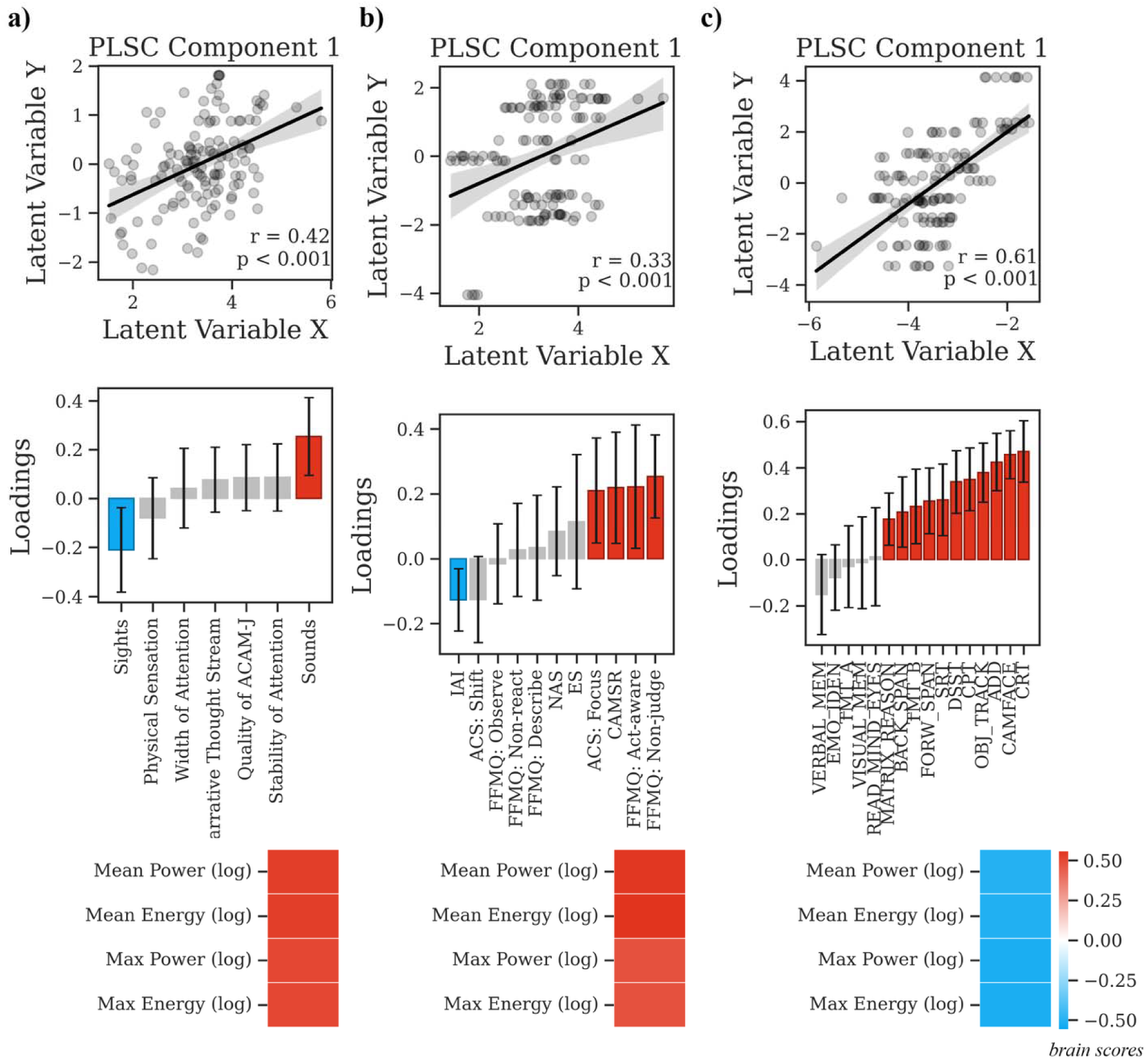
Multivariate relationship between global eigenmode activations during ACAM-J and behavioral patterns (meditative characteristics, and general cognitive performance). Partial least squares correlation (PLSC) analyses linking global eigenmode activations during ACAM-J to three behavioral domains: (a) phenomenology, (b) meditative characteristics, and (c) general cognitive performance. For each analysis, scatterplots (top) of the respective latent variable show significant brain–behavior correlations. Bar plots (middle) display behavioral loadings, with error bars denoting bootstrap-estimated confidence intervals. Heatmaps (bottom) show the global eigenmode activation loadings contributing to each multivariate pattern.

### Univariate brain-experience for ACAM-J factors

For ReHo, equanimity was selectively associated with reduced ReHo in the right hippocampus, while formlessness showed positive associations with temporal regions, and negative associations with prefrontal, precuneus-posterior cingulate cortex, PFC, and somatomotor cortex. For principal gradients, formlessness was positively associated with G1 values in the right medial cortex, and mean power/energy showed no associations with ACAM-J factors.

#### Psychobehavioral affinity map decoding

Across ACAM-J, we observed a highly consistent functional dissociation between attentional–executive versus affective–autobiographical psychobehavioral affinity maps (**Fig. 7**). Functions with positive associations with ACAM-J were dominated by attentional and control-related processes, including sustained and selective attention, spatial and visual attention, planning, action, and mental imagery. These domains were among the strongest and most consistent effects across all ACAM-J, indicating systematic engagement of executive and visuospatial control networks. In contrast, functions with negative associations with ACAM-J clustered around affective, autobiographical, and bodily–social processes, including emotion, mood, valence, fear, stress, loss, and anxiety, as well as autobiographical and episodic memory, social cognition, and eating. This pattern suggests systematic down-regulation of affective and self-referential functions during ACAM-J. Moreover, the magnitude of the correlations were larger as ACAM-J deepens toward ACAM-J8.

**Fig. 7.**
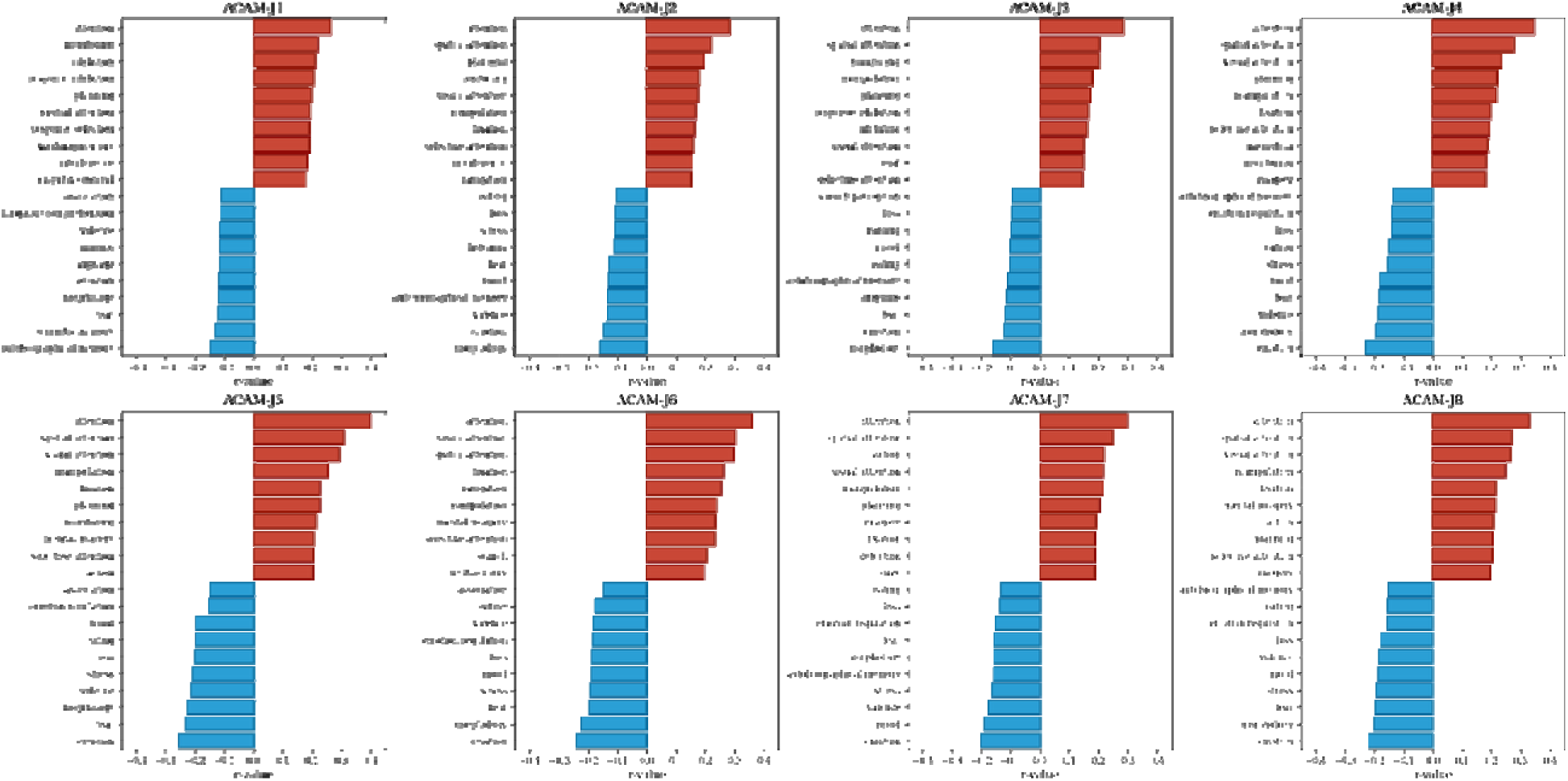
Reorganization of psychobehavioral domains during ACAM-J. For each ACAM-J, bar plots display the top ten functions with positive (red) and negative (blue) associations derived from Neurosynth decoding. Positive associations were dominated by attentional, executive, visuospatial, and action-related domains (e.g., sustained and selective attention, planning, imagery). In contrast, negative associations consistently involved affective, autobiographical, and bodily–social domains (e.g., emotion, mood, stress, fear, autobiographical memory, social cognition, eating, morphology). This dissociation highlights a robust psychobehavioral reorganization during ACAM-J, with enhanced attentional–executive functions accompanied by down-regulation of affective and self-referential processes.

## Discussion

This study provides the first multidimensional neuroscientific characterization of ACAM-J in a deeply phenotyped fMRI cohort (N= 20). Using ultra-high-field 7T fMRI and multiscale analyses, we show that ACAM-J constitutes a distinct, structured mode of consciousness marked by reproducible trends in local brain activity, macroscale gradients, global eigenmodes, and their associations with phenomenology, meditative traits, behavioral assessments, and psychological brain maps. Our findings reveal not only the capacity of the brain to volitionally sustain altered states of consciousness, but also its potential to evolve into structured, non-ordinary modes that align with meditative endpoints. These transformations may represent adaptive neurocognitive mechanism that support mental balance, clarity, and well-being, pointing to the brain’s endogenous capacity for human flourishing.

Using ReHo as an index of brain activity, we revealed a systematic redistribution of local activity: prefrontal and orbitofrontal hubs associated with narrative thought and cognitive control were progressively down-regulated, while posterior sensory and midline hubs (visual, somatomotor, precuneus/posterior cingulate) became more active. Furthermore, brainstem regions associated with arousal (locus coeruleus) and autonomic function (inferior medullary reticular formation) showed reduced activity throughout ACAM-J. We replicated findings from our previous intensive case study, demonstrating that early ACAM-J is characterized by lower activity frontal regions, while later ACAM-J is characterized by enhanced posterior sensory activity (*14*). Here we show that anterior-to-posterior reorganization reflects a transition from conceptual, effortful processing (*33–36*) toward refined, subtle, and open awareness (*14, 37*). Moreover, the progressive reduction of activity in the locus coeruleus and inferior medullary reticular formation suggests a gradual disengagement of sympathetic autonomic activity (*38*), consistent with the transition toward equanimity and diminished bodily awareness characteristic of later ACAM-J (*18*). Together, ACAM-J may be understood as producing a state-specific reallocation of resources away from conceptual elaboration toward stabilized sensory-attentional processing.

To probe the low-dimensional architecture of functional connectivity during ACAM-J, we assessed principal connectivity gradients. Connectivity-gradient analyses indicated that while the canonical unimodal–transmodal axis was preserved during control tasks, it was markedly flattened during ACAM-J. Across ACAM-J, principal gradient undergoes a systematic compression, indicating a progressive collapse of hierarchical differentiation between unimodal and transmodal systems (*25*). Importantly, the effect was not globally uniform: specific regions such as visual and temporal cortices shifted upward along the gradient, while the left temporal pole shifted downward. This non-uniform reconfiguration may reflect the brain’s transition from a modular architecture toward a globally diffused yet functionally specialized coherent, integrative conscious field that may support stable, non-conceptual awareness (*39, 40*).

Complementing this at the global scale, geometric eigenmode analysis revealed a U-shaped trajectory across ACAM-J where early ACAM-J were characterized by compression of spectral power and energy, followed by re-expansion in later ACAM-J. This dynamic reconfiguration suggests that absorption transiently downregulate cortical harmonics before reintegration at a higher level of coordination. This brain dynamic may reflect a brain mode that support focused attention, mental stillness, and reduced sensory processing typical of absorption in meditation (*41, 42*). Surprisingly, our findings contrast with our previous case study (*27*) and also a prior studies involving psychedelic substances (*43, 44*), which both reported expansion and increased activation cortical brain mode.

One possible explanation for this discrepancy could be that the current findings are based on group-level data, capturing population-wide patterns that may smooth over individual variability, whereas the previous case study focused on a single individual, potentially reflecting idiosyncratic dynamics. One could also argue that studies involving psychedelic substances have a similar number of participants to our study with ACAM-J. However, a key difference between these studies and ours lies in the range of eigenmodes used for analysis. While research on connectome eigenmodes typically explores a range of up to 10,000 eigenmodes (*43, 44*), our study focuses on geometric eigenmodes, using a range of up to 200, in line with standard methodology (*45*). As a result, the lower activation energies captured through geometric eigenmodes in our study may not represent the higher activation energies observed in the higher eigenmodes associated with psychedelic studies. Nevertheless, our findings may reflect a controlled reintegration of complexity, suggesting that similar altered states of consciousness can emerge via distinct neurodynamic pathways.

Strikingly, multivariate analyses using PLSC demonstrate that ACAM-J is not reducible to isolated neural or psychological effects but reflects an integrated reorganization of brain–mind dynamics. Posterior, parietal, and sensory hubs appear to support attentional qualities, while prefrontal systems contribute to the suspension of narrative and sensory processing. This was evident in both mesoscale activity and macroscale hierarchy of the brain, and eigenmode dynamics suggests that the brain is reconfigured to sustain enduring meditative traits. These convergences establishes the neurophenomenology of ACAM-J (*46*) where phenomenology, traits, behavior, and neural signatures converge to a unified phenomenon. More importantly, the neural configurations suggest a disengagement from self-referential processing and ordinary perception, marking a transformative shift toward enhanced sensory and attentional processing (*47, 48*). This reorganization reflects a more flexible, refined engagement with the present moment, and moves beyond ordinary narrative-driven cognition to a state of heightened, dynamic sensory awareness.

Beyond neurophenomenology, we also examined the psychobehavioral affinity domains most associated with ACAM-J using Neurosynth decoding. A striking and robust dissociation emerged in which attentional and executive domains were consistently positively associated with brain activity during ACAM-J, while negative affect, autobiographical, and self-referential domains were systematically downregulated. This functional reorganization indicates that ACAM-J is associated with a structured shift toward enhanced attentional stability and reduced negative affect and self-referential processing. Importantly, this mirrors practitioners’ first-person reports of effortless attentional absorption and awareness, bliss, happiness, and equanimity. These results emphasize ACAM-J not as a passive withdrawal of consciousness but an active reconfiguration of neural dynamics toward sustained attention and perceptual clarity, accompanied by systematic down-regulation of affective–autobiographical functions. Thus, our neurophenomenological findings highlights the value of neurophenomenology in studying non-ordinary states of consciousness such as ACAM-J with the same rigor as perception or memory (*49–52*).

Our findings directly contest leading models of consciousness. GNWT proposes that consciousness depends on ignition across a prefrontal–parietal “rich club” (*28*). This ignition process enables flexible access, working memory, and goal-directed cognition (*53*). In this view, conscious states should be accompanied by widespread prefrontal–parietal activation and recurrent signaling through thalamocortical loops (*54, 55*). Yet, our findings directly challenge this assumption. ACAM-J, and more specifically higher ACAM-J, marked by minimal content, was associated with prefrontal downregulation and increased posterior–midline activity. Rather than broadcasting through a prefrontal workspace, consciousness here persisted in the posterior regions. Importantly, ACAM-J is not characterized by dullness or reduced awareness. ACAM-J involve heightened awareness, clarity, and even refined forms of metacognition, indicating a state of open yet focused consciousness (*56*).

This combination of expansive awareness and highly refined attentional stability appears paradoxical within standard cognitive frameworks, which often treat focused attention and open awareness as mutually exclusive. Such paradoxes may instead reveal the limits of current theories in explaining absorptive states, rather than contradictions in the phenomenology itself. This suggests that while GNWT may capture consciousness in the service of ordinary cognition, it does not account for states of conscious awareness without prefrontal broadcasting. In this sense, ACAM-J exposes a boundary condition for GNWT, that prefrontal “workspace” activity may not be necessary for consciousness per se, but rather perhaps for consciousness in service of cognitive control.

Similarly, IIT proposes that the level and richness of consciousness depends on the amount of integrated information a system can generate (*29*). A system is most conscious when it is both highly differentiated and highly integrated (*57, 58*). Our findings challenge the basic principle of this model. During ACAM-J, cortical gradients become diffused, indicating reduced differentiation between unimodal and transmodal systems. At the same time, geometric eigenmodes showed coherent dynamics in a low-dimensional space. Phenomenology remained vivid despite this reduction in modular complexity. One interpretation is that while the quantity of content to be integrated decreases, the quality of integration among the remaining processes is exceptionally strong. This results in an integration high in absorption and unity greater than in ordinary experiences (*56*). Our results suggest that consciousness does not always scale linearly with differentiation and integration. Instead, consciousness of absorption states such as those accompanying bliss, openness, and awareness of subtle qualities of the mind may emerge from simplified yet strongly coordinated neural signatures. From this perspective, ACAM-J both challenges and partly aligns with IIT, indicating that integration can sustain vivid consciousness even when differentiation is low.

MPE describe states in which consciousness is preserved in its most reduced form and are devoid of narrative thought, sensory imagery, and affective tone, yet still minimally aware (*32*). ACAM-J supports this view but extends it by showing a structured, trainable trajectory of content-minimal states unfolding in reproducible neural reorganizations. Unlike passive or spontaneous MPE, ACAM-J offers a progressive refinement of consciousness into such states. Our data supports this by showing dominant posterior activity, reduced prefrontal activity, and their associations with ACAM-J phenomenology. Together, these results suggest that consciousness is not bound to ignition or complexity, but can reorganize into globally coherent and open modes, offering GNWT and IIT boundary conditions and an extension for MPE.

The implications extend beyond the contemplative sciences. The neural correlates of ACAM-J may have profound consequences for our understanding of mental health, well-being, and the potential for human transformation (*15, 16*). The ability to selectively intensify executive functions and self-generate bliss and joy while downregulating emotional reactivity, autobiographical and self-referential processing suggests a mode of brain function that can be optimized for clarity, non-reactivity, and present-centered awareness in daily life. This neural pattern directly inverts patterns observed in psychopathology, such as rumination in depression (*59*), hyper-reactivity in anxiety (*60*), and craving in addiction (*61, 62*). Instead of the disrupted prefrontal–limbic activity seen in these conditions, ACAM-J is characterized by lower prefrontal activity, higher posterior/midline activity, patterns that covaried with equanimity-related traits and were anticorrelated with psychological suffering. Equanimity has been proposed as a critical outcome for contemplative training and a contributor to stress resilience (*37, 63–67*), and advanced meditation such as ACAM-J and advanced investigative insight meditation explicitly cultivates deep absorption and equanimity compared to standard mindfulness programs (*17, 68*).

Moreover, cultivation of self-generated bliss and joy during early ACAM-J represents a transformative aspect of meditation practice with profound implications for mental health. Bliss, as experienced in ACAM-J, is not merely a fleeting positive emotional state but a self-generated, stable source of joy that emerges through absorption. The neural correlates of such ACAM-J suggest a shift away from self-referential processing towards a more unified, present-centered awareness that is richly positive and deeply fulfilling. This experience of joy, free from external stimulus, could fundamentally frameworks of psychiatric care by providing individuals with an internal reservoir of emotional well-being. Unlike conventional therapeutic models that focus on alleviating negative symptoms, practices like ACAM-J offer a means of fostering positive emotional states, potentially rewiring the brain to maintain these states even in the face of adversity.

In this context, advanced meditation could therefore provide a powerful naturalistic model of well-being and flourishing achieved through advanced mental training rather than pharmacological or technological interventions. Cultivation of intrinsic joy and bliss through advanced meditation may offer a novel pathway to alleviate cycles of rumination, self-reference, and emotional reactivity that characterize many psychiatric conditions. In essence, rather than merely reducing negative symptoms, ACAM-J could actively promote the cultivation of positive states, fostering emotional balance, and a profound sense of well-being. On this basis, we hypothesize that ACAM-J may result in stronger or qualitatively distinct well-being relative to standard mindfulness meditation, though direct comparative trials are lacking and are needed to test this prediction (*15, 16*).

Potential clinical applications of ACAM-J could include neuromodulations (*69*) to facilitate development of ACAM-J factors, or other advanced meditation states during meditative development (*70*). Techniques such as transcranial magnetic stimulation or focused ultrasound could be explored to target neural circuits associated with absorption, equanimity, and or self-generated bliss and happiness in clinical populations (*71*). Additionally, neurofeedback and closed-loop brain-computer interfaces may offer non-invasive methods to train and stabilize advanced meditation states. Ultimately, our study helps lay the groundwork for a rigorous science of advanced meditation, its states, development, endpoints, and flourishing, with profound implications for mental health and human well-being.

Notwithstanding its merits, this study has several limitations. First, even highly trained meditators may face challenges in reproducing deep absorption within the noisy MRI environment, and atypical supine position for practicing ACAM-J. Such ecological mismatch could alter phenomenology relative to naturalistic practice. However, we mitigated this by having participants practice ACAM-J with the MRI noise recording while laying supine prior to data collection. Moreover, the convergence across multiscale neural and experiential measures suggests that the core neural correlates of ACAM-J were reliably captured. The absence of electrophysiological recordings such as EEG or MEG limits characterization of fast spectral–temporal features such as microstates or non-linear oscillatory dynamics (*26*). The current findings may not generalize to other forms of absorption or concentration-based practices, which may differ in techniques, phenomenology, and neural signatures. Finally, deeper ACAM-J are increasingly subtle and inherently more difficult to sustain in the MRI system. While participants were able to enter these states, the requirement of button-press confirmation may perturb the absorption process. Unequal ACAM-J dwell times could also bias trajectory estimates. Nevertheless, participants, on average, maintained balanced durations across ACAM-J, mitigating this concern. Moreover, participants also indicated high quality of phenomenology of each ACAM-J and were able to indicate access to these eight states. Future research should incorporate multimodal recordings and longitudinal training designs to test and extend the results of the current study.

Our findings demonstrate that human consciousness is not fixed to its default modes but can be intentionally reconfigured into eight distinct absorption states marked by profound ease, stability, equanimity, and reduced ordinary perception and cognition. The eight distinct ACAM-J challenge prevailing neuroscientific models and theories by demonstrating that the brain harbors latent capacities capable of counteracting the maladaptive neural patterns that underlie rumination, craving, and hyper-reactivity. ACAM-J offers a novel experimental model of resilience, mental health, and flourishing, and demonstrates that the human brain carries within it the latent ability to catalyze its own transformative potential.

## Materials and Methods

### Participants

Participants were recruited as part of a larger study that aims to investigate neurophenomenology of various advanced meditation practices. These experienced meditators come from well-established meditation communities, all of whom demonstrated advanced expertise in ACAM-J meditation practices. The sample consisted of 20 participants, with an age range of 26-67 years old (mean age = 43.95 ± 15.10 years).

On average, participants had 17.40 ± 11.59 years of meditation experience and reported an estimated total of 20500.40 ± 17288.98 hours of dedicated meditation practice. Participants were further categorized based on their engagement with distinct styles of ACAM-J meditation, including intermediate (N=18), and deep (N=2) ACAM-J. The study received approval from the Mass General Brigham Institutional Review Board (IRB) and the ethics boards at Witten/Herdecke University. Participants also provided informed consent.

### Experimental design

#### Advanced concentrative absorption meditation (ACAM-J)

Participants followed their standard ACAM-J sequence with closed eyes, progressing from access concentration through ACAM-J1 to ACAM-J8, concluding with a post-ACAM-J8 state referred to as *afterglow*. We requested participants to stay within any ACAM-J for at least 3 minutes for reliable computation of fMRI metrics (*72*). For participants with ACAM-J duration of less than 2 minutes, we conducted another session of ACAM-J. The participants marked transitions from access concentration to ACAM-J8 with a button press. Note that 1 participant did not indicate transitions from ACAM-J6 to ACAM-J7 nor from ACAM-J7 to ACAM-J8, as it would disrupt the natural flow of ACAM-J practice for them. In total, 22 runs of ACAM-J were recorded, and the average duration for a complete run covering ACAM-J1 to ACAM-J8 was 33.60 ± 15.40 min (for more details on state duration, refer to Table S1).

#### Non-meditative control conditions

Two non-meditative control conditions were developed for comparison with meditation states, aimed at engaging the participant’s mind in non-meditative, ordinary cognitions while avoiding the induction of meditative states. Resting-state control was not used due to concerns that experienced meditators might enter meditative states during rest, which may complicate interpretations of results (*73*). These control conditions consider the phenomenology of meditative states and the nature of the experimental paradigm to ensure greater consistency and reliability in the intensive case-study design (*74*). The two control conditions involved a memory task, where the participant recalled events from the past one week, and a counting task, where the participant mentally counted down from 10,000 in decrements of 5. One 8 min runs of each non-meditative control were collected for most participants. Five participants who were recruited additionally for separate case studies conducted two sessions of the control tasks. In total, we collected 25 sessions of control tasks. Please note that some participants reported experiencing a small degree of ACAM-J during the control conditions. However, these meditative qualities were not as intense as the meditation states experienced during their meditation runs. Moreover, effects of ACAM-J during control conditions did not impact the differences between ACAM-J and non-meditative controls in prior analyses (*74*).

#### Phenomenology

To complement objective neuroimaging data, this study design also used a first-person phenomenological approach, in which the subject assessed the mental and physiological processes relevant to the experience of ACAM-J as they manifested during meditation. In brief, following a complete ACAM-J run (ACAM-J1 to ACAM-J8), the participant assigned a rating from 1 to 10 for phenomenological items including: (1) *stability of attention*, ranging from poor to excellent stability; (2) *width of attention*, varying from a narrow scope like a laser to very wide like a fisheye lens; (3) *quality of ACAM-J*, indicating the degree of alignment between experience of the ACAM-J quality and the defining features of the corresponding ACAM-J; (4) *early phenomenology of sights, sounds, physical sensations, and narrative thought stream*, denoted by the presence of sights, sounds, physical sensations, and narrative thought contents during the form ACAM-J (ACAM-J1-J4); (5) *late phenomenology of sights, sounds, physical sensations, and narrative thought stream*, characterized by the presence of sights, sounds, physical sensations, and narrative thought contents during the formless ACAM-J (ACAM-J5-J8); and (6) qualities of specific ACAM-J (ACAM-J factors) rated specifically for that ACAM-J, including *bliss/joy* (ACAM-J1-2), *contentment* (ACAM-J3), *equanimity* (ACAM-J4), and *formlessness* (ACAM-J5-8). Higher rating values indicated a more robust experience within the given item. *Stability of attention, width of attention,* and *quality of ACAM-J* were rated for each ACAM-J. Phenomenology of *sights, sounds, physical sensations, and narrative thought stream* were rated separately for early (ACAM-J1-J4) and late (ACAM-J5-8) aggregate states.

#### Questionnaires

Participants also completed several questionnaires regarding meditative traits as part of a larger study on advanced meditation. Here, we focused on questionnaires relevant to this study.

*Five facet mindfulness questionnaire (FFMQ-15).* The FFMQ-15 is a shortened version of the original 39-item measure, assessing five distinct facets of mindfulness: observing, describing, acting with awareness, non-judging of inner experience, and non-reactivity to inner experience (*75*). Each facet is measured by three items rated on a 5-point Likert scale (1 = “never or very rarely true” to 5 = “very often or always true”).

*Cognitive and Affective Mindfulness Scale-Revised (CAMS-R).* The CAMS-R is a 12-item self-report measure designed to capture mindfulness as a unified construct encompassing attention, awareness, present-focused acceptance, and non-judgment (*76*). Items are scored on a 4-point Likert scale (1 = “rarely/not at all” to 4 = “almost always”), with higher scores indicating greater mindfulness.

*Non-attachment scale-short form (NAS-SF).* The NAS-SF measures the Buddhist psychological construct of non-attachment, defined as a flexible, balanced way of relating to one’s experiences without clinging to or suppressing them (*77*). The 8-item scale assesses the extent to which individuals can engage in life with less fixation on outcomes and experiences. Items are rated on a 6-point Likert scale (1 = “strongly disagree” to 6 = “strongly agree”), with higher scores indicating greater non-attachment.

*Equanimity scale (ES-16).* The ES-16 measures the ability to maintain mental balance regardless of whether experiences are pleasant, unpleasant, or neutral (*78*). This 16-item self-report measure captures two subscales: experiential acceptance, an attitude which does not seek to resist or attach to the experience and involves acceptance of all internal experiences (thoughts, feelings, body sensations, etc.), and non-reactivity, which is non-reactivity to experiences preventing attachment or aversion to these experiences (e.g. thoughts, feelings). Items are rated on a 5-point Likert scale (1 = “not at all” to 5 = “very much”), assessing even-minded states and non-preferential reactivity to experiences.

*Attentional Control Scale (ACS).* The ACS is a 20-item scale assessing attentional control, focusing on the ability to focus, shift attention between tasks, and flexibly control thought (*79*). ACS has two subscales: attentional focusing, which is the capacity to intentionally hold the attentional focus, resisting unintentional shifting to irrelevant or distracting stimuli, and attentional shifting, which is the capacity to intentionally shift the attentional focus to the desired target, avoiding unintentional attention. Items are rated on a 4-point Likert scale (1 = “almost never” to 4 = “always”), with higher scores indicating better attentional control.

#### Cognitive tasks

We examined six 16 standardized tasks assessing various cognitive domains using TestMyBrain (TMB) Digital Neuropsychology Toolkit (*80*). Normative data for the TMB battery, collected from samples of 4000–60,000 individuals aged 12–90 years, are stratified by age, gender, level of education, and type of device used. Specific tests include domains of processing speed: Simple Reaction Time (SRT; measure of basic processing speed), Choice Reaction Task (CRT; measure of decision-making speed with multiple stimuli), Digit Symbol Matching (DSST; measure of processing speed), Trail Making Tests Part A (TMT_A; measure of processing speed); cognitive control: Trail Making Tests Part B (TMT_B; measure of cognitive flexibility), Gradual Onset Continuous Performance Test (CPT; measure of sustained attention and vigilance); working memory: Paced Serial Addition Test (ADD; measure of sustained attention and working memory), Forward and Backward Digit Spans (FORW and BACK_SPAN; measure of working memory), Multiple Object Tracking (OBJ_TRACK; measure of visuospatial attention and visual working memory); reasoning: Matrix Reasoning (MATRIX_REASON; measure of visual reasoning and general cognitive ability); long-term memory: Verbal and Visual Paired Associates Memory Tests (VERBAL_MEM and VISUAL_MEM; measures of verbal and visual memory, respectively), Cambridge Face Memory Test (CAMFACE; measure of face recognition), and social function: Multiracial Emotion Identification (EMO_IDEN; measure of face emotion recognition), and Reading the Mind in the Eyes (READ_MIND_EYES; measure of mental state inferencing and theory of mind). For more information about these measures and the domains that they measure, please refer to **Supplementary Table 1**.

#### Neuroimaging acquisition

Neuroimaging scans were obtained utilizing a 7T magnetic resonance (MR) scanner (SIEMENS MAGNETOM Terra) equipped with a 32-channel head coil. Functional imaging utilized a single-shot two-dimensional echo planar imaging sequence with T2∗-weighted BOLD-sensitive MRI. The imaging parameters included a repetition time (TR) of 2.9 s, echo time (TE) of 30 ms, flip angle (FA) of 75 degrees, field of view (FOV) of [189 × 255], matrix of [172 × 232], GRAPPA factor of 3, voxel size of 1.1 × 1.1 × 1.1 mm³, 126 slices, interslice distance of 0 mm, bandwidth of 1540 Hz/px, and echo spacing of 0.75 ms. Slice acquisitions covered the entire brain, with interleaved slices, sagittal orientation, and anterior-to-posterior phase encoding. Additionally, opposite phase-encoded slices (i.e., posterior-to-anterior) with identical parameters were acquired for distortion correction.

Whole-brain T1-weighted structural images were acquired with the following parameters: TR of 2.53 s, TE of 1.65 ms, inversion time of 1.1 s, flip angle of 7 degrees, 0.8 mm isotropic resolution, FOV of 240 × 240, GRAPPA factor of 2, and bandwidth of 1200 Hz/Px. Physiological signals, including heart rate and respiration, were continuously monitored throughout the scanning session. Heart rate was measured using a piezoelectric finger pulse sensor (ADInstruments, Colorado Springs, CO, USA), and respiration was recorded using a piezoelectric respiratory bellow (UFI, Morro Bay, CA, USA) positioned around the chest. The signals were acquired using a PowerLab data acquisition system (PowerLab 4/SP, ADInstruments, New Zealand) and LabChart (LabChart, ADInstruments, New Zealand) at a sampling rate of 1000 Hz. The same acquisition system also recorded MR scanner trigger pulses synchronized with the acquisition of each image volume.

#### Neuroimaging preprocessing

Detailed processing steps can be found in a previous case study (*14*). In brief, preprocessing pipeline encompassed the following steps: 1) de-spiking, 2) RETROspective Image CORrection (RETROICOR), 3) slice time correction, 4) distortion correction, 5) motion correction, 6) registration of the anatomical dataset (T1w) to a standard MNI template, 7) scrubbing and removal of any volume with motion >0.3 mm and more than 5% outlier voxels, 8) regression of eroded cerebrospinal fluid (CSF) mask time course and motion parameters (3 translations, 3 rotations) per run, and finally band-pass filtering (0.01–0.1 Hz). Each fMRI run was then partitioned into distinct segments corresponding to different ACAM-J. Finally, 9) the bias field of the T1 image was corrected using SPM12 (*82*).

#### Regional homogeneity

Regional homogeneity (ReHo) is a measure of similarity in temporal activation pattern of a voxel and its nearby voxels (*83*). This measure of local functional connectivity within brain regions is a close derivative of underlying brain activity (*83*). A higher ReHo value indicates stronger synchronization of local brain regions, indicating greater functional connectivity in that region and thus an index of greater brain activity. In this study, we defined a cluster size of 27 and calculated the ReHo values for each fMRI segment, standardized the ReHo values before smoothing the standardized ReHo maps using a 2mm full-width half-maximum (FWHM) kernel. The standardized ReHo values were then parcellated using the four different parcellation/segmentation schemes (*84–87*), yielding 498 ReHo values for each fMRI segment.

#### Connectivity gradient-mapping

To map the gradients, we first computed a template gradient through a multi-step process. First, data was smoothed using a 6 mm FWHM kernel, followed by computation of functional connectivity matrices for each segment using the Schaefer 400-regions 7-network atlas using Pearson’s correlation, thus producing a 400×400 functional connectivity matrix. These individual matrices were then averaged across all segments, resulting in a mean functional connectivity matrix. Subsequently, gradients for this mean matrix were derived by thresholding the mean correlation matrix at 90% sparsity to retain only the strongest connections, followed by generating a similarity matrix using cosine similarity to capture the connectivity pattern similarities among ROIs. The resulting similarity matrix served as input for the diffusion map embedding algorithm, from which the gradients were extracted. We computed a template gradient from an out-of-sample dataset of 134 subjects from the HCP dataset (the validation cohort used by (*88*)).

Following template gradient computation, alignment with individual segment gradients was performed. The gradient values for each fMRI segment were computed based on their respective functional connectivity matrices and then aligned with the template gradient using Procrustes rotations with 10 iterations to ensure optimal comparison across datasets. All analyses described herein were conducted using the *BrainSpace* toolbox implemented in Python (*88*). Using the gradient values obtained from gradient alignment, we computed the average network gradient values for each network by averaging the gradient values of all the parcels within that network. To ease interpretation and reduce the complexity of the current manuscript, we focused only on the principal gradients (G1). An extended analysis of the connectivity gradients may be conducted in future studies.

#### Geometric eigenmode decomposition

All fMRI data was smoothed with surface-smoothing of 6 mm FWHM, and was converted to CIFTI data format for further processing *using ciftify_recon_all* and *ciftify_subject_fmr i*(*89*). To decompose fMRI activity into 200 frequency-specific eigenmodes, we used geometric eigenmodes derived from a population-averaged midsurface thickness template of the neocortical surface from Pang et al., wherein the methods to generate these geometric eigenmode templates are also described (*45*). Briefly, the geometric eigenmodes were computed by constructing the Laplace-Beltrami operator from the cortical mesh and solving the eigenvalue problem. The resulting eigenvalues are ordered according to spatial frequency/wavelength of each mode, where mode 1 has the longest wavelength (lowest frequency) while higher modes have shorter wavelengths (higher frequency). Conversely, eigenmodes are the complete basis set—a weighted sum of varying-wavelength modes.

Using the geometric eigenmodes, we decomposed functional MRI data to reveal emerging cortical spatiotemporal dynamic patterns by examining the contribution of each eigenmode to the cortical activity at each time instance. At each time point of each fMRI time courses, fMRI data were projected onto each of the 200 geometric eigenmodes, yielding the temporal activity of the particular eigenmode in a method described previously (*44, 90*). Subsequently, the temporal activation of eigenmodes were analyzed in terms of their power and energy at each fMRI time point. Here, power is defined as the strength of an eigenmode’s activation at a given fMRI time instance: |ω_k_(t_i_)|. Alternatively, energy is the eigenmode’s frequency-weighted contribution, estimated by weighting the square of the eigenmode’s strength of activation by the square of its corresponding eigenvalue (i.e. its intrinsic energy (*λ_k_*)) at each time point: |ω_k_(t_i_)|^2^ *λ_k_^2^*. Mean power and energy were calculated by averaging the power and energy of each eigenmode across all time points in a single segment (i.e. yielding 200 mean power and energy values for 200 modes per fMRI segment). Maximum power and energy were calculated as the maximum spectra value for a given eigenmode across all time points within a segment (i.e. yielding 200 maximum power and energy values for 200 modes per fMRI segment). Unless otherwise noted, all reported maximum and mean power and energy values are reported on a log-scale (*44*). Brain total power and total energy were computed by summing power and energy across all eigenmodes for each time point and then averaging all time points within a segment. We focus on global mean power and energy patterns, trends, and frequency-specific mean power and energy across ACAM-J in the main results. We present the same analyses, but for max power and energy patterns in the supplementary results.

#### Decoding of brain activity using data-driven Neurosynth psychobehavioral affinity maps

To provide a complementary approach for analyzing brain activity patterns with behavior in the absence of behavioral data, we examined the spatial similarity of ReHo maps with 123 meta-analytic brain maps derived from the Neurosynth database. Neurosynth generates probabilistic measurements that can be interpreted as a quantitative representation of the association between regional fluctuations in activity and psychological processes. Although Neurosynth’s database contains over 1,000 terms, we narrowed our focus to terms related to mental processes and behavior, drawing from previous research (*91–93*). Our selected set of 123 terms encompasses broad categories, ranging from attention and emotion to more specific processes such as visual attention and episodic memory. The terms also include basic behaviors (e.g., eating and sleep) and emotional states (e.g., fear and anxiety).

### Statistical analysis

For participants with two sessions of neuroimaging sessions (either ACAM-J or control), metrics were averaged across sessions to reduce within-subject variability and simplify statistical modeling. Condition-level comparisons focused on a predefined mask derived from regions previously shown to be sensitive to ACAM-J effects in our earlier case studies. This targeted approach increased statistical power by restricting analyses to brain areas with known relevance. Complementary whole-brain analyses are reported in the **Supplementary results**. Missing phenomenology, and meditative traits were imputed using multivariate imputation by chained equations (MICE). This method was selected due to its robustness in preserving the multivariate relationships among variables, and its ability to produce unbiased parameter estimates under the assumption of missing at random (MAR) (*94, 95*).

#### Trends in neuroimaging metrics across ACAM-J

Linear mixed models (LMM) was used to examine polynomial trends up to the 3^rd^ degree (cubic trend) across ACAM-J for each neuroimaging metric. We implemented orthogonal polynomial contrast coding to test for linear, quadratic, and cubic relationships between the neuroimaging metric and the progression through ACAM-J. This approach allows us to capture potential non-linear patterns that might emerge across the ACAM-J progression, as shown in our previous case studies (*14, 25, 27, 68*). The polynomial terms were centered to reduce multicollinearity. In models with non-significant higher-order trends (e.g., non-significant cubic trend), we reduced the polynomial term iteratively (e.g. including quadratic and linear, or linear trend only) to simplify the model to best characterize the relationship between ACAM-J progression and the neuroimaging metric. Participant was also included as a random intercept individual differences in baseline neuroimaging metrics. Significance of models and polynomial terms were corrected for multiple comparisons using false discovery rate (FDR)–adjusted q-values. Models were corrected for multiple comparisons using FDR across the number of parcellations within each neuroimaging metric. Additional details for each neuroimaging metric are presented in subsequent sections. Analyses were conducted in *R* (*96*) loading on R Studio v2022.12.0.353.20 (*97*).

### Comparisons between ACAM-J and non-meditative control conditions

We first assessed whether the two non-meditative control conditions (counting and memory) differed significantly. To do so, we fit LMMs with control condition as the predictor and participant as the random intercept. Pairwise contrasts between counting and memory were computed, with FDR correction applied across the models to control for multiple testing.

Unless stated otherwise, using this composite control as the reference level, we then applied LMM to compare ACAM-J against the control condition. Each model included ACAM-J as a categorical predictor and participant as the random intercept. Estimated marginal means and pairwise contrasts were computed for each ACAM-J versus control comparison. Contrast coefficients within each model were corrected for multiple comparisons using FDR, and an additional FDR correction was applied across all models to control for multiple testing.

We also conducted supplementary analyses comparing the same neuroimaging metrics between ACAM-J and the control conditions separately to ensure robustness in our analysis. Results for this set of analysis are reported in the **supplementary materials**.

### Multivariate brain-behavior relationships: Linking neuroimaging metrics with phenomenology, meditative traits, and cognition

Unless stated otherwise, we used partial least squares correlation (PLSC) analysis to investigate the relationships between neuroimaging metrics and three domains of behavioral measures: phenomenology, meditative traits, and cognition. PLSC is a multivariate technique that identifies latent variables (LVs) representing patterns of covariance between brain activity and behavioral data (*98*). Phenomenological experience during ACAM-J and non-meditative tasks include quality of ACAM-J (none for control tasks), stability of attention, width of attention, sensation of sights, sounds, physical sensations, and narrative thought stream. Measures of meditative traits include the five dimensions of FFMQ (observing, describing, acting with awareness, non-judging of inner experience, and non-reactivity to inner experience), CAMS-R, NAS-SF, ES-16, two subscales of ACS (attentional focusing and attentional shifting), and IAI. Cognitive performance include all the 16 cognitive tasks as described previously.

For each analysis, we specified the number of components equal to the number of behavioral variables in the respective domain. Statistical significance of each latent variable was assessed using a 10,000 permutations to generate null distributions of singular values (*99*), with p_FDR_ < 0.05 considered significant (*100–102*). Bootstrap resampling with 10,000 iterations was used to estimate the reliability of brain and behavioral weights. The Kaiser criterion was applied to determine component significance, retaining components that explained more variance than the average across all components. Pearson’s correlations were conducted to examine the linear relationship between the latent X and Y variables.

For ReHo and principal connectivity gradients, PLSC was conducted at the ROI-level. For ReHo, we conducted PLSC using 498 ReHo ROIs, while PLSC was conducted using 400 cortical gradients ROIs for connectivity gradients. For geometric eigenmodes, we used global eigenmode activations (mean/max power/energy).

### Univariate brain-behavior relationships linking neuroimaging metrics with specific ACAM-J factors

Given that each ACAM-J has specific qualities, we also conducted Pearson’s correlation between the neuroimaging metrics and ACAM-J factors. In total, four sets of correlations were conducted within each neuroimaging metric: Bliss/joy (ACAM-J1-2), contentment (ACAM-J3), equanimity (ACAM-J4), and formlessness (ACAM-J5). Analyses were corrected for multiple testing with FDR within the specific set of analysis, i.e., within all ROIs for each ACAM-J factor.

### Robustness Checks: Subgroup Analyses

To assess the robustness of our primary findings, we conducted a series of subgroup analyses designed to account for potential heterogeneity in meditative experience and style. Specifically, we performed a secondary analysis excluding two deep-level ACAM-J practitioners (resulting in N = 18), as different forms or depths of ACAM-J practice may be associated with distinct phenomenological and neurobiological profiles. This exclusion aimed to isolate the effects of intermediate-level practice and reduce potential confounds introduced by advanced practitioners. Full statistical results and neuroimaging data for this subgroup analysis are reported in the **supplementary materials**.

Additionally, we evaluated the spatial correspondence of neural activation patterns between two subtypes of intermediate ACAM-J practice: Intermediate-Sutta-style and Intermediate-Tranquil Wisdom Insight Meditation (TWIM). To do this, we performed a permutation-based correlation analysis. First, we generated significant delta-ReHo maps for two subgroups: Subgroup A, which included all participants (N = 20), and Subgroup B, which excluded three TWIM-style ACAM-J practitioners (N = 3). We then computed the correlations of ReHo values across all ACAM-J between these two subgroups.

To assess the statistical robustness of the observed correlations, we conducted a permutation test with 10,000 iterations. This analysis allowed us to determine whether the inclusion or exclusion of TWIM-style practitioners significantly influenced the observed ReHo patterns. A high degree of correlation would indicate that TWIM and Sutta-style ACAM-J practices produce comparable neural signatures, suggesting a shared underlying phenomenological structure.

Finally, we conducted a simple linear regression analysis to examine whether spatial similarity between the two subgroups increased progressively across ACAM-J, thereby evaluating the stability and convergence of the neural signatures over ACAM-J.

### Contextualization of brain activity with Neurosynth psychobehavioral affinity maps

We used the *continuous.CorrelationDecoder* function from the NiMARE python package to contextualization of brain activity with Neurosynth psychobehavioral affinity maps (*103*). We first fetched Neurosynth dataset and converted it to NiMARE dataset. Following, we retrieved the 123 psychobehavioral terms from the database. We also computed delta-ReHo maps for every ACAM-J pair with the control composite (e.g., ACAM-J1-control, ACAM-J2-control) for decoding. Finally, we used the *CorrelationDecoder* function to decode the association between the unthresholded delta-ReHo maps and psychobehavioral affinity maps.

## Acknowledgements

We are grateful for the advanced meditators whose meditative capacities made this study possible. We also respectfully acknowledge the Buddhist traditions that have preserved and transmitted *jhana* meditation for more than two thousand years. The present study builds upon this historical knowledge by examining these advanced absorptive practices with contemporary scientific methods. Our aim here is not to uncritically validate religious claims, but rather to investigate such practices, and their outcomes, as reproducible human capacities that promise to contribute to human flourishing and well-being in the modern world.

## Declaration of generative AI and AI-assisted technologies in the writing process

ChatGPT (OpenAI, 2025) was used for copyediting and to help improve the clarity of this text. The authors reviewed and edited all AI-generated suggestions to ensure accuracy and appropriateness. All authors take full responsibility for the publication.

## Author contributions

Conceptualization: MDS, TS

Methodology: MDS, TS, MB

Investigation: WFZY, RP, TJ

Formal analysis: WFZY, GM, IB

Visualization: WFZY, GM, IB

Writing – original draft: WFZY, MDS

Writing – review & editing: WFZY, RP, MB, TJ, MDS

## Data and materials availability

Data may be requested and is subjected to the Massachusetts General Hospital Institutional Review Board’s guidelines and approval. All code used in this study may be made available by request from the corresponding contact email address.

## Supplementary Materials

### Supplementary Text

#### Dominant phenomenology of each ACAM-J

##### ACAM-J1

The dominant phenomenology of ACAM-J1 is one of joyful seclusion based on directed and sustained attention. Upon entering this state, the meditator experiences a distinct withdrawal from external sensations, internal monologue and narrative thoughts. The mind, with high stability and unification, is directed towards the meditation object with which it also experiences some degree of merger or absorption. This unified state is accompanied by five primary ACAM-J factors: directed attention, sustained attention, bliss, happiness, and one-pointedness of the mind. The most prominent feelings are the emergence of bliss and happiness. Bliss is an energetic, often physical joy, like thrilling waves, while happiness is a more serene, pervasive happiness. The meditator may still need to actively focus on the object and adjust the balance between effort and relaxation, so there is a quality of effortful engagement, but the overwhelming experience is seclusion, relief, delight, and the novel stability of a mind free from its usual mode of perception, planning, and thinking.

##### ACAM-J2

The dominant phenomenology of ACAM-J2 is bliss, which may be experienced as goosebumps, waves, or pervasive, intense pleasure. Directed attention, sustained attention falls away, as the mind no longer needs to actively fixate on the meditation object and adjust the balance between relaxation and effort. The mind now rests there immovably on its own to a much stronger degree. Bliss and happiness that emerges from concentration intensify and become the most prominent phenomenology. Bliss can feel more powerful and pervasive than in ACAM-J1, while the underlying happiness deepens into a profound contentment.

##### ACAM-J3

The dominant phenomenology of ACAM-J3 is mindful equanimity together with a deep happiness. The energetic and sometimes overwhelming waves of bliss have faded away entirely, seen now as a subtle distraction. Happiness that seems to permeate the entire body remains. It is a state of contentment and fulfilling happiness. The primary phenomenology is a gentle and refined, all-pervading physical and mental pleasure coupled with an unshakable mental balance.

##### ACAM-J4

The dominant phenomenology of ACAM-J4 is absolute neutrality. The subtle and refined happiness of ACAM-J3 is relinquished, along with any lingering trace of pain or dissatisfaction. The experience is defined by pure equanimity and one-pointedness. The mind is extremely still and unperturbed by internal or external stimulus. Consciousness is exceptionally bright and clear. ACAM-J4 serves as the foundation for the development of deep insight.

##### ACAM-J5

The dominant phenomenology of ACAM-J5 is the perception of boundless, limitless space. Attending to the concept “infinite space,” sets the mind free from all boundaries of location and physicality. The experience is one of profound expansion. The sense of bodily boundary and a physical space dissolves, replaced by a direct perception of space as an infinite, open field close to infinite awareness.

##### ACAM-J6

The dominant phenomenology is the direct experience of infinite and boundless consciousness. The object of meditation becomes the knowing quality of the mind itself, which is now seen to be as limitless as the space being perceived in ACAM-J5. The duality between the observer and the observed becomes blurred. It is a state of infinite field of knowing, aware of its own unbounded nature. The primary experience is one of luminous, self-aware knowing that has no center and no periphery.

##### ACAM-J7

The dominant phenomenology of ACAM-J7 is the perception of “nothingness” in which the mind let go of the object of infinite consciousness and takes “nothingness” as its focus. This is not a state of unconsciousness or a blank void, but a highly refined perception of absence where something could have been present. It is the most subtle form of object-based perception, where the mind clings to the very last concept of “something”, and in this case, the concept of “nothing”.

##### ACAM-J8

The dominant phenomenology of ACAM-J8 is so subtle that it is difficult to describe conceptually with words. The mind enters a state where consciousness cannot be said to be perceiving something, and yet it can also not be said to be perceiving a minimal something, like an absence. The mental factors are operating at their absolute minimal level. The mind is not cognizing any object, even the most refined, yet it is not unconscious. It is a state of residual mental formation, operating on the very edge of cessation.

#### Differing depths of ACAM-J and subgroup analyses

Different meditation traditions and schools have developed distinct interpretations and training methods for ACAM-J (*1–5*). Recent comparative work highlights three broad types of ACAM-J practice: light, intermediate, and deep (*1*). Intermediate ACAM-J emphasizes the cultivation of jhāna factors such as sustained effortless focus, joy, ease, and one-pointedness (*3, 6, 7*). These forms of ACAM-J are typically phenomenologically varied, permitting some thought-like activity, bodily awareness, and a more accessible entry into absorption. Teachers such as Leigh Brasington, Ayya Khema, and Bhante U Vimalaramsi represent this style. In contrast, light ACAM-J consists of less stable and more effortful focus, presence of narrative thought, and less pronounced presence of the ACAM-J factors both in terms of range and intensity. Deep ACAM-J, often associated with the *Visuddhimagga* (Path of purification in the Pali language) and teachers such as Pa Auk Sayadaw, is defined by absorption into a mental image or *nimitta* in the Pali language (*8, 9*). This form is phenomenologically stricter and entails very deep concentration in which bodily and sensory experience disappear entirely. Intermediate ACAM-J and deep ACAM-J differ most in absorption depth, the role of nimitta, and the extent of pacification of sensory input.

In our main analyses, we focus on the full sample of practitioners spanning these traditions (N=20), performing mostly intermediate ACAM-J, with a few cases of deep ACAM-J (N=2). We did not recruit ant light ACAM-J practitioners for this study. To assess robustness, however, we conducted a subgroup analysis which only includes intermediate ACAM-J (N=18). For the subgroup analysis, all control conditions were combined into a composite control, given minimal differences across them.

**Figure S1.**
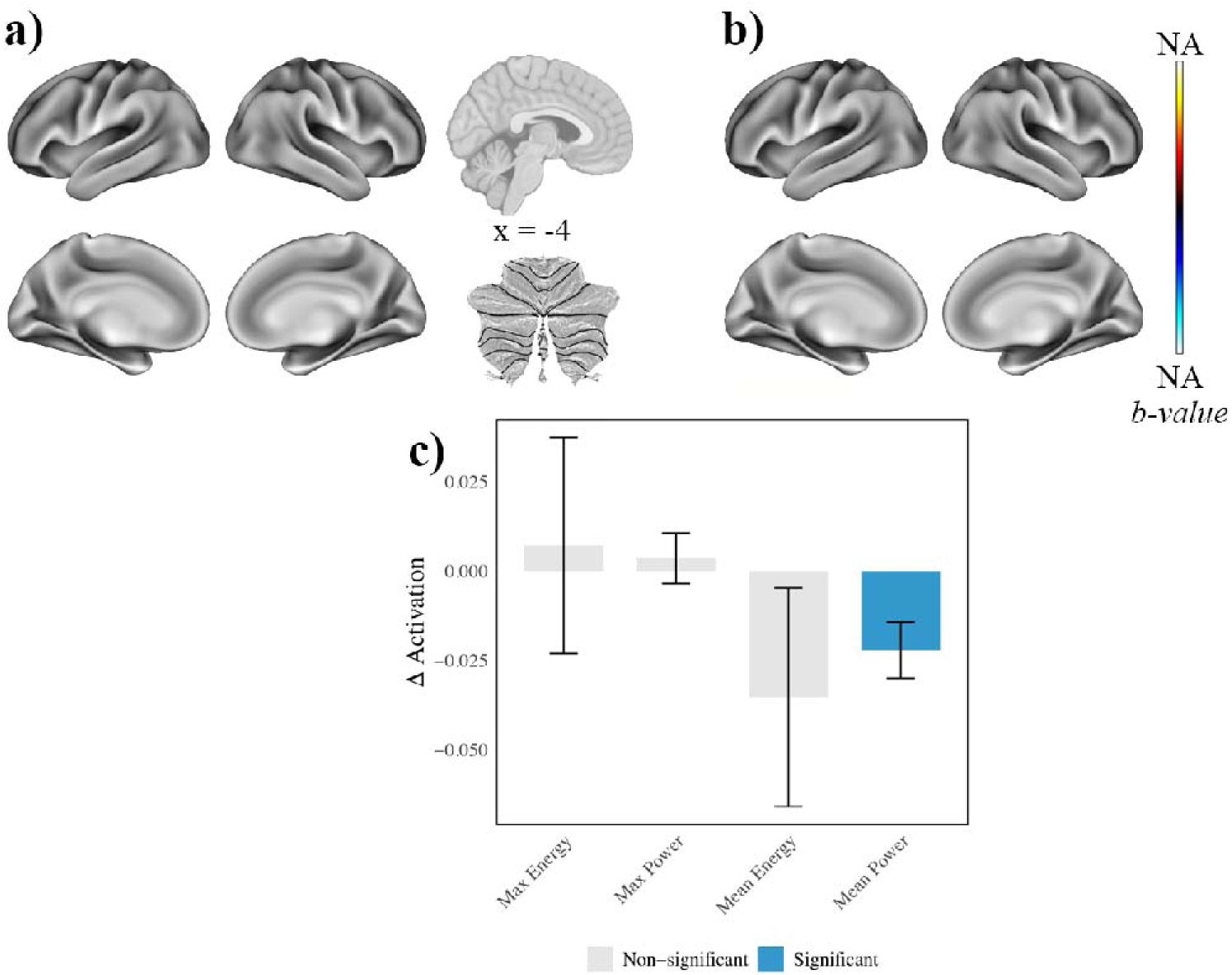
Comparison between memory and counting condition. (a-b) No significant differences in (a) ReHo and (b) G1 values were found between the two control conditions. (c) Mean power for the memory control condition was lower than the counting condition. Error bars represent standard errors.

**Figure S2.**
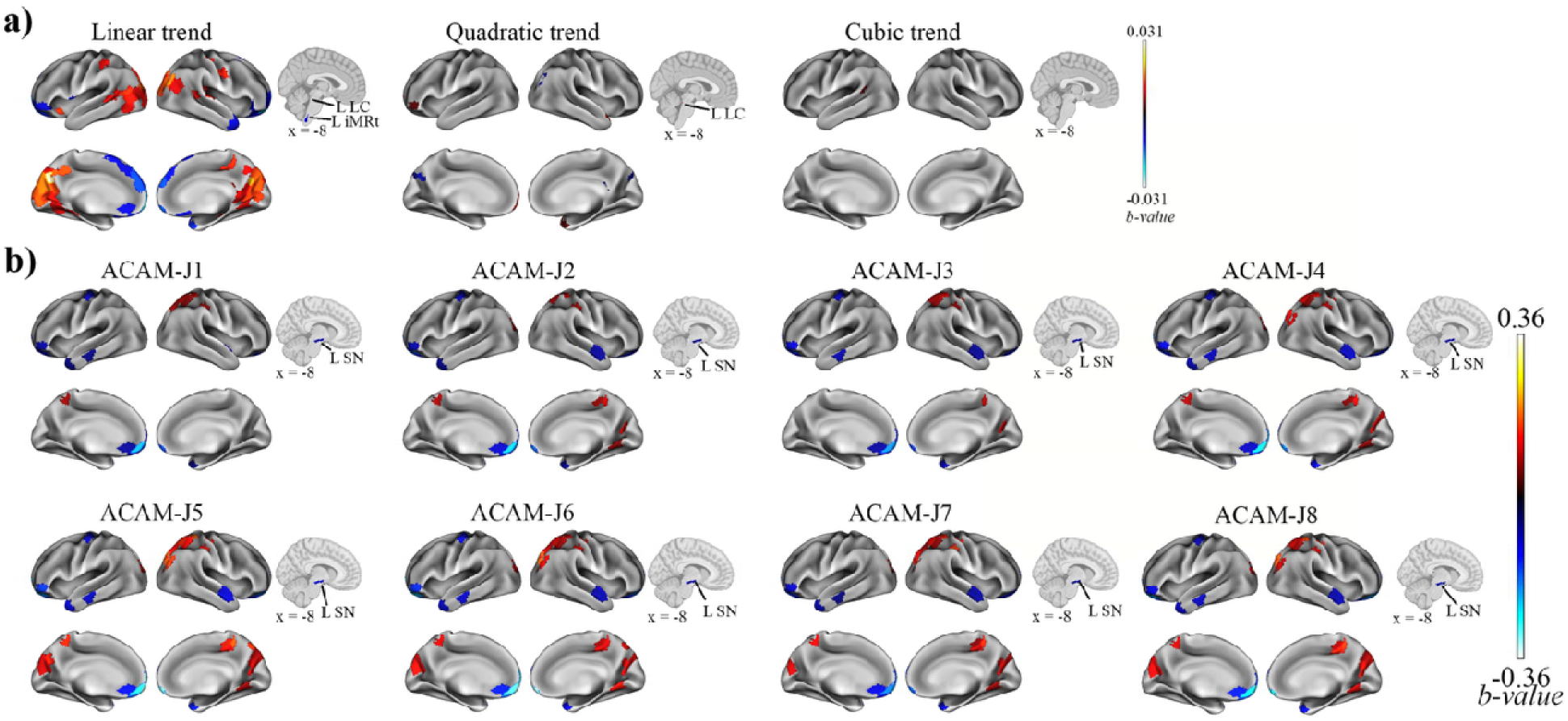
ReHo differences for ACAM-J vs the composite control condition (N=20) with multiple corrections across the entire brain (k = 498 ROIs). Trends analyses show regions exhibiting linear and quadratic trends across task ACAM-J. Linear trends highlights progressive and linear increase or decrease in specific brain regions while quadratic trends show mostly negative (inverted U-shaped) changes across ACAM-J. (b) ReHo differences between ACAM-J and the composite control reveal distinct patterns of cortical and brainstem substantia nigra activity.

**Figure S3.**
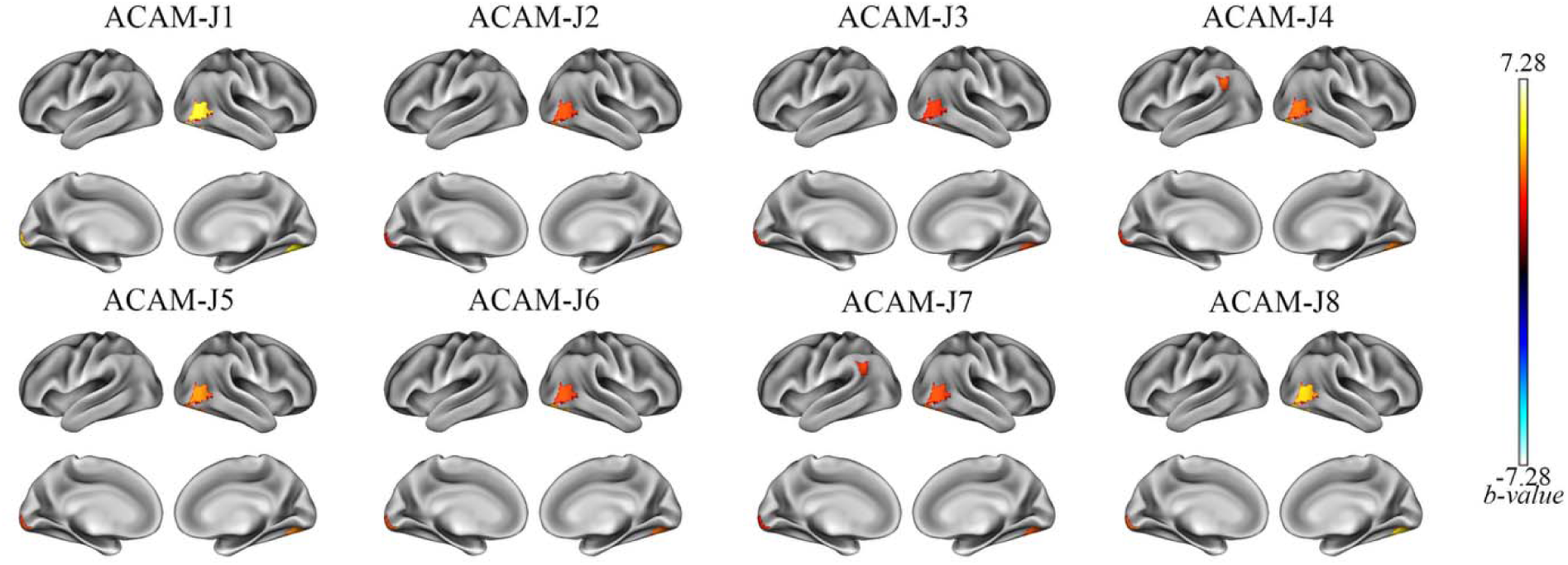
G1 differences for ACAM-J vs the composite control condition (N=20) with multiple corrections across the entire brain (k = 498 ROIs). No significant trends were found within ACAM-J. Contrasts between individual ACAM-J and the composite of control conditions reveal robust changes along G1 in visual and temporal hubs highlighting regional-specific reorganization of large-scale cortical hierarchy during ACAM-J.

**Figure S4.**
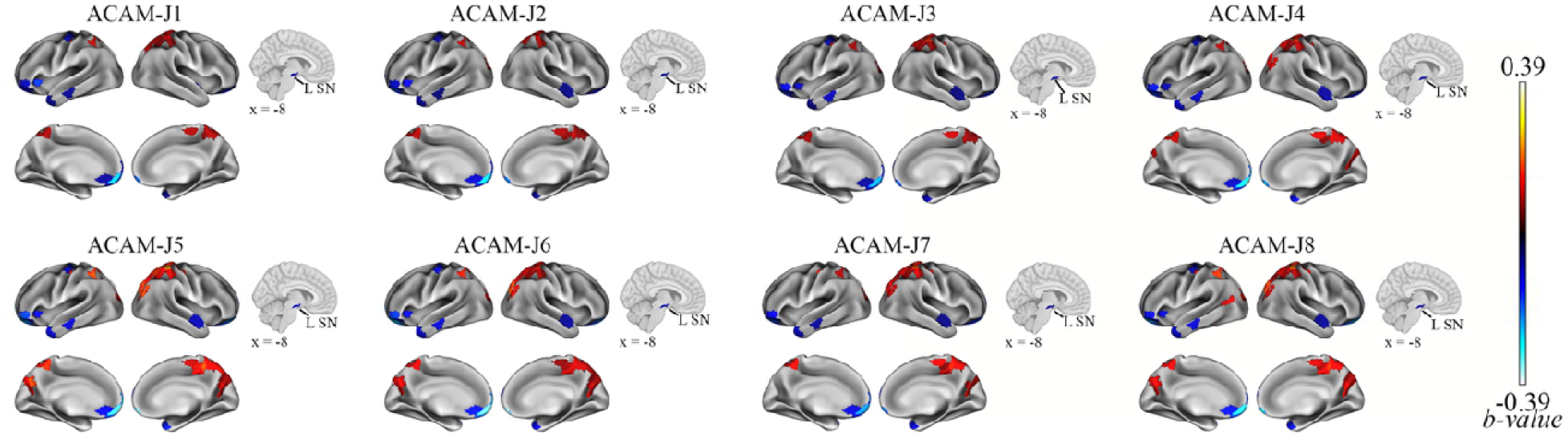
ReHo differences for ACAM-J vs counting condition in the full sample (N=20). Most ACAM-J were characterized by higher ReHo in the right posterior and somatomotor cortex and lower ReHo values in the bilateral orbitofrontal cortex (OFC), prefrontal cortex (PFC), and left substantia nigra. Later ACAM-J (ACAM-J5-8) had higher ReHo in the visual cortex.

**Figure S5.**
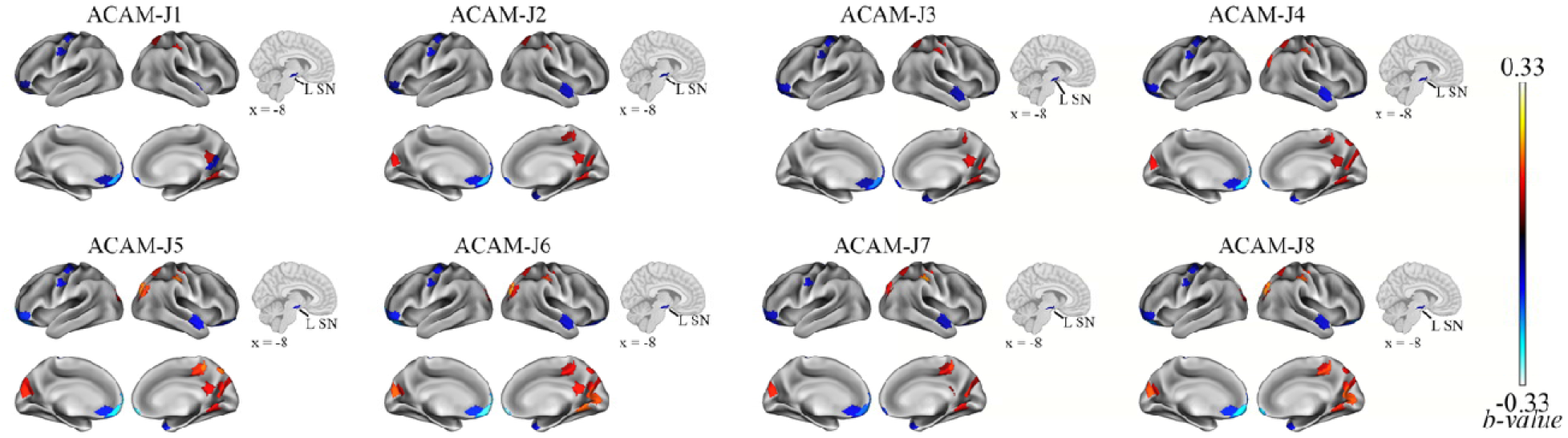
ReHo differences for ACAM-J vs memory condition in the full sample (N=20). Similar to **Figure S4**, most ACAM-J were characterized by higher ReHo in the right posterior and somatomotor cortex and lower ReHo values in the bilateral orbitofrontal cortex (OFC), prefrontal cortex (PFC), and left substantia nigra.

**Figure S6.**
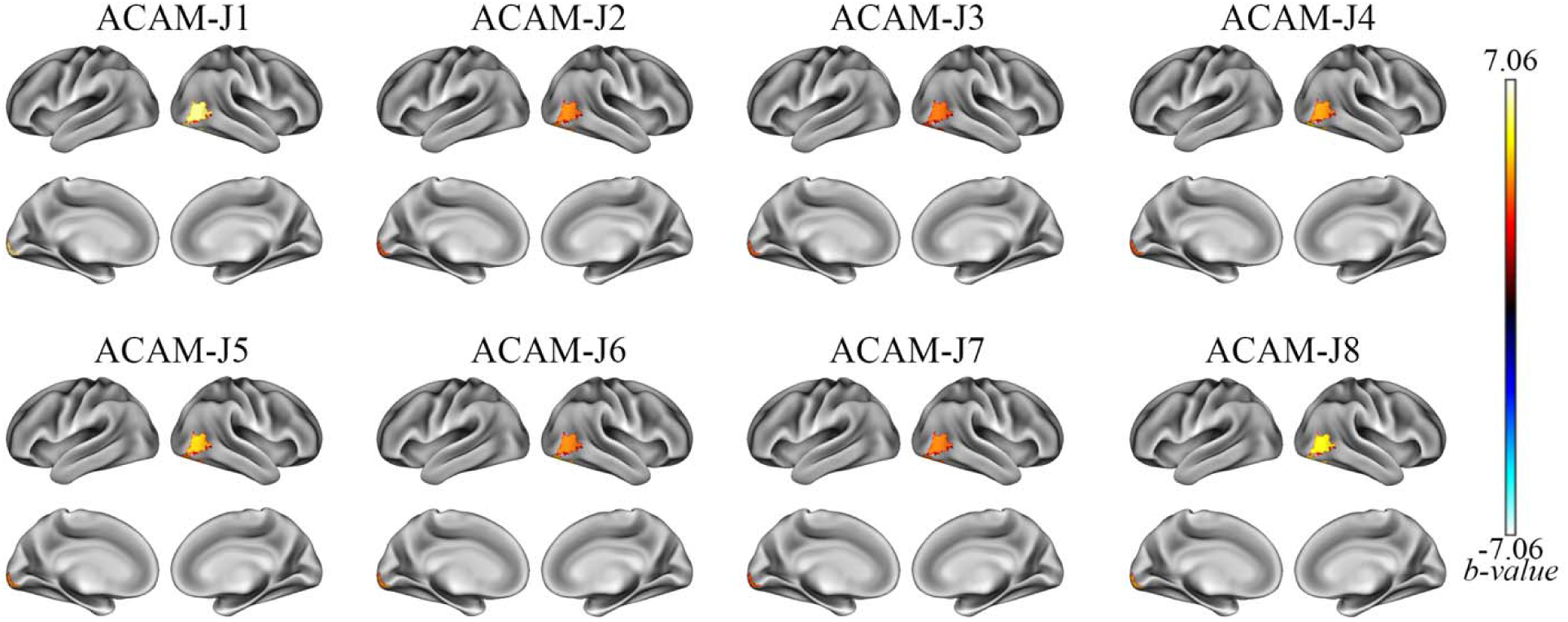
G1 differences for ACAM-J vs counting in the full sample (N=20). Relative to counting, most ACAM-J were characterized by higher G1 values in the visual cortex.

**Figure S7.**
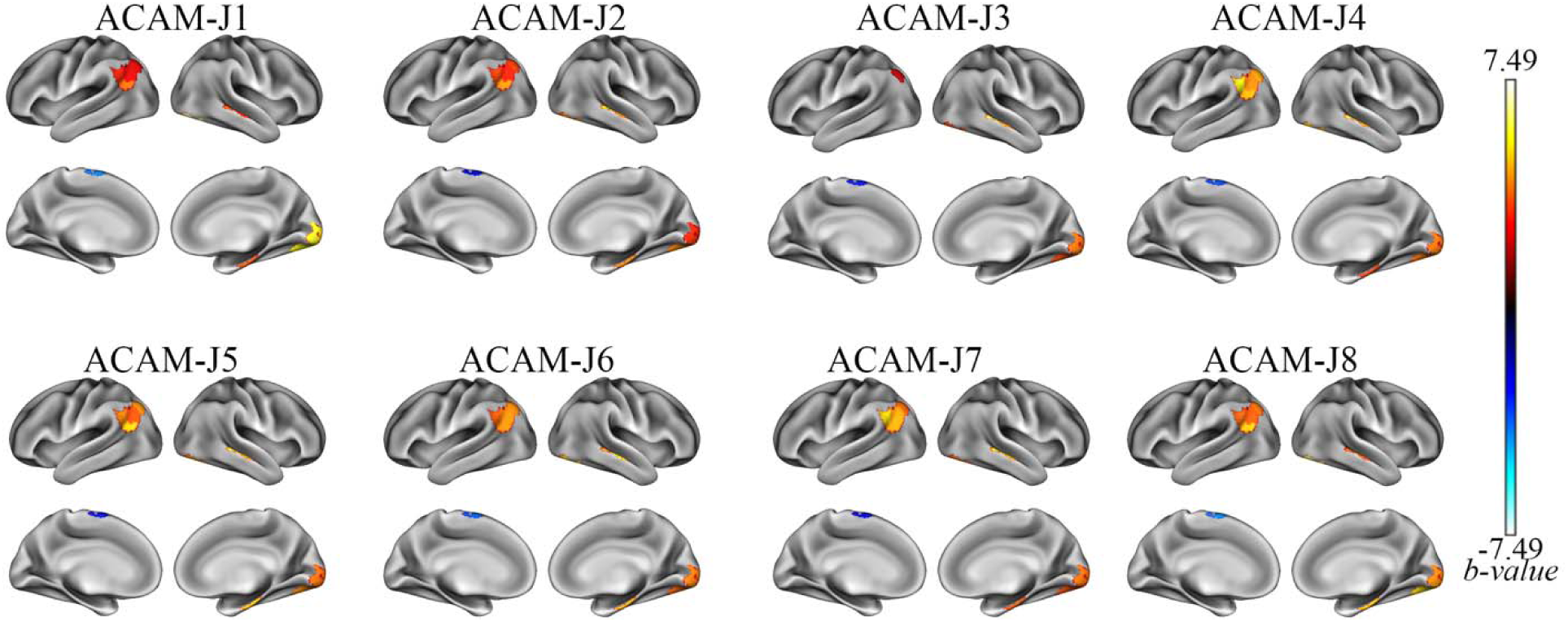
G1 differences for ACAM-J vs memory in the full sample (N=20). Compared to the memory control condition, most ACAM-J were characterized by higher G1 values in the visual cortex.

**Figure S8.**
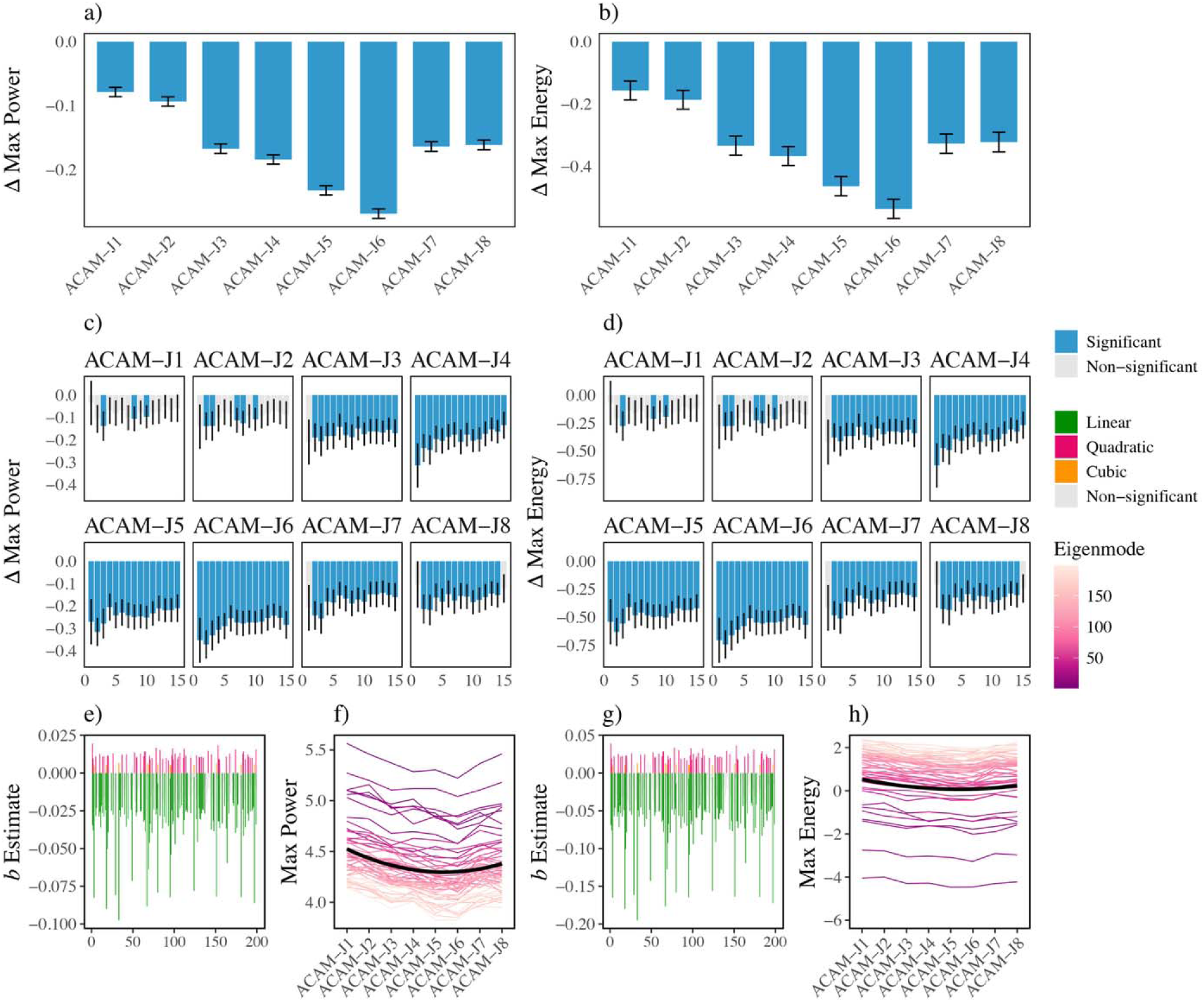
Global cortical dynamics in the full sample (N=20). (a-b) Relative to the control conditions, all ACAM-J showed lower max power/energy. (c-d) There were eigengroup nuances, with minimal significant differences for ACAM-J1 and ACAM-J2, and differences in most of the eigengroups from ACAM-J3 to ACAM-J8. (e-h) Both max power/energy exhibited similar negative linear and positive quadratic trends with mean power/energy, indicating an early decline from ACAM-J1 through ACAM-J5 followed by recovery in later ACAM-J. Error bars represent standard errors.

**Figure S8.**
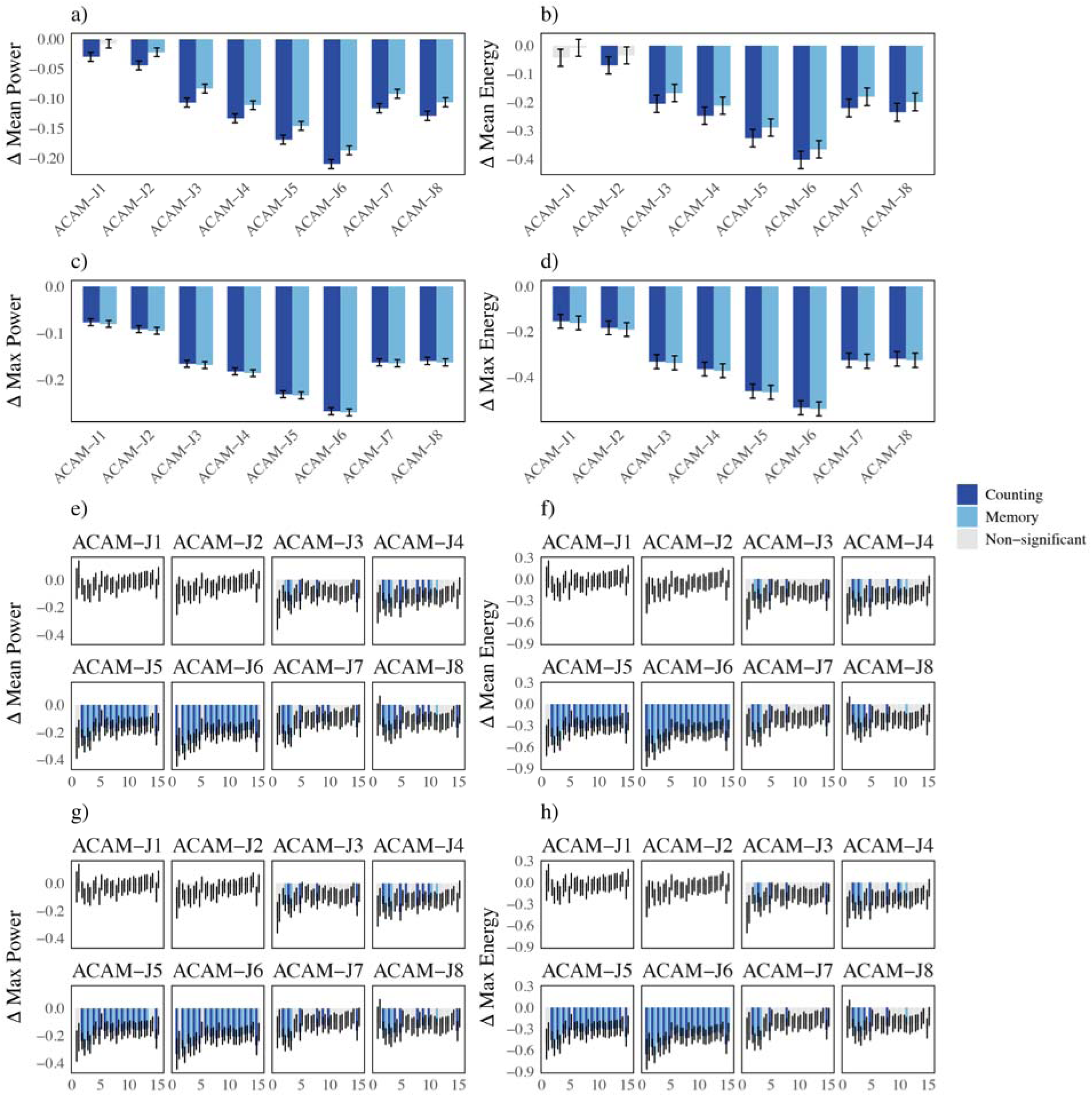
Global cortical dynamics compared to counting and memory control conditions (N=20). (a-d) Compared to all control conditions, most ACAM-J (except for ACAM-J1 and ACAM-J2) showed reduced (a-b) mean and (c-d) max power/energy. (e-h) There were subtle differences between ACAM-J, with minimal differences for ACAM-J1 and ACAM-J2. (e-f) Mean power/energy showed the most notable differences in eigengroup activations during ACAM-J5-6. In contrast, (g-h) max power/energy revealed differences in nearly all eigengroups between ACAM-J3 and ACAM-J8 when compared to both control conditions.

**Figure S10.**
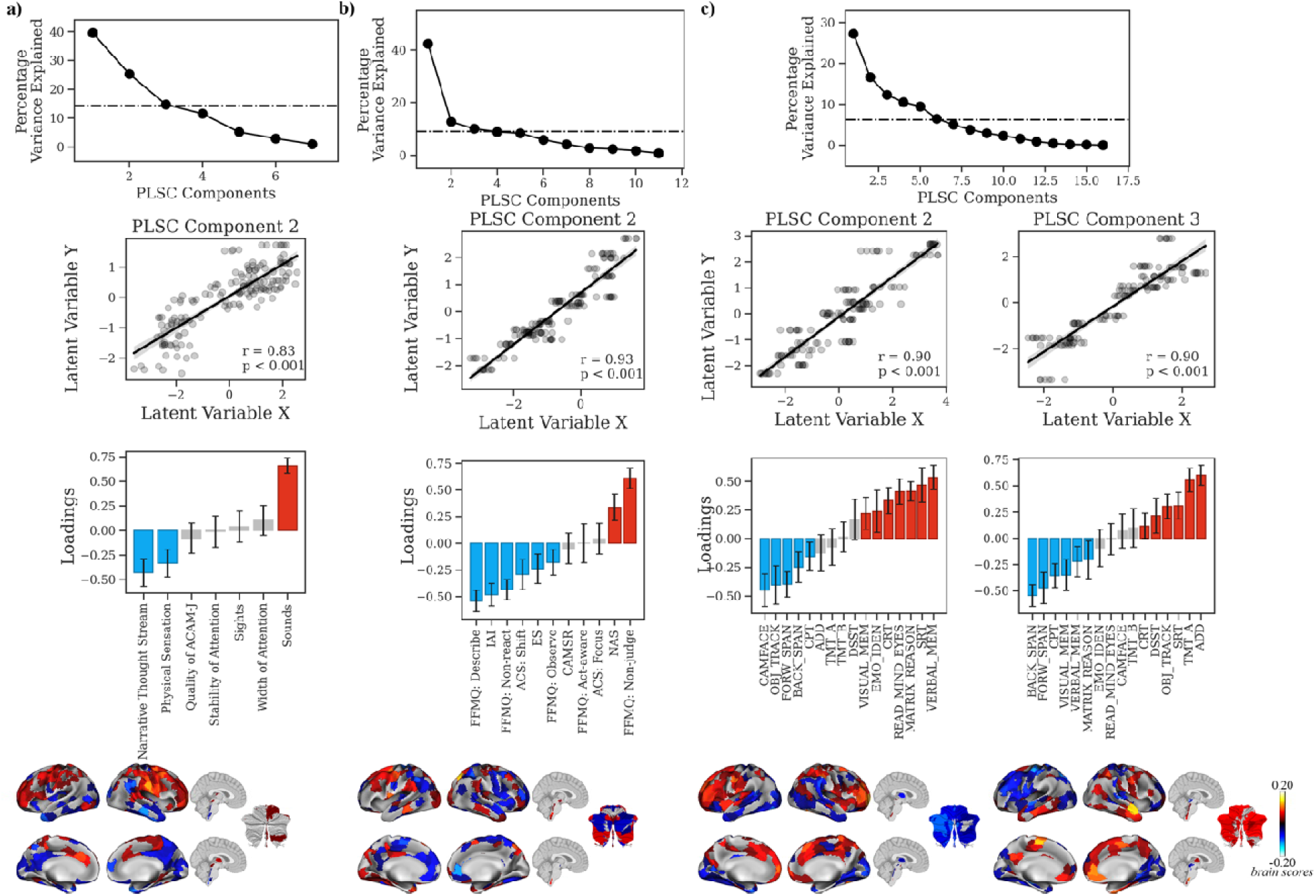
Additional multivariate relationship between ReHo during ACAM-J and behavioral patterns in the full sample (N=20). LV2 captured (a) ACAM-J phenomenology, with dorsal attention regions covaried with stability/width of attention, quality of ACAM-J, narrative thought stream, and physical sensations, and ventral sensory regions covaried with sounds; (b) meditative traits, with visuo-temporal, dorsolateral prefrontal cortex, and brainstem covaried with non-attachment/non-judgment and focus, and medial/lateral prefrontal regions aligned with general mindfulness; and (c) cognition, with PFC covaried with general cognition and visuo-temporal/basal ganglia covaried with working memory for latent variable 2 (LV2) and visual, prefrontal regions, brainstem, and cerebellum, covary with social function while lateral temporal and somatomotor regions covary with working memory functions. Error bars represent standard errors.

**Figure S11.**
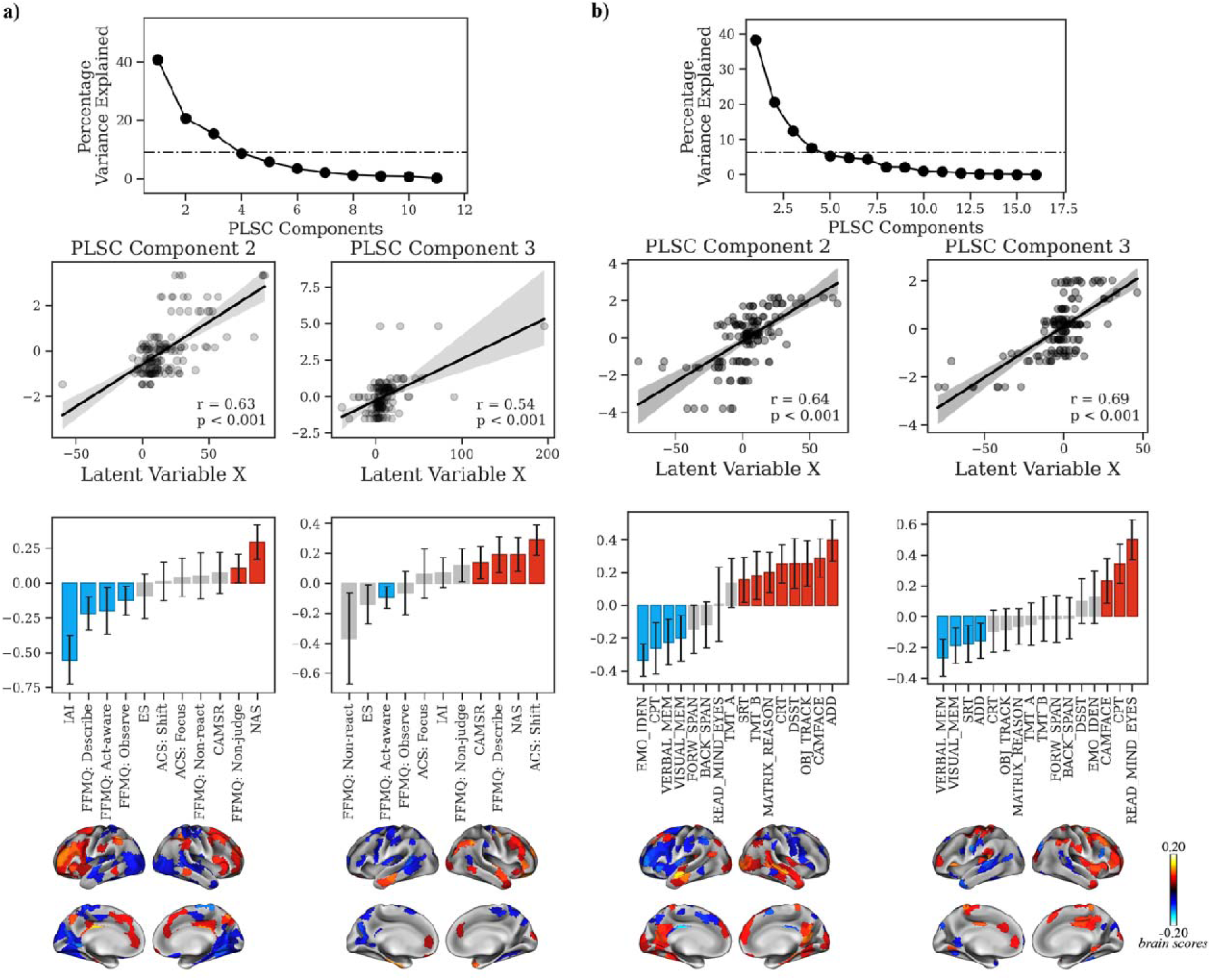
Multivariate relationship between G1 during ACAM-J and behavioral patterns. During ACAM-J, (a) For LV2 (left), visuo-temporal regions coavaried with nonattachment and dorsolateral PFC covaried with mindfulness. For LV 3 (right), visual, somatomotor, and posterior reigons covaried with awareness while frontotemporal regions covaried with mindfulness abilities (b) For cognition, LV2 showed covariance between general cognitive functions and visuo-temporal regions while working memory covaried with dorsolateral PFC. For LV 3, social function covaried with medial regions and PFC while working memory and sustained attention covaried with visual regions.

**Figure S12.**
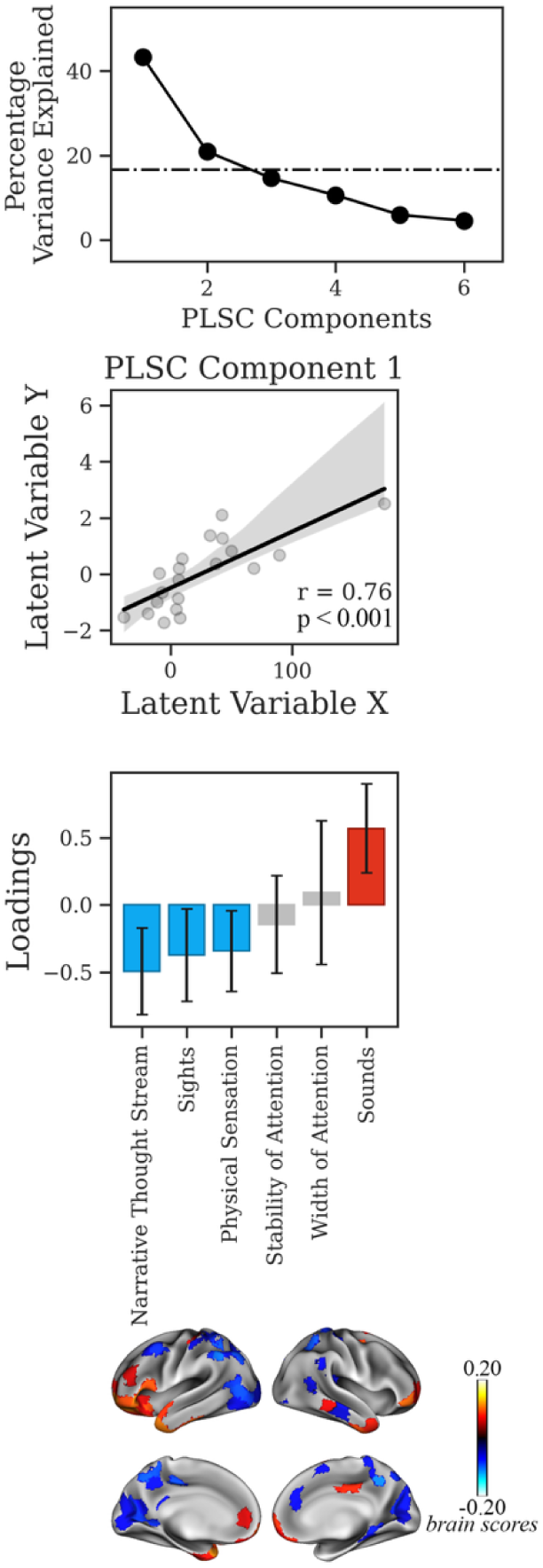
Multivariate relationship between G1 during control conditions and behavioral patterns. For the memory task, LV1 showed significant covariance between phenomenology and G1 values. Sensation of rounds covaried with G1 values in the PFC and temporal poles while other sensations (sights, physical sensations, narrative thought stream) covaried with G1 values in the visual and posterior regions.

**Figure S13.**
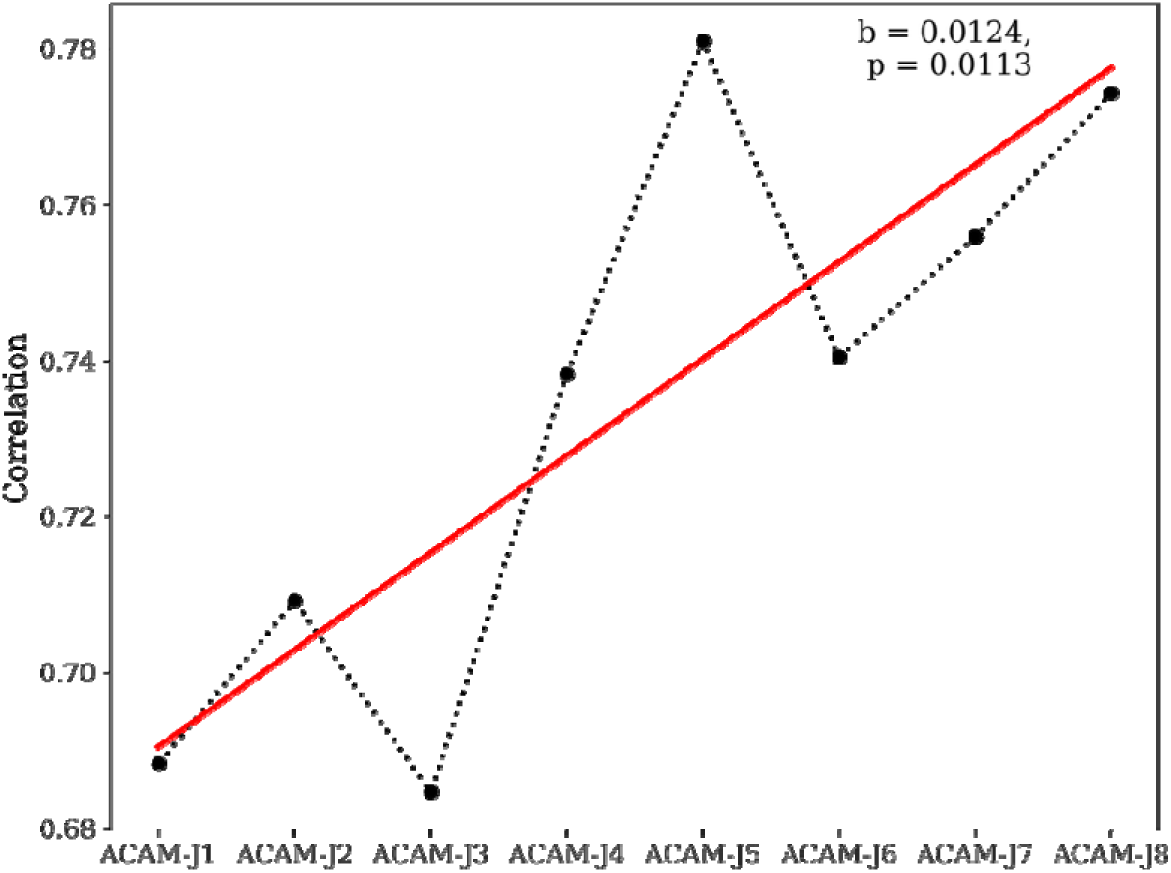
Spatial similarity between delta-ReHo maps of Sutta-style and TWIM-style ACAM-J across ACAM-J. Each point represents the spatial correlation at a ACAM-J. The red line indicates the best-fit linear regression (slope = 0.0124, = 0.0113), suggesting a significant positive trend in neural pattern similarity as ACAM-J deepens. This supports the hypothesis that TWIM-style and Sutta-style ACAM-J converge on similar neural signatures over the progression of meditative absorption.

**Figure S14.**
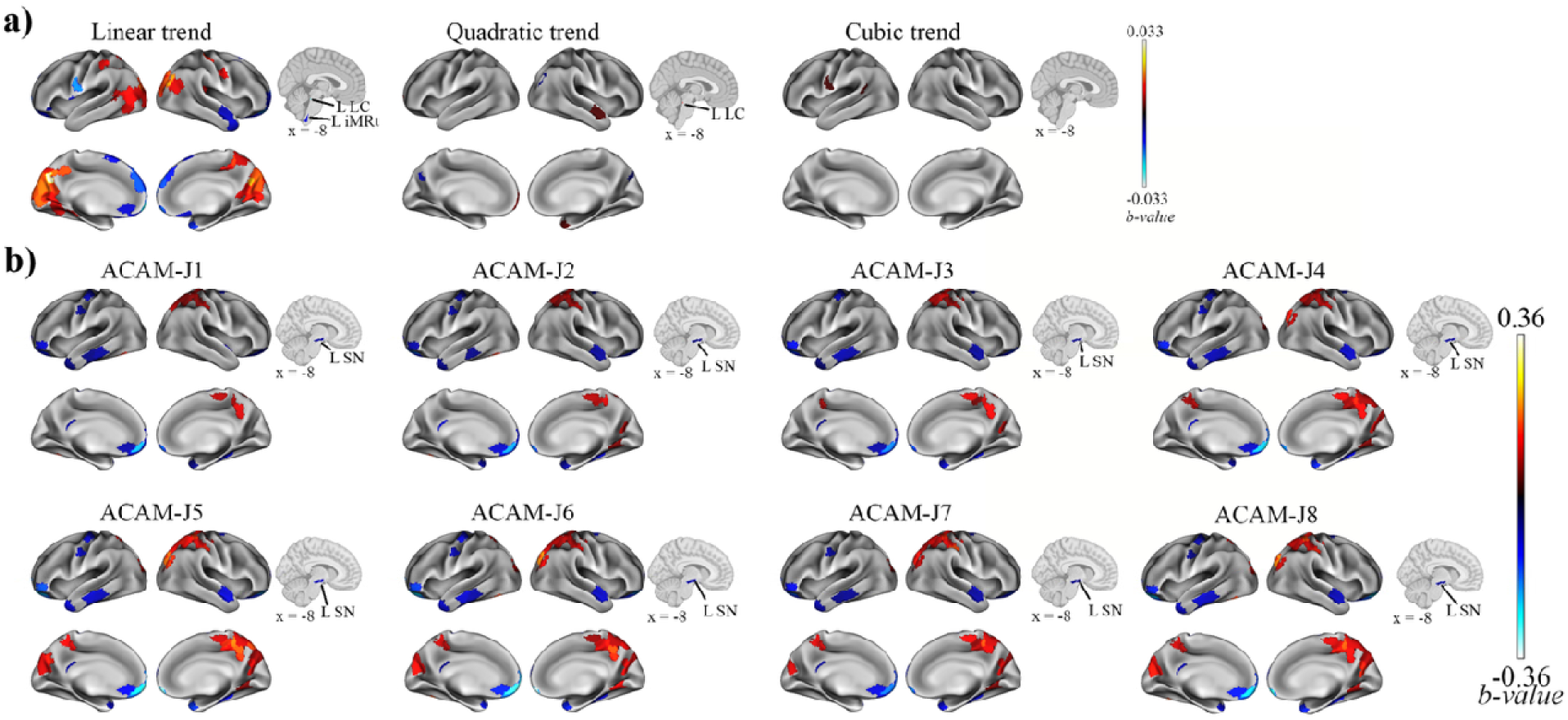
Distinct ReHo patterns intermediate ACAM-J. Excluding two deep ACAM-J practitioners did not influence the ReHo patterns in the results. (a) Polynomial trend analysis showing brain regions exhibiting significant linear, quadratic, and cubic changes in ReHo across ACAM-J. A significant linear trend is observed in medial prefrontal and posterior visual regions, suggesting progressive changes in brain activity across ACAM-J. Although quadratic or cubic trends were detected in very few parcels, their influence is not as remarkable as those of the linear trends. (b) Each ACAM-J showed consistent spatial patterns, particularly in the lateral temporal, prefrontal, and parietal cortices. Blue indicates decreased ReHo, while red indicates increased ReHo relative to the control condition. These findings confirm that excluding deep-level ACAM-J practitioners did not qualitatively alter the observed ReHo trajectories, supporting the robustness of the main results within the intermediate-level cohort.

**Figure S15.**
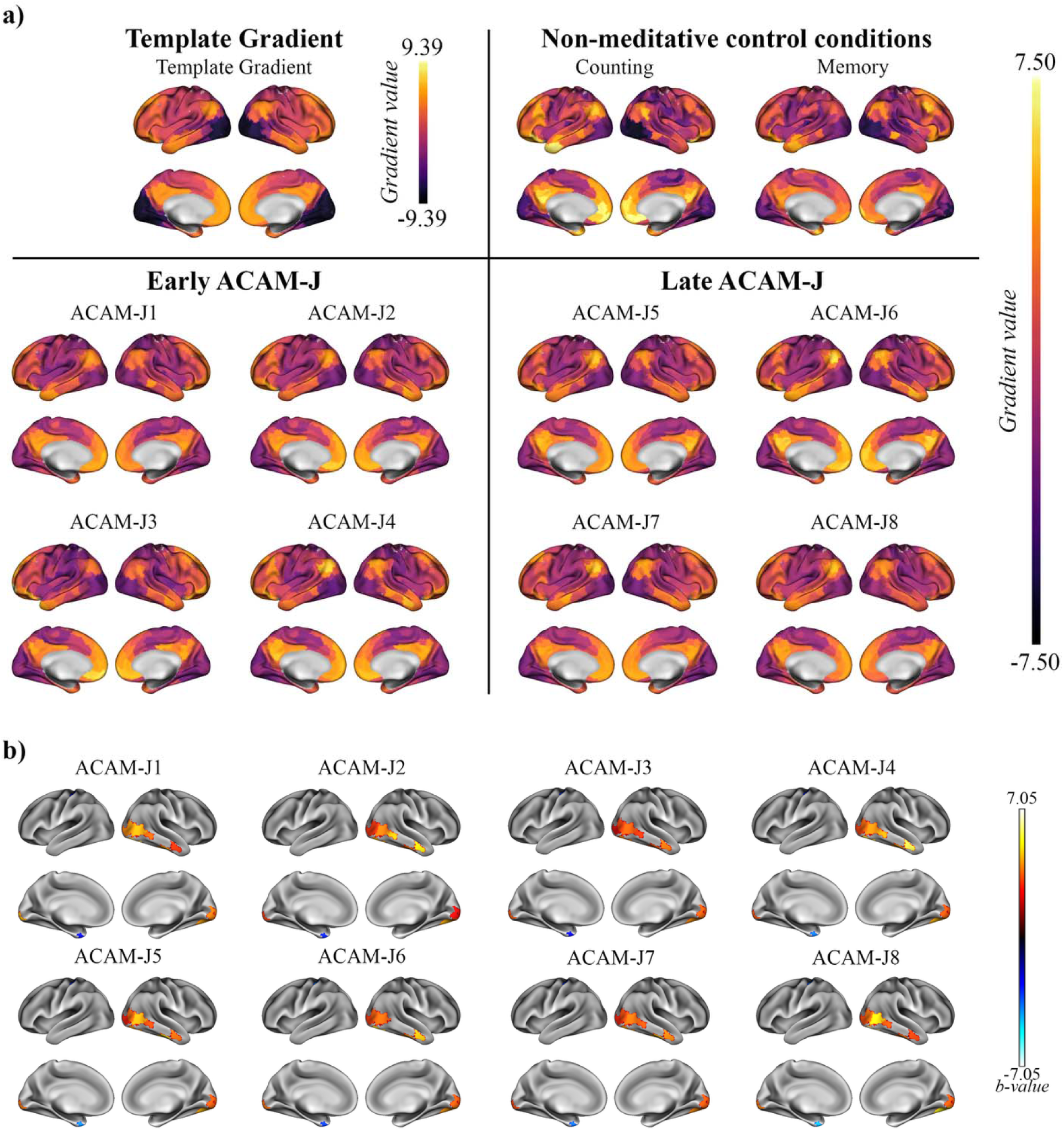
G1 dynamics across meditative and control conditions in intermediate ACAM-J practitioners (N = 18). (a) Template principal gradient (G1) used for gradient alignment in this study, and average gradient maps for the non-meditative control conditions (counting, memory) and ACAM-J, grouped into early and late absorption phases. (b) Contrasts between individual ACAM-J and the compositive control condition reveal robust changes along G1 in visual and temporal hubs highlighting regional-specific reorganization of large-scale cortical hierarchy during ACAM-J.

**Figure S16.**
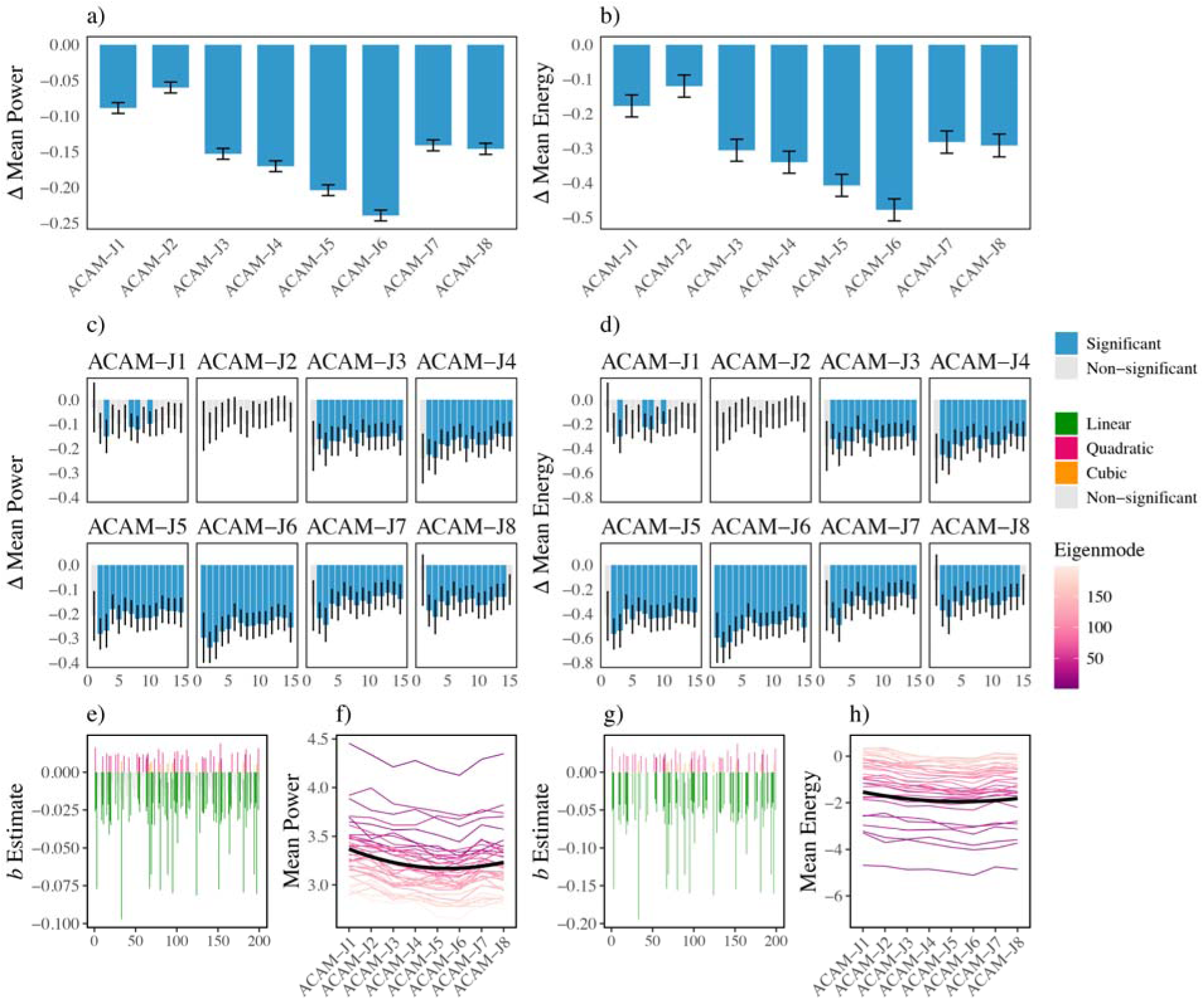
Geometric eigenmode decomposition during intermediate ACAM-J (N=18). (a–b) Overall mean/max power and energy reduced during ACAM-J compared with controls, except for ACAM-J1. (c-d) Most eigengroup differences were found in ACAM-J5-6. (e-h) Polynomial trends revealed consistent negative linear and positive quadratic trajectories across ACAM-J.

**Figure S17.**
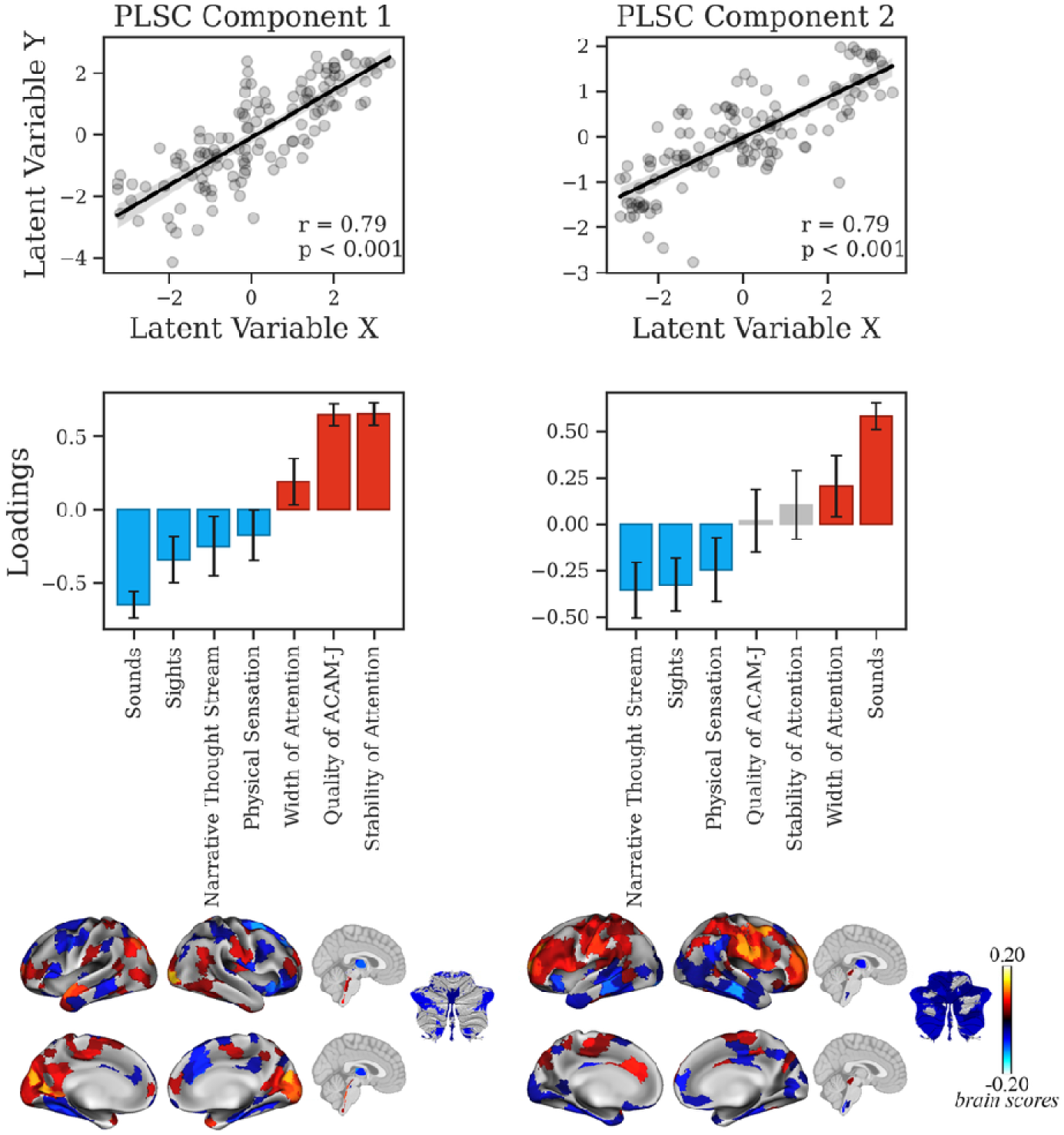
Multivariate relationship between ReHo and phenomenology during intermediate ACAM-J (N=18). PLSC identified two LVs linking ReHo with phenomenology. LV1 (left) primarily dissociated sensory and narrative processes (negative loadings: sounds, sights, narrative thought stream, physical sensation) from attentional qualities (positive loadings: width, stability of attention, quality of ACAM-J). In contrast, LV2 highlighted sensory salience, with sounds showing the strongest positive loading. Brain scores maps for LVs 1 and 2 reveal distributed cortical and subcortical systems covarying with phenomenological dimensions. Warm and cool colors represent positive and negative brain scores, respectively.

**Figure S18.**
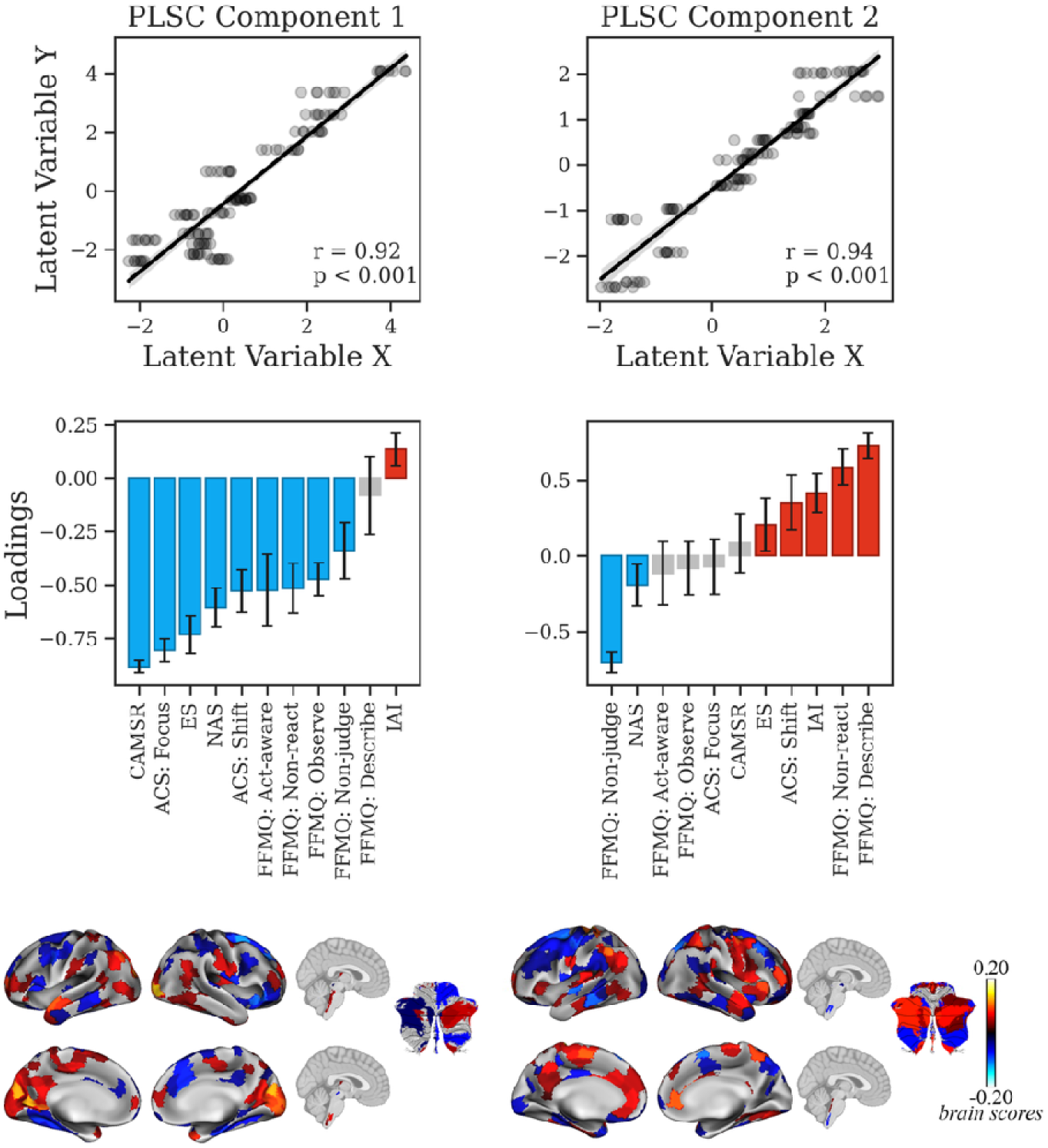
Multivariate relationship between ReHo and meditative traits during intermediate ACAM-J (N=18). PLSC identified two LVs linking ReHo with meditative traits. LV1 (left) primarily dissociated interoceptions (negative loadings: interoceptive awareness) from all other general mindfulness abilities (positrive loadings). In contrast, LV2 highlighted non-attachement (negative loadings) from mindfulness with equanimity (positive loadings). Brain scores maps for LVs 1 and 2 reveal distributed cortical and subcortical systems covarying with meditative traits. Warm and cool colors represent positive and negative brain scores, respectively.

**Figure S19.**
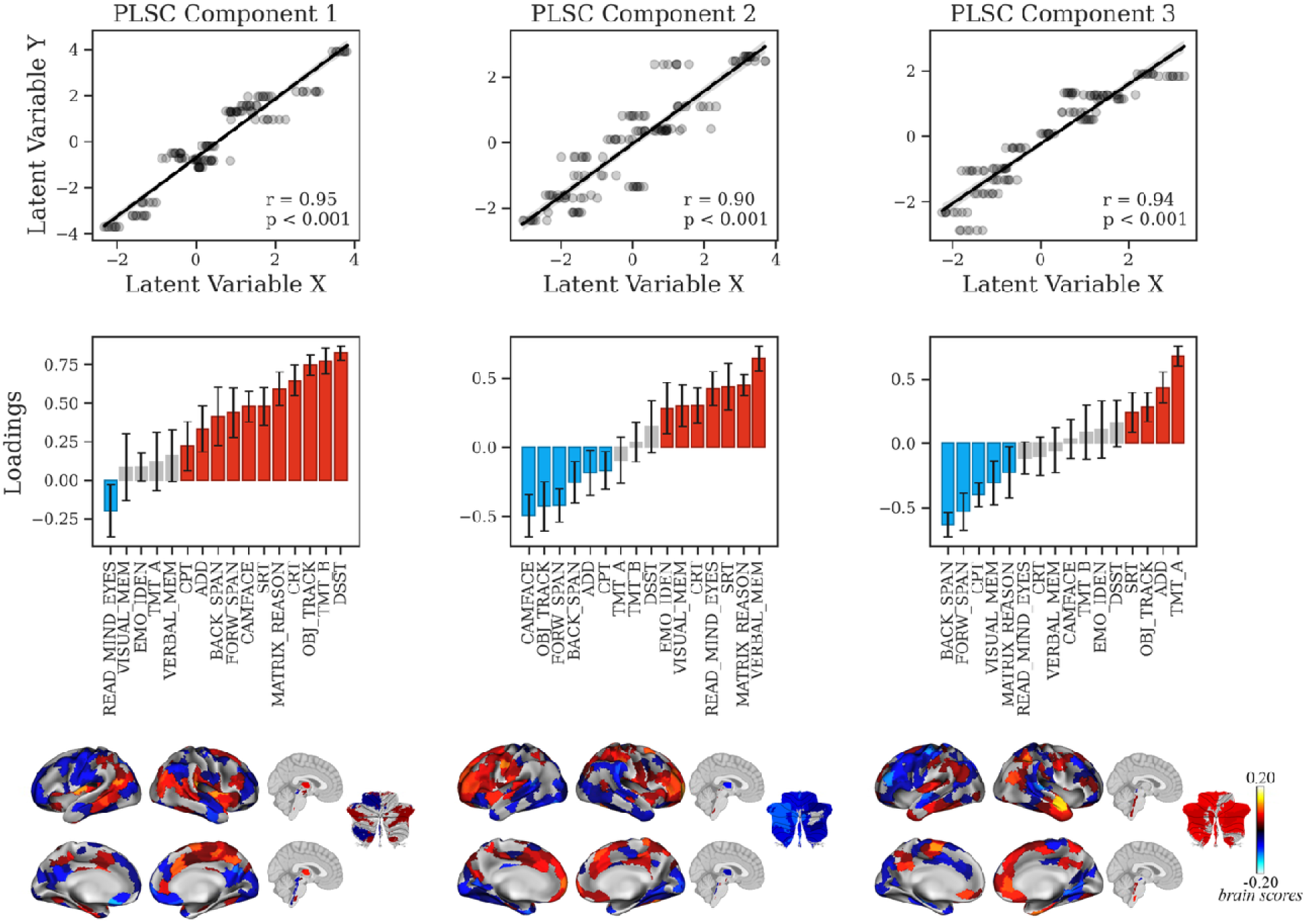
Multivariate relationship between ReHo and cognitive performance during intermediate ACAM-J (N=18). PLSC identified three LVs linking ReHo with cognitive performance. LV1 reflected a general cognitive proficiency axis, with strong positive loadings for most domains and negative loadings for social cognition (reading the mind in the eyes). LV2 dissociated socio-affective processing (negative loadings) from executive and attentional abilities (positive loadings). LV3 emphasized working memory and attentional control (positive loadings) relative to verbal memory and mentalizing abilities (negative loadings). Brain score maps for LVs 1 to 3 revealed distributed cortical and subcortical networks covarying with these distinct cognitive dimensions, with warm and cool colors indicating positive and negative brain scores, respectively.

**Figure S20.**
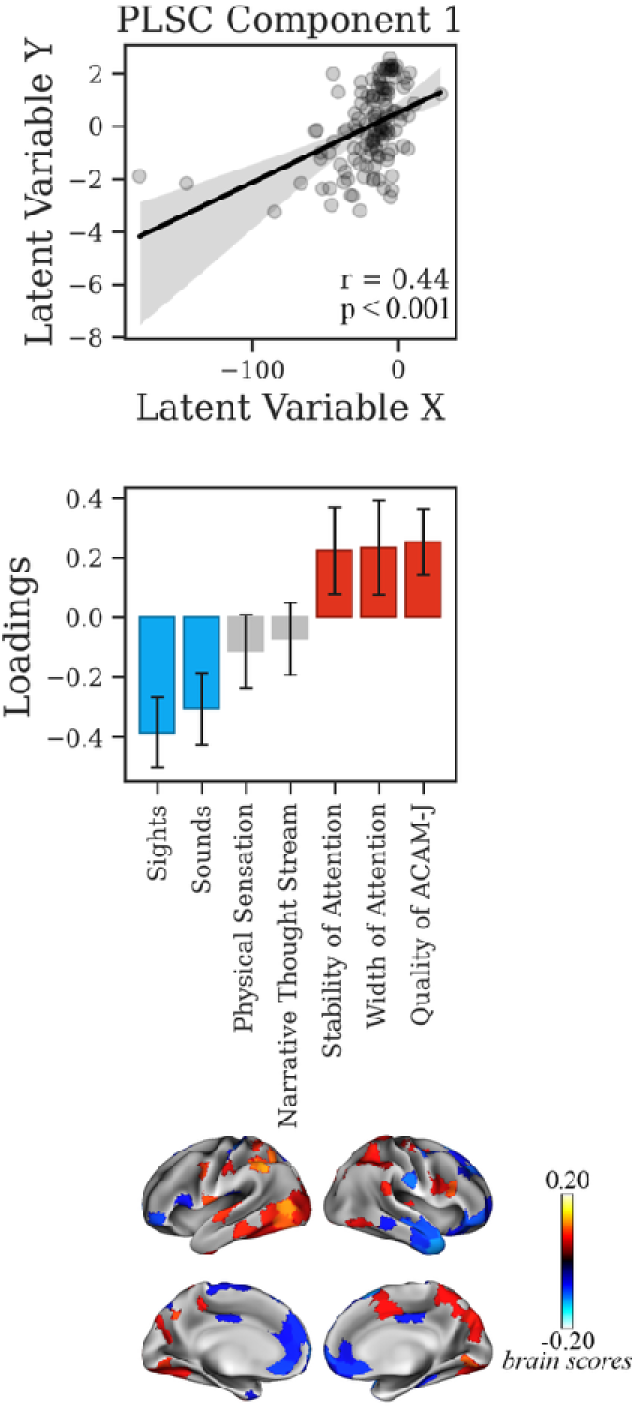
Multivariate relationship between G1 and phenomenology during intermediate ACAM-J (N=18). PLSC identified one LV linking G1 with phenomenology. Negative loadings were observed for attentional qualities (quality of ACAM-J, stability, width of attention), whereas positive loadings reflected sensory processes (sounds, sights,). Brain score maps revealed distributed cortical systems covarying with this phenomenological dissociation, with warm and cool colors representing positive and negative brain scores, respectively.

**Figure S21.**
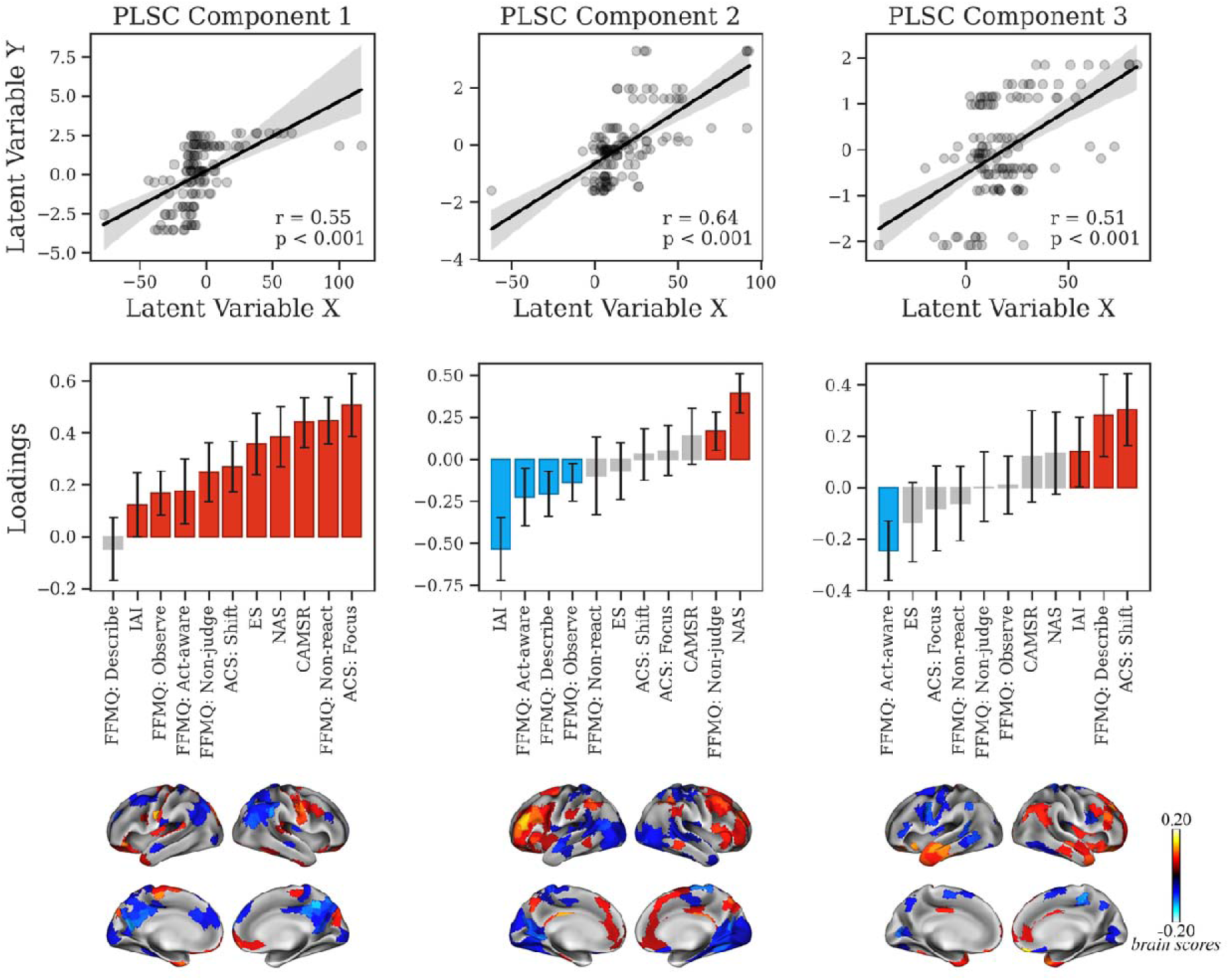
Multivariate relationship between G1 and meditative traits during intermediate ACAM-J (N=18). PLSC revealed three significant LVs linking large-scale cortical gradient organization with meditative traits. LV1 primarily reflected a broad positive association. LV2 dissociated mindfulness (negative loadings) from non-reactivity and non-attachment (positive loadings). LV3 further emphasized equanimity relative to descriptive mindfulness and attentional shifting (positive loadings). Brain score maps for LVs 1 to 3 revealed distributed cortical networks covarying with these trait dimensions, with warm and cool colors indicating positive and negative brain scores, respectively.

**Figure S22.**
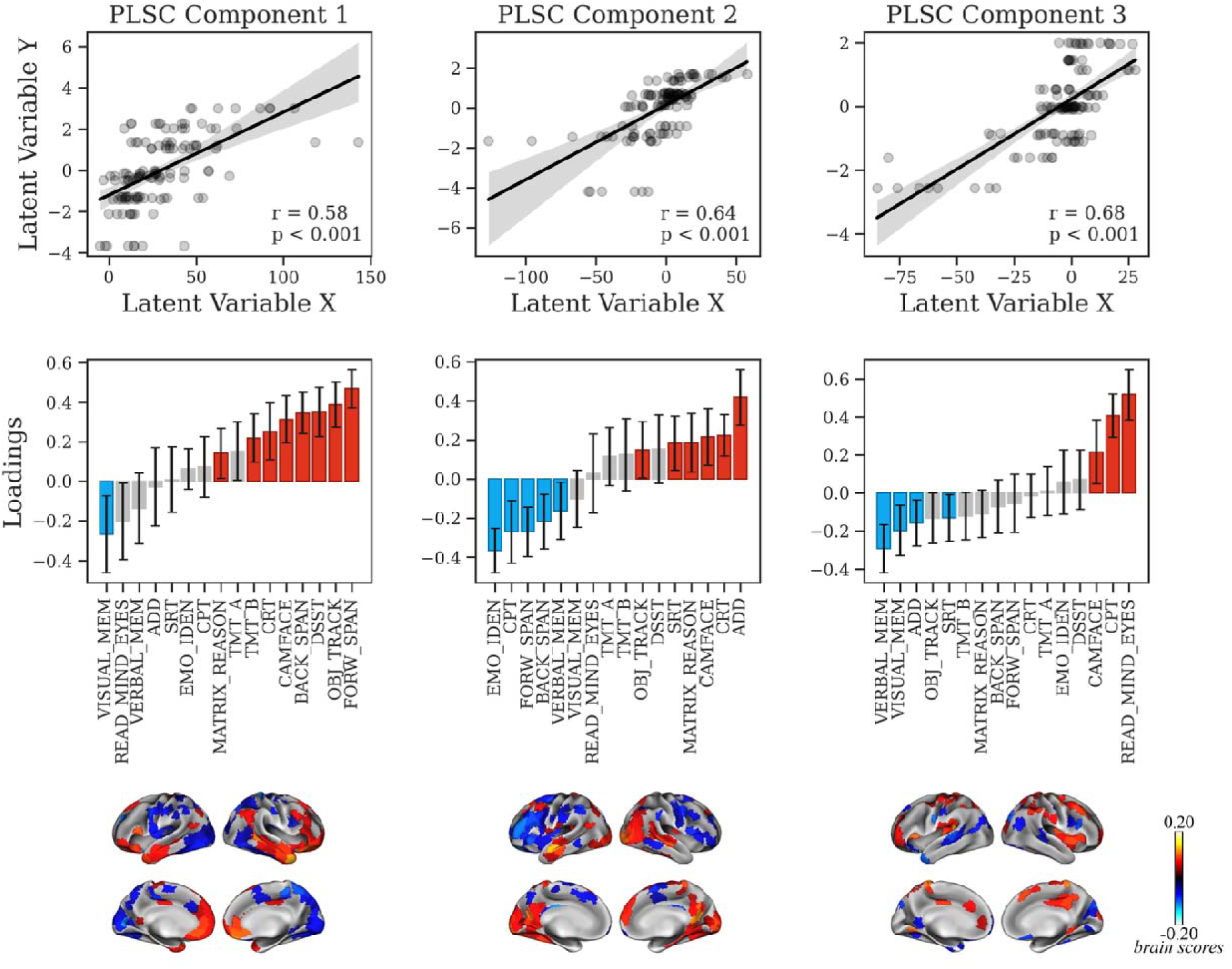
Multivariate relationship between G1 and cognition during intermediate ACAM-J (N=18). PLSC identified three significant LVs linking G1 with cognitive performance. LV1 reflected general cognitive perfofrmance, with strongest positive loadings for working memory, executive function, and attentional control, contrasted against contributions from verbal and visual memory (negative loadings). LV2 dissociated socio-affective processes (negative loadings) from executive reasoning and attentional measures (positive loadings). LV3 emphasized socio-cognitive and attentional skills (positive loadings) relative to verbal memory (negative loadings). Brain score maps for LVs 1 to 3 revealed distributed cortical and subcortical systems covarying with these cognitive dimensions, with warm and cool colors denoting positive and negative brain scores, respectively.

**Figure S23.**
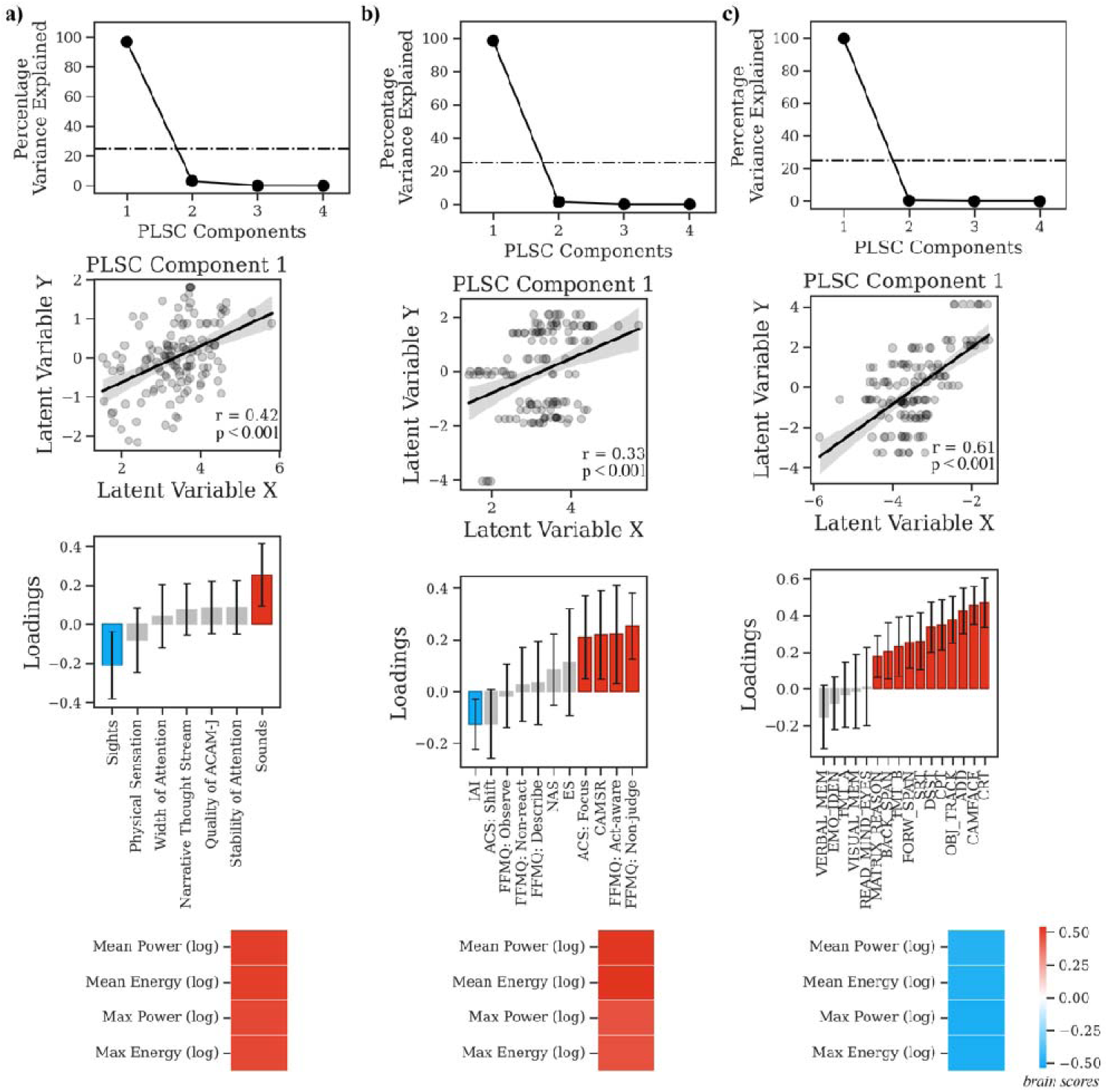
Multivariate relationship between global eigenmode activations and behavioral patterns during intermediate ACAM-J (N=18). PLSC revealed one significant LV in each domain linking eigenmode organization with (a) phenomenology, (b) meditative traits, and (c) cognitive performance. For phenomenology (a), the LV dissociated sensation of sights (negative loadings) from sounds (positive loadings). For meditative traits (b), the LV highlighted broad positive associations with trait mindfulness (positive loadings), contrasted against interoceptive awareness (negative loadings). For cognition (c), the LV captured a general cognitive performance axis, with positive loadings spanning working memory, executive function, and attentional control. Brain score maps indicated distributed cortical systems covarying with each domain, while bar plots show consistent positive relationships between eigenmode metrics (mean/max power and energy) and latent brain–behavior associations.

**Figure S24.**
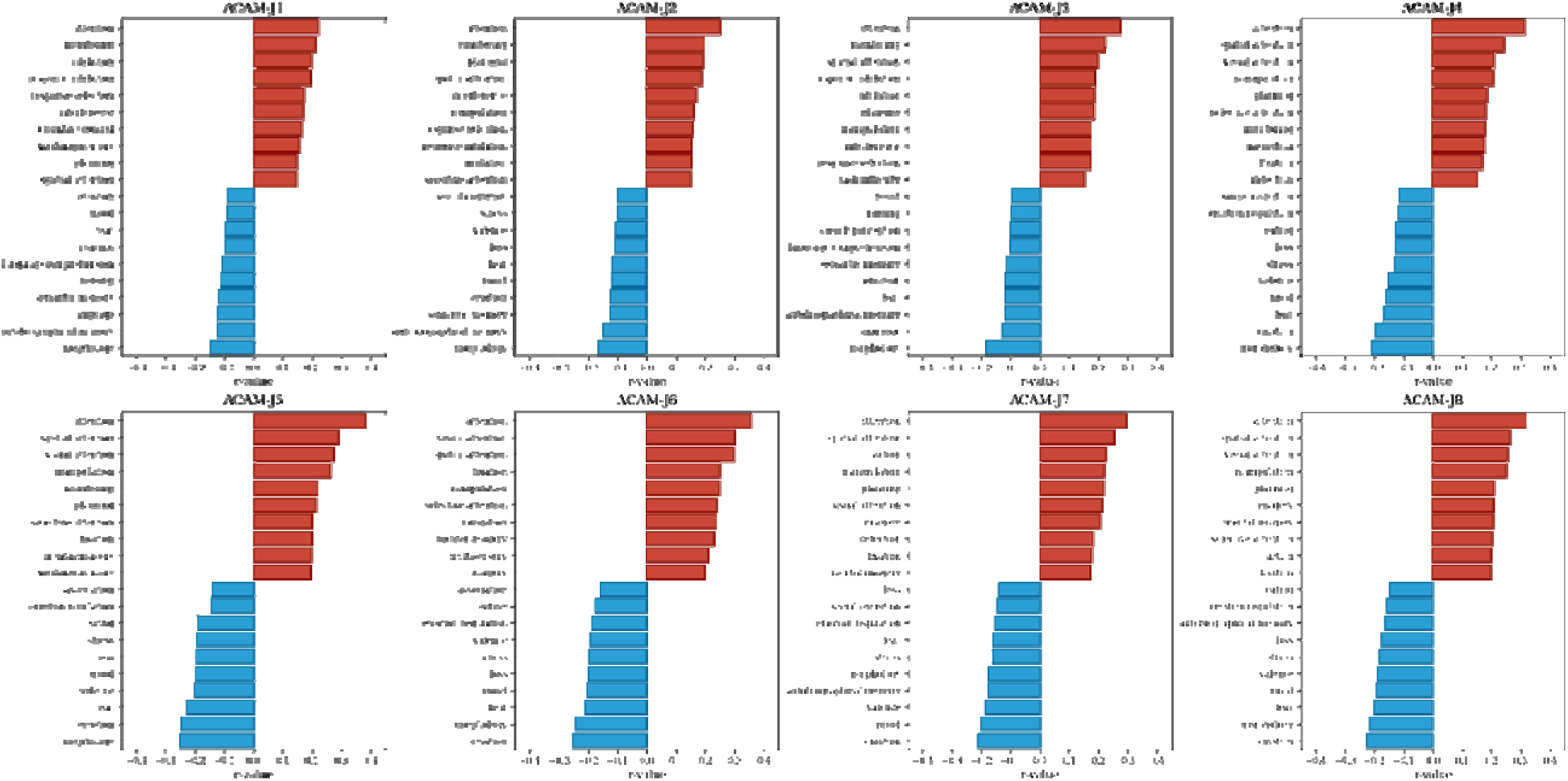
Reorganization of cognitive and affective domains during intermediate ACAM-J (N=18). For each ACAM-J, bar plots display the top ten functions with positive (red) and negative (blue) associations derived from Neurosynth decoding. Positive associations were dominated by attentional, executive, visuospatial, and action-related domains (e.g., sustained and selective attention, planning, imagery). In contrast, negative associations consistently involved affective, autobiographical, and bodily–social domains (e.g., emotion, mood, stress, fear, autobiographical memory, social cognition, eating, morphology).

**Table S1.** Average time spent in each ACAM-J after data cleaning.

| ACAM-J | Duration in minutes (M ± SD) |
| --- | --- |
| ACAM-J1 | 4.04 ± 4.29 |
| ACAM-J2 | 3.63 ± 1.38 |
| ACAM-J3 | 3.38 ± 1.04 |
| ACAM-J4 | 3.38 ± 1.24 |
| ACAM-J5 | 3.25 ± 1.47 |
| ACAM-J6 | 4.08 ± 4.69 |
| ACAM-J7 | 3.21 ± 1.38 |
| ACAM-J8 | 3.25 ± 1.63 |

**Table S2.** Description of cognitive tasks.

| **Task Name** | **Abbreviation** | **Description** |
| --- | --- | --- |
| Simple Reaction Time | SRT | Participant presses a key whenever a green square appears. Measures basic psychomotor response speed. |
| Choice Reaction Task | CRT | Participant indicates the direction of the arrow that is a different color from the rest. Measures processing speed, response selection/inhibition, and attention. |
| Digit Symbol Matching | DSST | Participant matches as many number/symbol pairs as possible in 90 seconds. Measures processing speed. |
| Trail Making Test Part A | TMT_A | Participant connects a series of circles with numbers in ascending order. Measures processing speed. |
| Trail Making Test Part B | TMT_B | Participant connects a series of circles with numbers and letters in alternating numerical and alphabetical order. Measures processing speed and task-switching (or cognitive flexibility). |
| Continuous Performance Test | CPT | Participant presses a button for each city image and withholds button presses for mountain images. Measures cognitive control, sustained attention, and response inhibition. |
| Paced Serial Addition Test | ADD | Participant adds sequentially presented pairs of numbers and judges whether their sum is less than or greater than 10. Measures sustained attention and working memory. |
| Forward Digit Span | FORW_SPAN | Participant is shown a series of numbers in increasing length and asked to recall the numbers in the order displayed. |
| Backward Digit Span | BACK_SPAN | asked participant is shown a series of numbers in increasing length and asked to recall the numbers in reverse order. |
| Multiple Object Tracking | OBJ_TRACK | Participant tracks a set of hidden dots across the screen. The dots increase in number and speed as the test progresses. Measures visuospatial attention and visual working memory. |
| Matrix Reasoning | MATRIX_REASON | Participant selects which smaller image completes the pattern in the larger image. Measures visual reasoning and general cognitive ability. |
| Verbal Paired Associates Memory | VERBAL_MEM | Participant memorizes a set of word pairs. Then, after completing TMB Digit Symbol Matching (90sec), the participant selects which words were paired together. |
| Visual Paired Associates Memory | VISUAL_MEM | Participant memorizes a set of picture pairs. Then, after completing TMB Digit Symbol Matching (90sec), the participant selects which pictures were paired together. |
| Cambridge Face Memory Test | CAMFACE | Participant memorizes a set of six faces. Then, the participant identifies the faces learned in the memorization phase from images where lighting and face angle vary. Measures unfamiliar face recognition. |
| Multiracial Emotion Identification | EMO_IDEN | Participant picks which of four basic emotions (happy, fearful, sad, or angry) best describes a face. Measures face emotion recognition. |
| Reading the Mind in the Eyes | READ_MIND_EYES | Participant judges which of four complex emotion words best describes a pair of eyes. Measures mental state inferencing and theory of mind. |

**Table S3.**
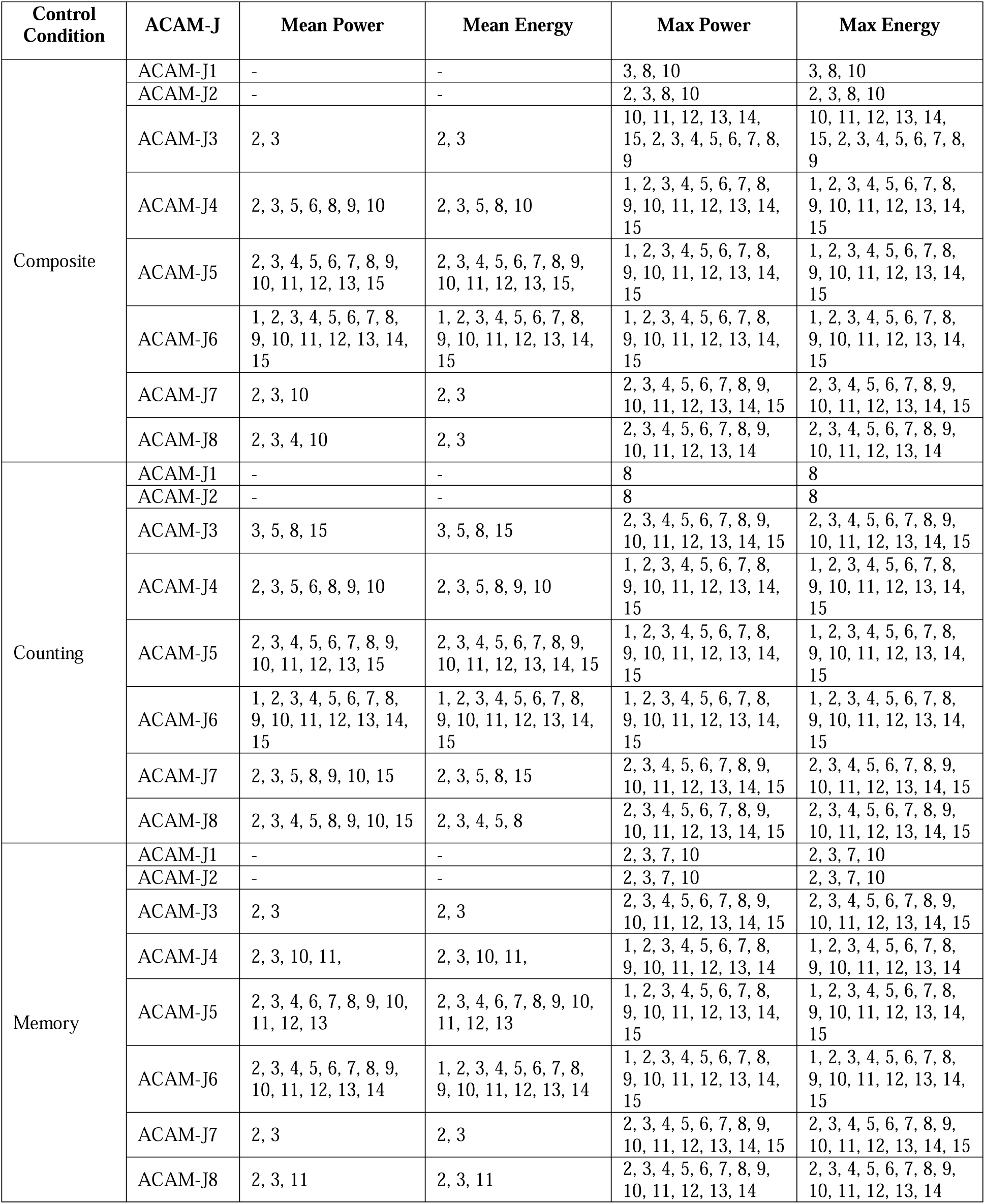
Significant mean/max eigenmode power/energy during ACAM-J compared to non-meditative control conditions in the full sample.

| Control Condition | ACAM-J | Mean Power | Mean Energy | Max Power | Max Energy |
| --- | --- | --- | --- | --- | --- |
| Composite | ACAM-J1 | - | - | 3, 8, 10 | 3, 8, 10 |
|  | ACAM-J2 | - | - | 2, 3, 8, 10 | 2, 3, 8, 10 |
|  | ACAM-J3 | 2, 3 | 2, 3 | 10, 11, 12, 13, 14, 15, 2, 3, 4, 5, 6, 7, 8, 9 | 10, 11, 12, 13, 14, 15, 2, 3, 4, 5, 6, 7, 8, 9 |
|  | ACAM-J4 | 2, 3, 5, 6, 8, 9, 10 | 2, 3, 5, 8, 10 | 1, 2, 3, 4, 5, 6, 7, 8, 9, 10, 11, 12, 13, 14, 15 | 1, 2, 3, 4, 5, 6, 7, 8, 9, 10, 11, 12, 13, 14, 15 |
|  | ACAM-J5 | 2, 3, 4, 5, 6, 7, 8, 9, 10, 11, 12, 13, 15 | 2, 3, 4, 5, 6, 7, 8, 9, 10, 11, 12, 13, 15, | 1, 2, 3, 4, 5, 6, 7, 8, 9, 10, 11, 12, 13, 14, 15 | 1, 2, 3, 4, 5, 6, 7, 8, 9, 10, 11, 12, 13, 14, 15 |
|  | ACAM-J6 | 1, 2, 3, 4, 5, 6, 7, 8, 9, 10, 11, 12, 13, 14, 15 | 1, 2, 3, 4, 5, 6, 7, 8, 9, 10, 11, 12, 13, 14, 15 | 1, 2, 3, 4, 5, 6, 7, 8, 9, 10, 11, 12, 13, 14, 15 | 1, 2, 3, 4, 5, 6, 7, 8, 9, 10, 11, 12, 13, 14, 15 |
|  | ACAM-J7 | 2, 3, 10 | 2, 3 | 2, 3, 4, 5, 6, 7, 8, 9, 10, 11, 12, 13, 14, 15 | 2, 3, 4, 5, 6, 7, 8, 9, 10, 11, 12, 13, 14, 15 |
|  | ACAM-J8 | 2, 3, 4, 10 | 2, 3 | 2, 3, 4, 5, 6, 7, 8, 9, 10, 11, 12, 13, 14 | 2, 3, 4, 5, 6, 7, 8, 9, 10, 11, 12, 13, 14 |
| Counting | ACAM-J1 | - | - | 8 | 8 |
|  | ACAM-J2 | - | - | 8 | 8 |
|  | ACAM-J3 | 3, 5, 8, 15 | 3, 5, 8, 15 | 2, 3, 4, 5, 6, 7, 8, 9, 10, 11, 12, 13, 14, 15 | 2, 3, 4, 5, 6, 7, 8, 9, 10, 11, 12, 13, 14, 15 |
|  | ACAM-J4 | 2, 3, 5, 6, 8, 9, 10 | 2, 3, 5, 8, 9, 10 | 1, 2, 3, 4, 5, 6, 7, 8, 9, 10, 11, 12, 13, 14, 15 | 1, 2, 3, 4, 5, 6, 7, 8, 9, 10, 11, 12, 13, 14, 15 |
|  | ACAM-J5 | 2, 3, 4, 5, 6, 7, 8, 9, 10, 11, 12, 13, 15 | 2, 3, 4, 5, 6, 7, 8, 9, 10, 11, 12, 13, 14, 15 | 1, 2, 3, 4, 5, 6, 7, 8, 9, 10, 11, 12, 13, 14, 15 | 1, 2, 3, 4, 5, 6, 7, 8, 9, 10, 11, 12, 13, 14, 15 |
|  | ACAM-J6 | 1, 2, 3, 4, 5, 6, 7, 8, 9, 10, 11, 12, 13, 14, 15 | 1, 2, 3, 4, 5, 6, 7, 8, 9, 10, 11, 12, 13, 14, 15 | 1, 2, 3, 4, 5, 6, 7, 8, 9, 10, 11, 12, 13, 14, 15 | 1, 2, 3, 4, 5, 6, 7, 8, 9, 10, 11, 12, 13, 14, 15 |
|  | ACAM-J7 | 2, 3, 5, 8, 9, 10, 15 | 2, 3, 5, 8, 15 | 2, 3, 4, 5, 6, 7, 8, 9, 10, 11, 12, 13, 14, 15 | 2, 3, 4, 5, 6, 7, 8, 9, 10, 11, 12, 13, 14, 15 |
|  | ACAM-J8 | 2, 3, 4, 5, 8, 9, 10, 15 | 2, 3, 4, 5, 8 | 2, 3, 4, 5, 6, 7, 8, 9, 10, 11, 12, 13, 14, 15 | 2, 3, 4, 5, 6, 7, 8, 9, 10, 11, 12, 13, 14, 15 |
| Memory | ACAM-J1 | - | - | 2, 3, 7, 10 | 2, 3, 7, 10 |
|  | ACAM-J2 | - | - | 2, 3, 7, 10 | 2, 3, 7, 10 |
|  | ACAM-J3 | 2, 3 | 2, 3 | 2, 3, 4, 5, 6, 7, 8, 9, 10, 11, 12, 13, 14, 15 | 2, 3, 4, 5, 6, 7, 8, 9, 10, 11, 12, 13, 14, 15 |
|  | ACAM-J4 | 2, 3, 10, 11, | 2, 3, 10, 11, | 1, 2, 3, 4, 5, 6, 7, 8, 9, 10, 11, 12, 13, 14 | 1, 2, 3, 4, 5, 6, 7, 8, 9, 10, 11, 12, 13, 14 |
|  | ACAM-J5 | 2, 3, 4, 6, 7, 8, 9, 10, 11, 12, 13 | 2, 3, 4, 6, 7, 8, 9, 10, 11, 12, 13 | 1, 2, 3, 4, 5, 6, 7, 8, 9, 10, 11, 12, 13, 14, 15 | 1, 2, 3, 4, 5, 6, 7, 8, 9, 10, 11, 12, 13, 14, 15 |
|  | ACAM-J6 | 2, 3, 4, 5, 6, 7, 8, 9, 10, 11, 12, 13, 14 | 1, 2, 3, 4, 5, 6, 7, 8, 9, 10, 11, 12, 13, 14 | 1, 2, 3, 4, 5, 6, 7, 8, 9, 10, 11, 12, 13, 14, 15 | 1, 2, 3, 4, 5, 6, 7, 8, 9, 10, 11, 12, 13, 14, 15 |
|  | ACAM-J7 | 2, 3 | 2, 3 | 2, 3, 4, 5, 6, 7, 8, 9, 10, 11, 12, 13, 14, 15 | 2, 3, 4, 5, 6, 7, 8, 9, 10, 11, 12, 13, 14, 15 |
|  | ACAM-J8 | 2, 3, 11 | 2, 3, 11 | 2, 3, 4, 5, 6, 7, 8, 9, 10, 11, 12, 13, 14 | 2, 3, 4, 5, 6, 7, 8, 9, 10, 11, 12, 13, 14 |

## References

1. M. J. Wright, J. L. Sanguinetti, S. Young, M. D. Sacchet, Uniting contemplative theory and scientific investigation: Toward a comprehensive model of the mind. Mindfulness 14, 1088–1101 (2023).

2. J. Galante et al., A framework for the empirical investigation of mindfulness meditative development. Mindfulness 14, 1054–1067 (2023).

3. K. Shinozuka, W. F. Z. Yang, R. M. Potash, T. Sparby, M. D. Sacchet. (Preprint), vol. 2025.

4. W. F. Z. Yang et al. (Preprint), vol. 2025.

5. A. Amaro, Realization Is Here and Now. Mindfulness 12, 795–797 (2020).

6. B. Anālayo, The Four Levels of Awakening. Mindfulness 12, 831–840 (2020).

7. K. Rose, Yoga, Meditation, and Mysticism: Contemplative Universals and Meditative Landmarks. (Bloomsbury Academic, New York, 2016).

8. L. N. Toan, Salvation, Enlightenment and Awakening in Shaping Followers’ Perception and Understanding of Their Faith: A Linguistic Exploration. International Journal of Religion 5, 286–299 (2024).

9. A. Berkovich-Ohana, A case study of a meditation-induced altered state: increased overall gamma synchronization. Phenomenology and the Cognitive Sciences 16, 91–106 (2015).

10. R. van Lutterveld, A. Chowdhury, D. M. Ingram, M. D. Sacchet, Neurophenomenological Investigation of Mindfulness Meditation “Cessation” Experiences Using EEG Network Analysis in an Intensively Sampled Adept Meditator. Brain Topogr, (2024).

11. R. van Lutterveld, T. Cahaly, D. Ingram, M. D. Sacchet, Brain criticality and advanced meditation “cessation” events: An intensively sampled electroencephalography case study. SSRN, (2025).

12. A. Chowdhury et al., Investigation of advanced mindfulness meditation “cessation” experiences using EEG spectral analysis in an intensively sampled case study. Neuropsychologia 190, 108694 (2023).

13. R. E. Laukkonen et al., Cessations of consciousness in meditation: Advancing a scientific understanding of nirodha samāpatti. Progress in Brain Research 280, 61–87 (2023).

14. W. F. Z. Yang et al., Intensive whole-brain 7T MRI case study of volitional control of brain activity in deep absorptive meditation states. Cerebral Cortex 34, bhad408 (2024).

15. T. Sparby, M. D. Sacchet, The Third Wave of Meditation and Mindfulness Research and Implications for Challenging Experiences: Negative Effects, Transformative Psychological Growth, and Forms of Happiness. Mindfulness, (2025).

16. T. Sparby, M. D. Sacchet, Toward a Unified Model of Advanced Meditation, Human Development, Meditation Maps, and Transtradition Metaphors: Facing Impermanence, Suffering, and Death. Mindfulness, (2025).

17. W. F. Z. Yang, T. Sparby, M. Wright, E. Kim, M. D. Sacchet, Volitional mental absorption in meditation: Toward a scientific understanding of advanced concentrative absorption meditation and the case of jhana. Heliyon 10, E31223 (2024).

18. T. Sparby, M. D. Sacchet, Toward a unified account of advanced concentrative absorption meditation: A systematic definition and classification of jhāna. Mindfulness 15, 1375–1394 (2024).

19. P. Dennison, The human default consciousness and its disruption: Insights from an EEG study of Buddhist jhana meditation. Frontiers in Human Neuroscience 13, 178–178 (2019).

20. M. R. Hagerty et al., Case study of ecstatic meditation: fMRI and EEG evidence of self-stimulating a reward system. Neural Plasticity 2013, 653572 (2013).

21. A. Chowdhury et al., Multimodal neurophenomenology of advanced concentration absorption meditation: An intensively sampled case study of Jhana. Neuroimage 305, 120973 (2024).

22. I. N. Treves, W. F. Z. Yang, T. Sparby, M. D. Sacchet, Dynamic brain states underlying advanced concentrative absorption meditation: A 7T fMRI intensive case study. Network Neuroscience, 1–55 (2024).

23. S. Ganesan, W. F. Z. Yang, A. Chowdhury, A. Zalesky, M. D. Sacchet, Within-subject reliability of brain networks during advanced meditation: An intensively sampled 7 Tesla MRI case study. Human Brain Mapping 45, e26666 (2024).

24. A. Chowdhury et al., Multimodal neurophenomenology of advanced concentration absorption meditation: An intensively sampled case study of Jhana. Neuroimage 305, 120973 (2025).

25. U. Demir, W. F. Z. Yang, M. D. Sacchet, Advanced concentrative absorption meditation reorganizes functional connectivity gradients of the brain: 7T MRI and phenomenology case study of jhana meditation. Cereb Cortex 35, (2025).

26. R. M. Potash, S. van Mil, M. Estarellas, A. Canales-Johnson, M. D. Sacchet, Integrated phenomenology and brain connectivity demonstrate changes in nonlinear processing in jhana advanced meditation. Journal of Cognitive Neuroscience, 1–24 (2025).

27. R. M. Potash et al., Investigating the complex cortical dynamics of an advanced concentrative absorption meditation called jhanas (ACAM-J): a geometric eigenmode analysis. Cereb Cortex 35, (2025).

28. G. A. Mashour, P. Roelfsema, J. P. Changeux, S. Dehaene, Conscious Processing and the Global Neuronal Workspace Hypothesis. Neuron 105, 776–798 (2020).

29. G. Tononi, M. Boly, M. Massimini, C. Koch, Integrated information theory: from consciousness to its physical substrate. Nat Rev Neurosci 17, 450–461 (2016).

30. A. K. Seth, T. Bayne, Theories of consciousness. Nat Rev Neurosci 23, 439–452 (2022).

31. H. Tal, M. Wright, S. Prest, L. Sandved-Smith, M. D. Sacchet. (Preprint), vol. 2025.

32. T. Metzinger, Minimal phenomenal experience. Philosophy and the Mind Sciences 1, 1–44 (2020).

33. T. Brandmeyer, A. Delorme, Reduced mind wandering in experienced meditators and associated EEG correlates. Experimental Brain Research 236, 2519–2528 (2018).

34. K. C. Fox et al., Functional neuroanatomy of meditation: A review and meta-analysis of 78 functional neuroimaging investigations. Neuroscience and Biobehavioral Reviews 65, 208–228 (2016).

35. M. D. Lieberman, M. A. Straccia, M. L. Meyer, M. Du, K. M. Tan, Social, self, (situational), and affective processes in medial prefrontal cortex (MPFC): Causal, multivariate, and reverse inference evidence. Neuroscience and Biobehavioral Reviews 99, 311–328 (2019).

36. J. J. F. Ribas-Fernandes, D. Shahnazian, C. B. Holroyd, M. M. Botvinick, Subgoal- and Goal-related Reward Prediction Errors in Medial Prefrontal Cortex. Journal of Cognitive Neuroscience 31, 8–23 (2019).

37. G. Desbordes et al., Moving beyond Mindfulness: Defining Equanimity as an Outcome Measure in Meditation and Contemplative Research. Mindfulness 6, 356–372 (2015).

38. K. Singh et al., Structural connectivity of autonomic, pain, limbic, and sensory brainstem nuclei in living humans based on 7 Tesla and 3 Tesla MRI. Hum Brain Mapp 43, 3086–3112 (2022).

39. M. Girn et al., Serotonergic psychedelic drugs LSD and psilocybin reduce the hierarchical differentiation of unimodal and transmodal cortex. Neuroimage 256, 119220 (2022).

40. C. Timmermann et al., Human brain effects of DMT assessed via EEG-fMRI. Proc Natl Acad Sci U S A 120, e2218949120 (2023).

41. Z. Josipovic, Neural correlates of nondual awareness in meditation. Ann N Y Acad Sci 1307, 9–18 (2014).

42. Z. Josipovic, V. Miskovic, Nondual Awareness and Minimal Phenomenal Experience. Front Psychol 11, 2087 (2020).

43. J. Vohryzek et al., N,N-dimethyltryptamine effects on connectome harmonics, subjective experience and comparative psychedelic experiences. Neuropsychopharmacology, (2025).

44. S. Atasoy et al., Connectome-harmonic decomposition of human brain activity reveals dynamical repertoire re-organization under LSD. Scientific Reports 7, 17661 (2017).

45. J. C. Pang et al., Geometric constraints on human brain function. Nature 618, 566–574 (2023).

46. S. Ehmann, I. Sezer, I. N. Treves, J. D. E. Gabrieli, M. D. Sacchet, Mindfulness, cognition, and long-term meditators: Toward a science of advanced meditation. Imaging Neuroscience, (2025).

47. D. Y. Panitz, A. Mendelsohn, J. Cabral, A. Berkovich-Ohana, Long-term mindfulness meditation increases occurrence of sensory and attention brain states. Front Hum Neurosci 18, 1482353 (2024).

48. A. Berkovich-Ohana, M. Harel, A. Hahamy, A. Arieli, R. Malach, Alterations in task-induced activity and resting-state fluctuations in visual and DMN areas revealed in long-term meditators. Neuroimage 135, 125–134 (2016).

49. A. Lutz, O. Abdoun, Y. Dor-Ziderman, F. M. Trautwein, A. Berkovich-Ohana, An Overview of Neurophenomenological Approaches to Meditation and Their Relevance to Clinical Research. Biol Psychiatry Cogn Neurosci Neuroimaging 10, 411–424 (2025).

50. A. Lutz, E. Thompson, Neurophenomenology: Integrating subjective experience and brain dynamics in the neuroscience of consciousness. Journal of consciousness studies 10, 31–52 (2003).

51. C. Petitmengin, M. van Beek, M. Bitbol, J. M. Nissou, A. Roepstorff, Studying the experience of meditation through micro-phenomenology. Current Opinion in Psychology 28, 54–59 (2019).

52. C. Timmermann et al., A neurophenomenological approach to non-ordinary states of consciousness: hypnosis, meditation, and psychedelics. Trends Cogn Sci 27, 139–159 (2023).

53. S. Dehaene, J. P. Changeux, Experimental and theoretical approaches to conscious processing. Neuron 70, 200–227 (2011).

54. M. T. Alkire, A. G. Hudetz, G. Tononi, Consciousness and anesthesia. Science 322, 876–880 (2008).

55. M. J. Redinbaugh, Y. B. Saalmann, Contributions of Basal Ganglia Circuits to Perception, Attention, and Consciousness. Journal of Cognitive Neuroscience 36, 1620–1642 (2024).

56. T. Sparby, Phenomenology and contemplative universals - The meditative experience of dhyana, Coalescence, or access concentration. Journal of Consciousness Studies 26, 130–156 (2019).

57. G. Tononi, An information integration theory of consciousness. BMC Neuroscience 5, 42–42 (2004).

58. G. Tononi, Consciousness as integrated information: a provisional manifesto. Biological Bulletin 215, 216–242 (2008).

59. M. Jung, K. M. Han, Behavioral Activation and Brain Network Changes in Depression. J Clin Neurol 20, 362–377 (2024).

60. X. Shan et al., Abnormal regional activity in the prefrontal-limbic circuit at rest: Potential imaging markers and treatment predictors in drug-naive anxiety disorders. CNS Neurosci Ther 30, e14523 (2024).

61. H. Kober et al., Brain Activity During Cocaine Craving and Gambling Urges: An fMRI Study. Neuropsychopharmacology 41, 628–637 (2016).

62. L. S. Paludetto, L. L. A. Florence, J. Torales, A. Ventriglio, J. M. Castaldelli-Maia, Mapping the Neural Substrates of Cocaine Craving: A Systematic Review. Brain Sci 14, (2024).

63. Y. Hadash, N. Segev, G. Tanay, P. Goldstein, A. Bernstein, The Decoupling Model of Equanimity: Theory, Measurement, and Test in a Mindfulness Intervention. Mindfulness 7, 1214–1226 (2016).

64. B. Anālayo, Relating Equanimity to Mindfulness. Mindfulness 12, 2635–2644 (2021).

65. P. Jijina, U. N. Biswas, Understanding equanimity from a psychological perspective: implications for holistic well-being during a global pandemic. Mental Health, Religion & Culture 24, 873–886 (2021).

66. M. Uebel, Equanimity in psychiatric medicine: the mind in the middle - Psychiatry in history. Br J Psychiatry 225, 413 (2024).

67. B. Lord, J. J. B. Allen, S. Young, J. L. Sanguinetti, Enhancing Equanimity With Noninvasive Brain Stimulation: A Novel Framework for Mindfulness Interventions. Biol Psychiatry Cogn Neurosci Neuroimaging 10, 384–392 (2025).

68. W. F. Z. Yang, A. Chowdhury, T. Sparby, M. D. Sacchet, Deconstructing the self and reshaping perceptions: An intensive whole-brain 7T MRI case study of the stages of insight during advanced investigative insight meditation. Neuroimage 305, 120968 (2025).

69. K. Abellaneda-Perez, R. M. Potash, A. Pascual-Leone, M. D. Sacchet, Neuromodulation and meditation: A review and synthesis toward promoting well-being and understanding consciousness and brain. Neurosci Biobehav Rev 166, 105862 (2024).

70. S. Ehmann, I. Sezer, A. S. Keller, I. N. Treves, M. D. Sacchet. (Preprint), vol. 2025.

71. K. Rayani et al., Brain stimulation enhances dispositional mindfulness in PTSD: an exploratory sham-controlled rTMS trial. Front Psychiatry 16, 1494567 (2025).

72. R. S. Wilson et al., Influence of epoch length on measurement of dynamic functional connectivity in wakefulness and behavioural validation in sleep. Neuroimage 112, 169–179 (2015).

73. Y. Y. Tang, M. K. Rothbart, M. I. Posner, Neural correlates of establishing, maintaining, and switching brain states. Trends Cogn Sci 16, 330–337 (2012).

74. W. F. Z. Yang et al., Intensive whole-brain 7T MRI case study of volitional control of brain activity in deep absorptive meditation states. Cerebral Cortex 34, (2024a).

75. R. A. Baer, J. Carmody, M. Hunsinger, Weekly change in mindfulness and perceived stress in a mindfulness-based stress reduction program. J Clin Psychol 68, 755–765 (2012).

76. G. Feldman, A. Hayes, S. Kumar, J. Greeson, J.-P. Laurenceau, Mindfulness and Emotion Regulation: The Development and Initial Validation of the Cognitive and Affective Mindfulness Scale-Revised (CAMS-R). Journal of Psychopathology and Behavioral Assessment 29, 177–190 (2006).

77. F. H. N. Chio, M. H. C. Lai, W. W. S. Mak, Development of the Nonattachment Scale-Short Form (NAS-SF) Using Item Response Theory. Mindfulness 9, 1299–1308 (2017).

78. H. T. Rogers, A. G. Shires, B. A. Cayoun, Development and Validation of the Equanimity Scale-16. Mindfulness 12, 107–120 (2020).

79. D. Derryberry, M. A. Reed, Anxiety-related attentional biases and their regulation by attentional control. J Abnorm Psychol 111, 225–236 (2002).

80. S. Singh et al., The TestMyBrain Digital Neuropsychology Toolkit: Development and Psychometric Characteristics. J Clin Exp Neuropsychol 43, 786–795 (2021).

81. R. W. Cox, AFNI: software for analysis and visualization of functional magnetic resonance neuroimages. Comput Biomed Res 29, 162–173 (1996).

82. K. J. Friston, Statistical parametric mapping: The analysis of functional brain images., (Academic Press., 2011).

83. Y. Zang, T. Jiang, Y. Lu, Y. He, L. Tian, Regional homogeneity approach to fMRI data analysis. Neuroimage 22, 394–400 (2004).

84. A. Schaefer et al., Local-Global Parcellation of the Human Cerebral Cortex from Intrinsic Functional Connectivity MRI. Cerebral Cortex 28, 3095–3114 (2018).

85. Y. Tian, D. S. Margulies, M. Breakspear, A. Zalesky, Topographic organization of the human subcortex unveiled with functional connectivity gradients. Nature Neuroscience 23, 1421–1432 (2020).

86. M. Bianciardi et al., In vivo functional connectome of human brainstem nuclei of the ascending arousal, autonomic, and motor systems by high spatial resolution 7-Tesla fMRI. MAGMA 29, 451–462 (2016).

87. M. King, C. R. Hernandez-Castillo, R. A. Poldrack, R. B. Ivry, J. Diedrichsen, Functional boundaries in the human cerebellum revealed by a multi-domain task battery. Nature Neuroscience 22, 1371–1378 (2019).

88. R. Vos de Wael et al., BrainSpace: a toolbox for the analysis of macroscale gradients in neuroimaging and connectomics datasets. Commun Biol 3, 103 (2020).

89. M. F. Glasser et al., The minimal preprocessing pipelines for the Human Connectome Project. Neuroimage 80, 105–124 (2013).

90. S. Atasoy, I. Donnelly, J. Pearson, Human brain networks function in connectome-specific harmonic waves. Nature Communications 7, 10340 (2016).

91. J. Y. Hansen et al., Mapping neurotransmitter systems to the structural and functional organization of the human neocortex. Nat Neurosci 25, 1569–1581 (2022).

92. A. I. Luppi et al., In vivo mapping of pharmacologically induced functional reorganization onto the human brain’s neurotransmitter landscape. Science Advances 9, eadf8332 (2023).

93. T. Yarkoni, R. A. Poldrack, T. E. Nichols, D. C. Van Essen, T. D. Wager, Large-scale automated synthesis of human functional neuroimaging data. Nature Methods 8, 665–670 (2011).

94. S. van Buuren, K. Groothuis-Oudshoorn, mice: Multivariate imputation by chained equations in R. Journal of Statistical Software 45, 1–67 (2011).

95. I. R. White, P. Royston, A. M. Wood, Multiple imputation using chained equations: Issues and guidance for practice. Stat Med 30, 377–399 (2011).

96. R Core Team. (R Foundation for Statistical Computing, Vienna, Austria, 2022).

97. Posit team. (Posit Software, PBC http://www.posit.co, Boston, MA, 2022).

98. A. Krishnan, L. J. Williams, A. R. McIntosh, H. Abdi, Partial Least Squares (PLS) methods for neuroimaging: a tutorial and review. Neuroimage 56, 455–475 (2011).

99. B. Vazquez-Rodriguez et al., Gradients of structure-function tethering across neocortex. Proc Natl Acad Sci U S A 116, 21219–21227 (2019).

100. A. F. Alexander-Bloch et al., On testing for spatial correspondence between maps of human brain structure and function. Neuroimage 178, 540–551 (2018).

101. R. D. Markello, B. Misic, Comparing spatial null models for brain maps. Neuroimage 236, 118052 (2021).

102. F. Vasa, B. Misic, Null models in network neuroscience. Nat Rev Neurosci 23, 493–504 (2022).

103. T. Salo et al., NiMARE: neuroimaging meta-analysis research environment. Aperture Neuro 3, 1–32 (2023).

## References

1. T. Sparby, M. D. Sacchet, Toward a unified account of advanced concentrative absorption meditation: A systematic definition and classification of jhāna. Mindfulness 15, 1375–1394 (2024).

2. B. Anālayo, A brief history of Buddhist absorption. Mindfulness 11, 571–586 (2019).

3. L. Brasington, Right concentration: A practical guide to the Jhanas. (Shambala Publications, Boston, MA, 2015).

4. S. Snyder, T. Rasmussen, Practicing the Jhanas: Traditional concentration meditation as presented by the Venerable Pa Auk Sayadaw. (Shambala Publications, Boston, MA, 2009).

5. N. E. F. Quli, Multiple buddhist modernisms: Jhāna in convert Theravāda. Pacific World 10, 225–249 (2008).

6. R. E. Buswell et al., The Princeton Dictionary of Buddhism. (Princeton University Press, Princeton, NJ, 2014).

7. R. Shankman, The Experience of Samadhi: An In-depth Exploration of Buddhist Meditation. (Shambhala Publications, Inc., Boston, MA., 2008).

8. A. Brahm, Mindfulness, Bliss, and Beyond: A Meditator’s Handbook. (Wisdom Publications, Inc, MA, MA, 2005).

9. T.-f. Kuan, Cognitive operations in Buddhist meditation: interface with Western psychology. Contemporary Buddhism 13, 35–60 (2012).

